# skNAC is a Key Driver of Cardiomyocyte Integrity Against Pathological Cardiac Hypertrophy and Heart Failure

**DOI:** 10.1101/2025.10.23.684272

**Authors:** Laura Guilbert, Justine Dontaine, Natacha Fourny, Ettore Vanni, Michele Russo, Hugo Vanderroost, Jade Dron, Jérôme Ambroise, Hrag Esfahani, Caroline Bouzin, Younes Achouri, Emilie Hendrickx, Nassiba Menghoum, B. Bearzatto, D. Vertommen, Laure Dumoutier, Andreas Unger, Wolfgang A. Linke, Laurent Bultot, Alice Marino, Sandrine Horman, Christophe Beauloye, Luc Bertrand

## Abstract

Chronic pressure overload induces cardiac hypertrophy and heart failure through coordinated alterations in proteome homeostasis, metabolism and sarcomere organisation. The muscle-specific α-isoform of the nascent polypeptide-associated complex (skNAC) is essential for sarcomere assembly during development, but its role in adult hearts remains largely unknown. Here, we show that skNAC expression is reduced in hypertrophic cardiomyocytes, mouse models of pressure overload, and human hypertrophic hearts, in association with disease severity. Cardiomyocyte-specific skNAC deletion results in basal hypertrophy, systolic dysfunction, and premature death, and exacerbates pressure overload-induced heart failure. At the molecular level, skNAC associates with ribosomes and is required for sarcomere organisation maintenance, while its loss induces autophagy and ultrastructural defects. Integrated transcriptomic and proteomic analyses reveal early downregulation of metabolic gene expression despite increased abundance of corresponding proteins, indicating compensatory metabolic responses. Gain-of-function studies confirm a protective role against hypertrophy. Together, these data establish skNAC as a key regulator of cardiac proteome homeostasis and metabolic adaptation during pathological remodelling.

## Introduction

Heart failure (HF) is a complex, progressive syndrome that arises as the endpoint of diverse cardiac stressors, including chronic pressure overload, ischemic injury, and inherited cardiomyopathies ^1,2^. Under pressure overload, transition to HF is driven by maladaptive cardiac remodelling, a process characterized by cardiomyocyte hypertrophy, reactivation of foetal gene programs, interstitial fibrosis, and progressive contractile dysfunction ^3^. HF remains a major clinical challenge, highlighting the need to identify novel therapeutic targets. The heterodimeric nascent polypeptide–associated complex (NAC), composed of αNAC and βNAC subunits, is a highly abundant ribosome-bound chaperone that binds emerging polypeptides and prevents inappropriate signal recognition particle engagement and mistargeting to the endoplasmic reticulum ^4,5^. Beyond its canonical role, αNAC has been reported to modulate transcription in specific contexts ^6–8^. In addition to αNAC, which is ubiquitously expressed and composed of 215 amino acids, there is a muscle-specific variant of NAC termed skeletal muscle-specific isoform NAC (skNAC) that is generated by the insertion of a large alternative exon (exon 3, ∼6 kb), producing a high molecular weight protein of 2,187 residues ^9,10^. Splicing toward skNAC has been shown to be regulated by muscle-enriched RNA-binding proteins, predominantly Rbm24, a major driver of muscle-specific exon inclusion ^11^. Rbm24 frequently cooperates with Rbm20, a key cardiac splicing factor best known for controlling titin isoforms ^12,13^. Developmental studies established skNAC as essential for striated muscle formation. In skeletal muscle, skNAC rises during myogenesis and supports myoblast fusion, regeneration, and gene programs linked to differentiation ^9,14,15^. skNAC also partners with the histone methyltransferase Smyd1, coupling transcriptional control to myofibril assembly ^16^. *In vivo* genetic evidence (zebrafish and mouse deficient models) shows that skNAC is required for sarcomeric organisation and stabilisation of contractile proteins, with deficiency causing defective myofibrillogenesis and cardiac structural abnormalities ^15–17^. Of note, skNAC has been previously reported to undergo O-GlcNAcylation, a post-translational modification that has been shown to participate in the development of cardiac hypertrophy and HF ^18,19^. Together, these data position skNAC as a putative integrator of RNA splicing, co-translational protein quality control, and sarcomere integrity. To date, one single and recent study has shown that partial cardiac deletion of skNAC leads to cardiac dysfunction associated with mitochondrial protein downregulation and impaired energy metabolism ^20^. However, its precise role in the adult heart, under both physiological conditions and pathological remodelling, remains largely unclear.

Here, we test the hypothesis that skNAC acts as a stress-contingent regulator of cardiomyocyte structure and function in the healthy, hypertrophic and failing adult heart. Using complementary *in vitro* and *in vivo* models, we aim to: (i) define the expression of skNAC under basal, hypertrophic and HF conditions, investigating its regulation through Rbm24 and Rbm20 proteins in cardiomyocyte; (ii) establish the structural and functional consequences of skNAC deficiency in the adult heart under basal and pressure overload-induced hypertrophic and HF conditions; (iii) determine whether skNAC overexpression protects against pro-hypertrophic stimuli and, (iv) characterise skNAC’s subcellular localization, ribosomal interactions and its effects on mRNA and protein expression through proteomic and transcriptomic analyses. This integrative approach identifies skNAC as a critical determinant of cardiomyocyte proteome and structural homeostasis and suggests that modulation of skNAC could provide new opportunities for therapeutic intervention in HF.

## Results

### skNAC expression is reduced in hypertrophic mouse and human hearts

skNAC and αNAC isoforms arise from alternative splicing of the single *Naca* gene (**Figure 1a**). Besides the commercially available αNAC antibody, we first developed a polyclonal antibody specific to exon 3 to monitor skNAC protein. Western blot analysis of adult mouse tissues confirmed that skNAC exhibits tissue specificity, being exclusively expressed in cardiac and skeletal muscles, in contrast to αNAC, which is ubiquitously expressed (**Figure 1a**). To precisely determine skNAC expression in adult mouse, we performed an absolute quantification by RT-qPCR. Our results indicated comparable levels of skNAC expression across the heart and skeletal muscles (gastrocnemius and quadriceps), with no detectable expression in the liver, confirming a muscle-specific profile (**Figure 1b**). We next sought to compare skNAC and αNAC gene expression in these tissues. In skeletal muscle tissues, such as the gastrocnemius (**Figure 1c**) and quadriceps (**Figure 1d**), αNAC expression was three-fold higher than skNAC, while only αNAC was detectable in the liver (**Figure 1e**). In the heart, αNAC expression was approximately twelve-fold higher than that of skNAC (**Figure 1f**). Of note, an αNAC/skNAC ratio of 0.5 was observed in neonatal rat ventricular myocytes (NRVMs) (**Figure 1g**), confirming higher expression of skNAC in embryonic and neonatal stages ^16^. We then evaluated whether skNAC and αNAC expression were affected in models of cardiac hypertrophy. *In vitro*, the frequently used model of NRVMs treated for 24 hours with pro-hypertrophic agents, such as phenylephrine (PE) or angiotensin II (Ang II), was chosen. We previously showed that such treatment provokes NRVMs hypertrophy ^21^. Both PE- and Ang II-treated NRVMs exhibited a significant reduction in skNAC expression, while αNAC levels were unchanged (**Figure 1h,i).** These results were corroborated *in vivo* using two murine models of cardiac hypertrophy that we previously characterised, transverse aortic constriction (TAC), and chronic Ang II infusion ^21^. skNAC mRNA level was significantly reduced in both TAC and Ang II models, whereas αNAC expression remained stable (**Figure 1j,k**). For confirmation purposes, we analysed publicly available RNA-seq datasets ^22^ from hypertrophic mouse hearts at various time points following TAC surgery, confirming that skNAC expression progressively declined during hypertrophic progression, whereas αNAC levels were unchanged or slightly increased (**Figure S1**). From a translational perspective, we explored human cardiac biopsies from patients previously validated in our clinical institution ^23^. Those patients suffered from severe aortic stenosis leading to cardiac hypertrophy (**Table S1**). Gene expression evaluation revealed an inverse correlation between skNAC mRNA level and left ventricular mass index (LVMi, echocardiography), indicating that the most hypertrophic human hearts had lower skNAC expression (**Figure 1l**). A similar negative correlation can be found between skNAC expression and circulating NT-proBNP level, a reliable biomarker of cardiac dysfunction (**Figure 1m**). This demonstrates that human hypertrophied hearts have lower skNAC expression.

**Figure 1:**
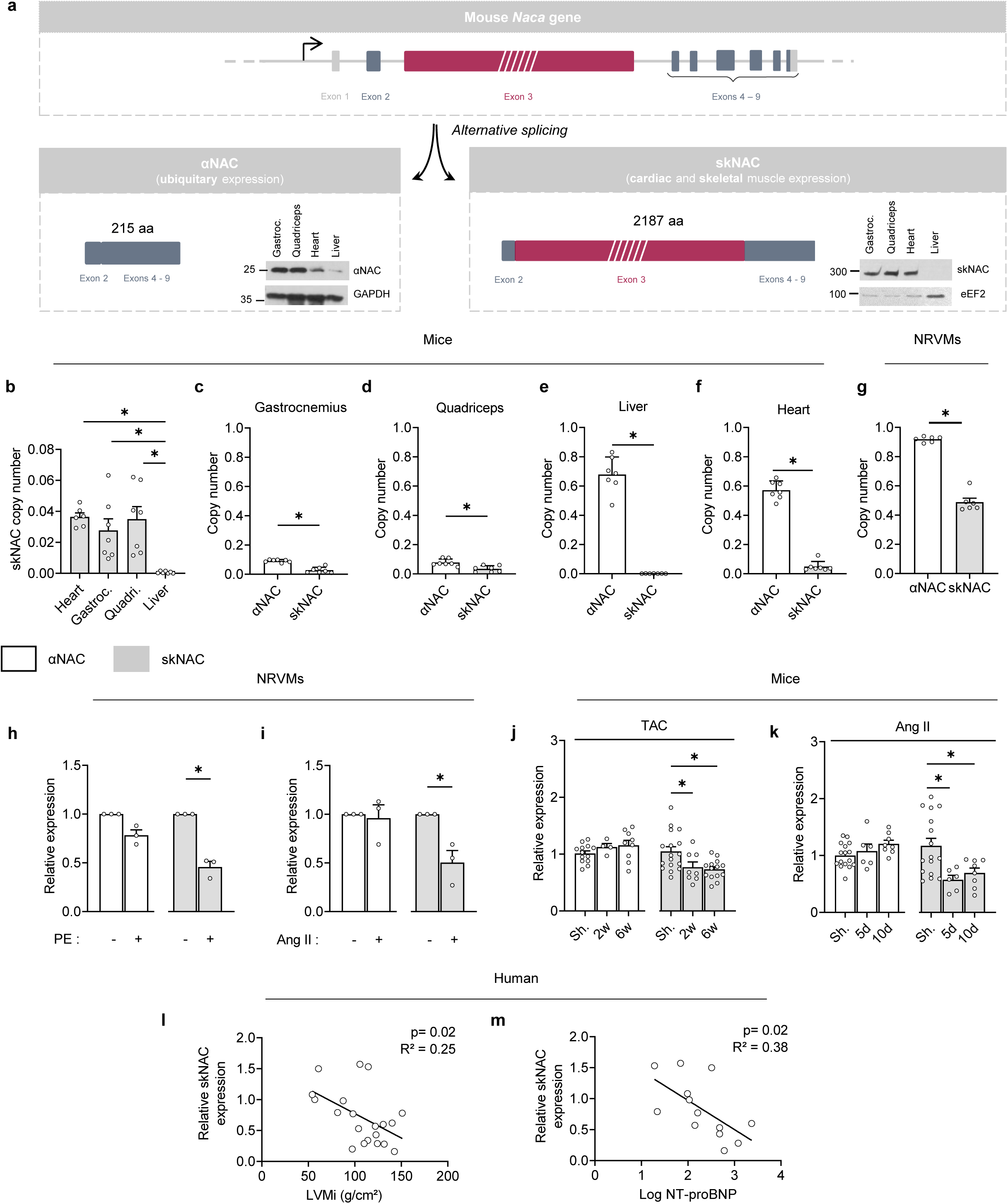
skNAC expression is reduced in hypertrophic and failing mouse heart and in human cohort. **a,** Schematic representation of mouse *Naca* gene. Alternative splicing results in ubiquitously expressed αNAC and muscle-specific skNAC isoforms. **b,** skNAC mRNA copy number in heart, gastrocnemius, quadriceps, and liver of adult mice (N=7). **c-f,** αNAC and skNAC mRNA copy number in adult mouse gastrocnemius **(c)**, quadriceps **(d)**, liver **(e)** and cardiac **(f)** tissues (N=7). **g,** αNAC and skNAC mRNA copy number in NRVMs (N=6). **h-i,** αNAC and skNAC mRNA level in NRVMs treated or not with phenylephrine (PE, 20µM) **(H)** or with Angiotensin II (Ang II, 100 nM) for 24h **(i)** (N=3). **j-k,** αNAC and skNAC cardiac mRNA level of mouse submitted (TAC) or not (Sham, Sh.) to transverse aortic constriction for 2 or 6 weeks (w) (N=4-17) **(j)** or treated with Ang II (2mg/kg/d) for 5 or 14 days (d) (N=6-16) **(k)**. **l-m,** Correlation of skNAC mRNA level in human cardiac biopsies with left ventricular mass index (LVMi, g/cm^2^) **(l)** or plasma NT-pro BNP level **(m)**. All mRNA copy numbers or relative values are normalised to *Rpl32.* Each point represents an independent biological replicate, animal or patient. Values are expressed as mean ± SEM. Statistical analyses are performed using unpaired student t-test **(c-i)**, one-way ANOVA followed Bonferroni post hoc test **(b),** two-way ANOVA followed Bonferroni **(j, k)** or linear regression analysis **(l, m)**. \**p<0.05* vs. corresponding untreated cells or sham (Sh.) mice.

Lastly, we searched for *NACA* gene mutations potentially linked to cardiac diseases. The analysis of the NHGRI-EBI GWAS Catalog revealed that SNPs at the *NACA* locus are predominantly associated with cardiovascular traits (**Figure S2a,b**). Among these, cardiac conduction abnormalities, particularly QRS duration, were the most frequent associations (**Figure S2c**) and mostly mapped on the exon 3 specifically spliced in the skNAC isoform (**Figure S2d**). Altogether, these results demonstrate that skNAC expression is strongly correlating to the development of cardiac complications and, most importantly, is downregulated during cardiac hypertrophy and remodelling, suggesting that this protein plays an anti-hypertrophic role.

### Cardiac skNAC expression is regulated by RNA binding proteins during hypertrophy development

skNAC expression results from the alternative splicing of *NACA* gene. Because cardiac hypertrophy induces a specific reduction of skNAC but not of αNAC expression, we reasonably hypothesized that the hypertrophic-mediated decrease of skNAC results from a regulation of pre-mRNA splicing rather than a change in pre-mRNA levels. Consequently, we investigated key regulators known to be involved in similar processes, namely the RNA-binding motif (Rbm) proteins. Rbm24 is a well-characterised RNA-binding protein that regulates the alternative splicing of numerus skeletal and cardiac specific genes, including *NACA* gene ^24^. Rbm24 has been shown to specifically promote the inclusion of exon 3 to produce skNAC isoform in skeletal and cardiac muscle cells ^11^. In line with the reduction of skNAC, Rbm24 mRNA levels were significantly reduced after 24 hours of PE treatment in comparison to untreated NRVMs (**Figure 2a**). A similar reduction of Rbm24 expression can be found *in* hypertrophic TAC treatment (**Figure 2b**). The functional implication of Rbm24 was further supported by siRNA-mediated knockdown experiments in NRVMs (**Figure 2c,d**). An approximatively 50% efficacy of siRNA-mediated reduction of Rbm24 expression was sufficient to significantly reduce skNAC expression whereas αNAC was slightly and non-significantly increased.

**Figure 2:**
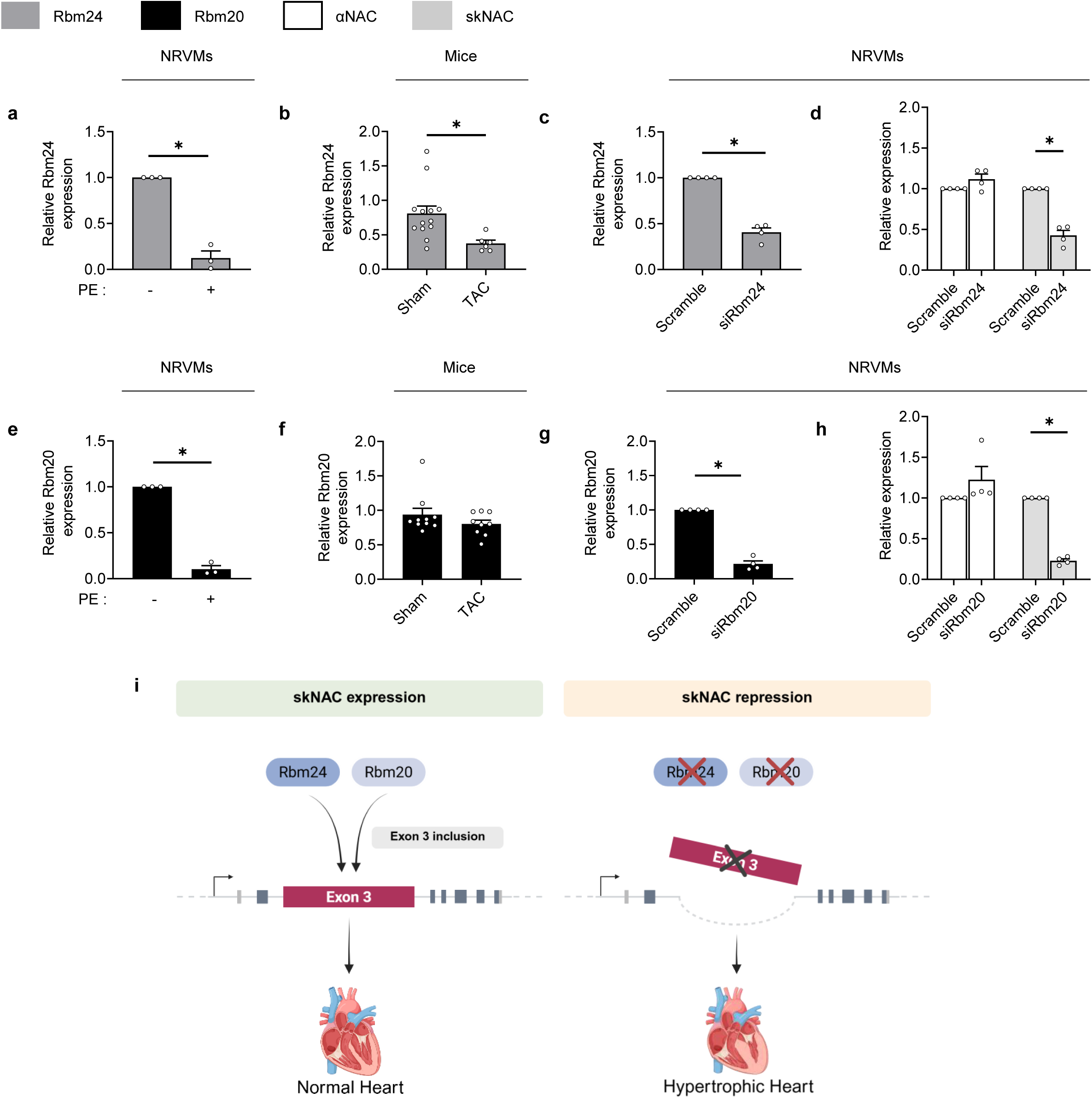
skNAC expression is under the control of RNA binding proteins Rbm24 and Rbm20 in cardiomyocytes. **a-b,** Rbm24 mRNA level in NRVMs treated or not with phenylephrine (PE, 20µM) for 24h (N=3) **(a)**, or in heart of mice submitted (TAC) or not (Sham) to transverse aortic constriction for 6 weeks (w) (N=6-12) **(b)**. **c-d,** Rbm24 **(c)**, αNAC and skNAC **(d)** mRNA level in NRVMs treated with control non-targeting siRNA (Scramble) or Rbm24 siRNA (siRbm24) for 72h (N=4). **e-f,** Rbm20 mRNA level in NRVMs treated or not with phenylephrine (PE, 20µM) for 24h (N=3) **(e)**, or in heart of mice submitted (TAC) or not (Sham) to transverse aortic constriction for 6 weeks (N=6-12) **(f)**. **g-h,** Rbm20 **(g)**, αNAC and skNAC **(h)** mRNA level in NRVMs treated with control non-targeting siRNA (Scramble) or Rbm20 siRNA (siRbm20) for 72h (N=4). All mRNA relative values are normalised to *Rpl32.* Each point represents an independent biological replicate or animal. Data are presented as mean ± SEM. Statistical analyses are performed using unpaired student t-test **(a, b, c, e, f, g)** or using two-way ANOVA followed Bonferroni post hoc test **(d, h)**. \**p<0.05* vs. corresponding untreated cells or sham mice. **i,** Schematic representation of regulation of *NACA* pre-mRNA splicing by Rbm24 and Rbm20. In healthy heart, Rbm24 and Rbm20 induce exon 3 inclusion for skNAC expression. Rbm24/20 reduction promotes exon 3 splicing, leading to skNAC repression, which participates in cardiomyocyte hypertrophy. Original Figure 2i created with a licensed version of Biorender.com.

We next investigated Rbm20, another Rbm family member, which is implicated in inherited cardiomyopathies ^12,13^. Rbm20 was recently found to cooperate with Rbm24 for the maturation of complex cardiac pre-mRNAs, such as *TTN*, the gene encoding titin protein ^25–28^. Similar to Rbm24, Rbm20 mRNA expression was markedly reduced in PE-treated NRVMs (**Figure 2e**). Whereas Rbm20 expression was lowered under such acute *in vitro* settings, it looked unchanged six weeks after TAC intervention (**Figure 2f**). Although further studies are required to elucidate the complex regulation of Rbm20 during hypertrophy, siRNA-induced knockdown of Rbm20 recapitulated the effect of Rbm24 depletion, leading to reduced skNAC with slightly increased αNAC levels (**Figure 2g,h**). Together, these findings identify Rbm24 and Rbm20 as critical regulators of skNAC expression through exon 3 inclusion (**Figure 2i**). The systemic downregulation of Rbm24, observed both *in vitro* and *in vivo*, provides a mechanistic explanation for the loss of skNAC expression during pressure overload-mediated cardiac remodelling.

### skNAC overexpression prevents NRVM hypertrophy development, whereas its silencing promotes NRVM hypertrophy

Given that skNAC expression is constantly downregulated in all investigated hypertrophic models to date, we first investigated the possible effect of its overexpression and siRNA-mediated knockdown in NRVMs under control and hypertrophic conditions. To address the consequences of skNAC overexpression, we utilized a pre-existing rodent skNAC cDNA (gift from Prof. B. Munz, University of Tübingen, Germany) and built different skNAC expressing vectors (**Figure S3**). Unfortunately, the excessive length of the full-length skNAC cDNA (which far exceeds the 4.7 kb packaging capacity of AAV9 when combined with necessary regulatory elements) precludes the use of this vector, despite AAV9 being considered the gold standard for cardiac targeting *in vivo*. So, we first generated non-viral constructs with a hemagglutinin (HA) tag either at the N-terminus (HA-skNAC) or the C-terminus (skNAC-HA) of skNAC, not knowing the native 3-D structure of skNAC (NAC family members are characterized by known extended intrinsically disordered regions, preventing the resolution of their full 3-D structure ^29^ and, therefore, the possible consequences of adding the tag on the stability of the expressed protein. skNAC overexpression was confirmed using anti-HA immunofluorescence staining with both HA-skNAC and skNAC-HA constructs, with a transfection efficiency similar to the GFP-expressing control plasmid (**Figure 3a,b and S4**). Both HA-skNAC and skNAC-HA overexpression did not affect cardiomyocyte size under unstimulated conditions but prevented PE-induced cardiomyocyte hypertrophy, compared to control cells transfected with GFP-expressing plasmid (**Figure 3a,b**). To discard the unlikely possibility that this may be the result of an artefact caused by the presence of the HA tag, we completed the construction of a vector allowing the co-expression of untagged skNAC and GFP (to be able to select transfected cells) (**Figure 3c,d**). This vector fully reproduced the data obtained with tagged skNAC constructs, skNAC overexpression preventing PE-induced cardiomyocyte hypertrophy.

**Figure 3:**
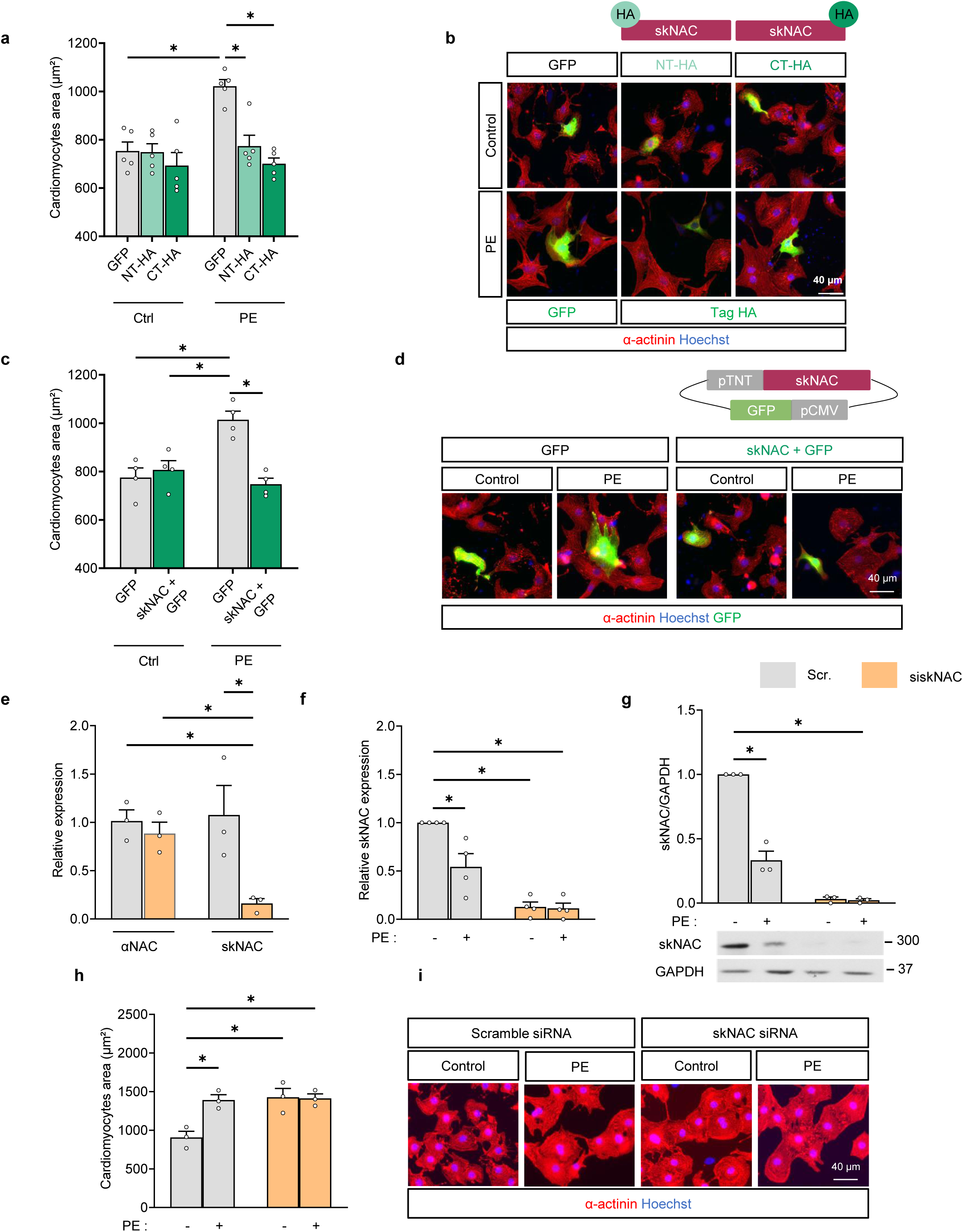
skNAC overexpression prevents cardiomyocyte hypertrophy while its deletion leads to basal hypertrophy. **a-b,** Effect of N-terminal HA-tagged skNAC (NT-HA) and C-terminal HA-tagged skNAC (CT-HA) overexpression on phenylephrine (PE, 20 µM)-mediated NRVMs hypertrophy. **c-d,** Effect of skNAC+GFP overexpression on PE-mediated NRVMs hypertrophy. Cells expressing GFP are used as control **(a-d)**. **e-i,** Effect of skNAC knockdown on PE-mediated NRVMs hypertrophy. αNAC and skNAC mRNA level in NRVMs treated with control non-targeting siRNA (Scramble, Scr.) or skNAC siRNA (siskNAC) for 72h (N=3) **(e)**. skNAC mRNA level **(f)**, skNAC protein level **(g)**, and cell size **(h, i)** in NRVMs treated with Scramble siRNA or siskNAC for 72h and treated or not with PE for 24h (N=3). mRNA levels are normalised to *Rpl32,* protein levels are normalised to GAPDH. Quantification **(a, c, h)** and representative images **(b, d, i)** of NRVM area (µm²) are analysed after α-actinin immunostaining. Scale bar, 40µm. Each point represents an independent biological replicate, with an average of 100–200 cells analysed per condition for each replicate. Data are represented ± SEM. Statistical analyses are performed using two-way ANOVA followed Bonferroni post hoc test. \**p<0.05* vs. corresponding control cells.

To further define the potential role of skNAC in protecting against hypertrophy, we next assessed whether reducing skNAC expression could exert the opposite effect and promote cardiomyocyte hypertrophy. For that purpose, we designed siRNA specifically targeting skNAC in exon 3. As shown in **Figure 3e**, siRNA transfection in NRVMs resulted in a significant 85% decrease in skNAC mRNA levels without affecting αNAC expression. Downregulation of skNAC was confirmed at mRNA levels (**Figure 3e,f**) but also at protein levels using the anti-skNAC antibody we developed, in both Western blot (**Figure 3g**) and immunofluorescence staining (**Figure S5**), validating the specificity of our home-made antibody. To determine the functional consequences of skNAC knockdown, we measured cardiomyocyte size under basal and PE treatment conditions. In line with our previous data, PE treatment induced the reduction of skNAC at both mRNA and proteins levels in siRNA control cells (**Figure 3f,g**). Remarkably, silencing skNAC led to an increase in NRVMs surface area under unstimulated conditions (**Figure 3h,i**) reaching levels comparable to those observed in PE-stimulated control cells, indicating that loss of skNAC alone is sufficient to trigger hypertrophy. Of note, PE could not further increase cardiomyocyte hypertrophy in absence of skNAC. Intriguingly, the expression of conventional foetal hypertrophic markers such as ANP and β-MHC was unaffected by skNAC silencing, while responding correctly to PE treatment (**Figure S6a**). The same applies for central mediators of cardiac hypertrophy, namely MAPK and protein translation signalling, represented by ERK1/2 or S6 phosphorylation, respectively (**Figure S6b,c**).

Taken together, these results identify skNAC as a central regulator of cardiomyocyte growth, acting as a molecular brake that prevents cardiomyocyte hypertrophy, by a yet undefined mechanism.

### Constitutive cardiomyocyte deletion of skNAC in mice induced hypertrophic remodelling and heart failure

The next step was to evaluate the consequences of skNAC deletion *in vivo*. We first generated transgenic mice by introducing two LoxP sites surrounding exon 3 of *Naca* gene by CRISPR/Cas9 gene-editing technology (**Figure S7a**). Resulting skNAC^flx/flx^ mice were crossed with *αMyHC-*Cre^+/−^ transgenic mice, which express Cre recombinase under cardiac-specific alpha myosin-heavy chain promoter to generate a constitutive and cardiac-specific knockout (cKO) mice. Deletion of cardiac skNAC was confirmed at the mRNA and protein level whereas αNAC was increased, most likely to compensate for the absence of the long isoform (**Figure S7b-e**). skNAC was normally expressed in skeletal muscle tissues (**Figure S7f**). Adult cKO mice were monitored longitudinally between the ages of two and ten months under baseline conditions. It has been previously demonstrated that *αMyHC-*Cre^+/−^ transgenic male mice exhibit decreased cardiac function by six months due to prolonged Cre expression ^30,31^. In consequence, we decided to firstly investigate cKO phenotype in females. Most notably, skNAC deletion negatively affected mice survival with a survival rate of less than 10% at ten months (**Figure 4a**). This was associated with an increased ventricular mass, a progressive left ventricular dilation (increased end-diastolic volume, EDV), and severe systolic dysfunction, reaching an ejection fraction of less than 20% for surviving animals (**Figure 4b-d, Table S2**). Consistent with previous reports ^30,31^, the male control littermates (cWT) exhibited progressive cardiac dysfunction and mortality. Similarly to the female animals, male cKO mice displayed a significant increase in mortality (**Figure 4e)** and more pronounced cardiac dysfunction than the cWT mice (**Figure 4f-h, Table S3**). To investigate structural abnormalities between cKO and control mice, we performed transmission electron microscopy analysis in six-month aged female mouse hearts. Loss of skNAC resulted in profound myofibrillar disorganization, extensive inter-sarcomeric vesiculation, and marked accumulation of numerous autophagic vacuoles, whereas control cardiomyocytes displayed intact and well-aligned sarcomeres (**Figure 4i**). The accumulation of autophagosomes was confirmed by an elevated LC3B-II/I ratio, the increase in p62 levels suggesting an impairment of autophagic flux (**Figure 4j-k**).

**Figure 4:**
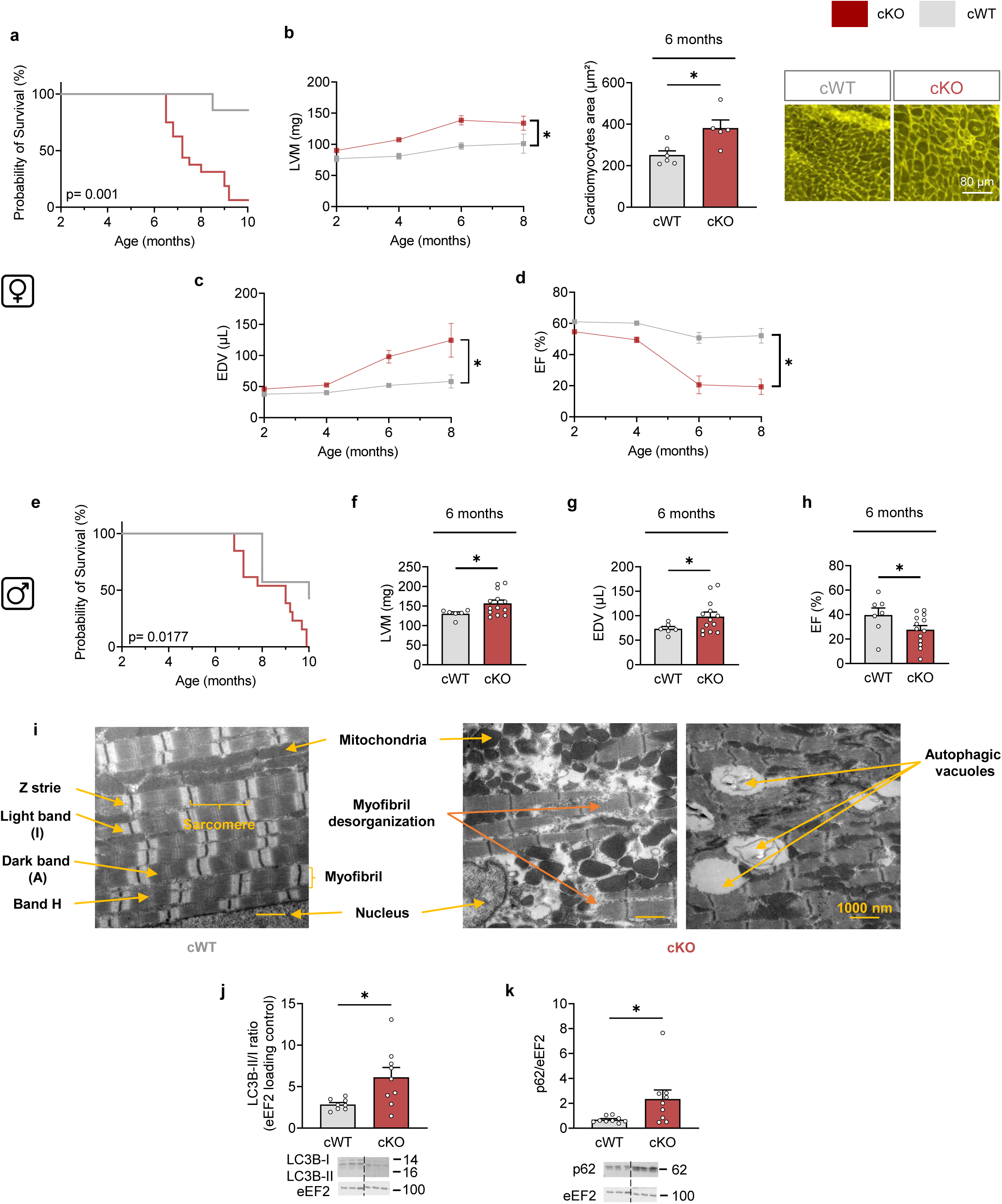
Constitutive cardiomyocyte-specific deletion of skNAC induced hypertrophic remodelling and heart failure in mice. **a,** Kaplan–Meier survival curves of cardiomyocyte-specific skNAC knockout (cKO) and control littermate (cWT) female mice (N=7–16). **b-d,** Longitudinal echocardiographic evaluation of left ventricular mass (LVM) **(b)**, end-diastolic volume (EDV) **(c)** and ejection fraction (EF) **(d)** (N=7-16). **e,** Kaplan–Meier survival curves of cWT and cKO male mice (N=7–13). **f-h,** Longitudinal echocardiographic evaluation of LVM **(f)**, EDV **(g)** and EF **(h)** (N=7-13). **i,** Electron microscopy analysis of left ventricular tissue from cWT and cKO mice at 6 months. cWT hearts display normal myofibril organisation, whereas cKO hearts exhibit myofibrillar disarray and autophagic vacuole accumulation (N=2–3). **j-k**, Immunoblots and quantification of LC3B-II/I ratio **(j)** and p62 **(k)** protein levels, normalised to eEF2 (N=8-9). Each point represents an independent animal. Values are expressed as mean ± SEM. Statistical analyses are performed using a two-way ANOVA followed Bonferroni post hoc test. *p<0.05 vs. cWT mice.

All together, these findings support a role for skNAC in maintaining myofibrillar organization and cell integrity, its loss compromising the animal lifespan through the onset of pathological cardiac hypertrophy, remodelling and systolic dysfunction.

### Constitutive skNAC cardiac deletion provokes rapid and severe heart failure under pressure overload condition

We, then, evaluated the pivotal role of skNAC in response to pressure overload. We subjected two-month-old male (**Figure 5**) and female (**Figure S8**) cKO mice to TAC surgery. We deliberately chose two-month-old animals, as this is before any signs of hypertrophic phenotype and cardiac dysfunction appear (see **Figure 4**). Blood flow velocities were measured to confirm a consistent aortic constriction between groups (**Figure 5a**). cKO exhibited an accelerated progression of HF following TAC (**Figure 5b, Table S4**), with substantial increase in EDV (**Figure 5c,d)**, coupled to decline in ejection fraction (**Figure 5e,f**) and fractional shortening (**Figure 5g,h**) within three weeks. Values of ejection fraction and fractional shortening fell below 20% and 10%, respectively, prompting us to terminate the follow-up at three weeks in all cohorts. This was associated with a more severe increase in left ventricular mass measured by echocardiography (**Figure 5i,j**) and heart weight/tibia length ratio after sacrifice (**Figure 5k**). Histological analyses confirmed exaggerated cardiomyocyte hypertrophy, with increased cell cross-sectional area (**Figure 5l,m**) and enhanced interstitial fibrosis (**Figure 5n,o**). Remarkably, as observed in the *in vitro* skNAC knockdown experiments, foetal gene reprogramming, including ANP, BNP and β-MHC, was promoted by TAC but showed no significant difference between cKO and control littermates (**Figure 5p-r**), supporting the concept that skNAC deficiency promotes cardiac remodelling through non-canonical mechanisms. Two-month-old female cKO mice challenged by TAC surgery showed a similar phenotype with an accelerated systolic dysfunction (**Figure S8a-h, Table S5**), more important left ventricular mass (**Figure S8i-m**), more pronounced increase in interstitial fibrosis (**Figure S8n,o**), and similar transcript levels of canonical hypertrophic markers (ANP, BNP, β-MHC) (**Figure S8p-r**).

**Figure 5:**
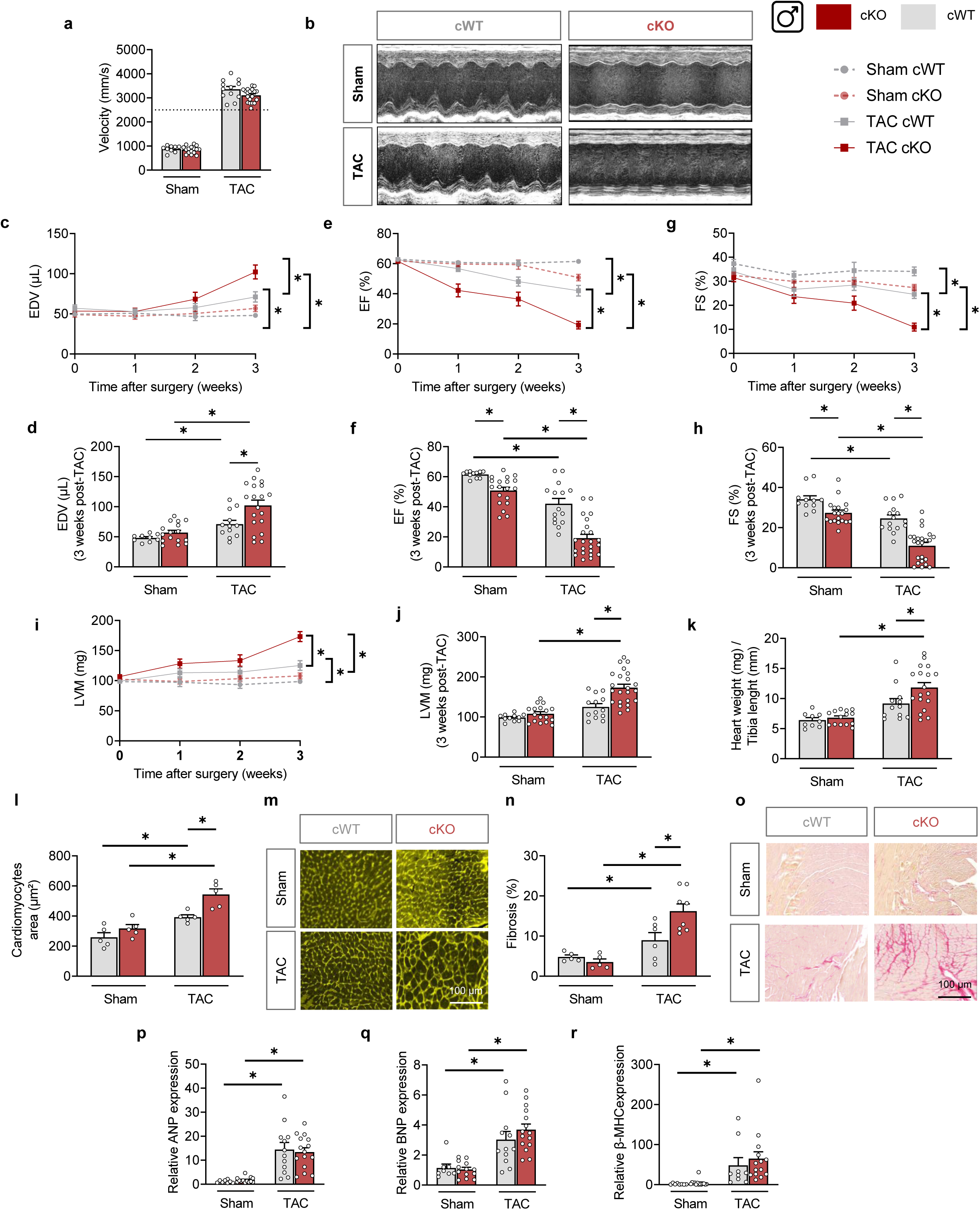
Constitutive cardiomyocyte-specific deletion of skNAC elicits rapid and severe heart failure under pressure overload condition. **a-j,** Echocardiographic analysis of sham- and transverse aortic constriction (TAC)-operated cardiomyocyte-specific skNAC knockout (cKO) and control littermate (cWT) mice at 1-, 2- and 3-weeks post-surgery (N=9-19). Quantification of peak aortic velocity at the site of constriction in sham- and TAC-operated cWT and cKO mice **(a)**. Representative image of M-mode in sham- and TAC-operated cWT and cKO mice at 3 weeks post-surgery **(b)**. End-diastolic volume (EDV) **(c, d)**, ejection fraction (EF) **(e, f)** and fractional shortening (FS) **(g, h)** evaluated by echocardiography over time and at end-time point (3 weeks post-surgery). Cardiac hypertrophy (Left ventricular mass, LVM) evaluated by echocardiography over time **(i)** and at end-time point (3 weeks post-surgery) **(j)**, and at time of sacrifice by the heart weight to tibia length ratio **(k)**. **l-o,** Histological analysis of sham- and TAC-operated cWT and cKO mice (N=5-8). Quantification of cross-sectional cardiomyocyte area **(l)** and representative images **(m)** of wheat-germ agglutinin immunofluorescence staining, at 3 weeks post-surgery. Scale bar, 100µm. Quantification of fibrosis **(n)** and representative images **(o)** of collagen staining with Picrosirius red, at 3 weeks post-surgery. Scale bar, 100µm. **p-r,** ANP **(p)**, BNP **(q)** and β-MHC **(r)** mRNA level in heart mice, normalised to *Rpl32* (N=8-14). Each point represents an independent animal. Values are expressed as mean ± SEM. Statistical analyses are performed using two-way ANOVA followed by Bonferroni post hoc test. \**p<0.05*.

### Deletion of skNAC in adults analogously accelerated hypertrophy and heart failure under basal and pressure overload conditions

One may wonder whether the severe cardiac phenotype generated by constitutive skNAC deletion could result from developmental defect. To address this question, an inducible cardiac-specific skNAC knockout model (iKO) was generated using αMHC-MerCreMer^+/−^ transgenic mice (**Figure S9a**). In this model, skNAC gene was deleted in adulthood by a transient nucleus translocation of the MerCreMer double fusion protein, induced by tamoxifen administration (**Figure S9a**). Dose of tamoxifen was chosen to avoid any transient cardiac defect as described previously ^32–34^. Tamoxifen-induced MerCreMer stimulation resulted in efficient skNAC deletion at the mRNA and protein levels whereas αNAC was slightly upregulated (**Figure S9b-e**). Predictably, skNAC expression remained unaltered in skeletal muscles (**Figure S9f**). Left ventricular mass and cardiac function were monitored for up to one year. Consistent with the cKO findings, a progressive decline in cardiac systolic function and an increase in left ventricular mass were observed in iKO mice compared with corresponding controls (**Figure S10, Table S6)**.

Acute deletion of skNAC in iKO mice also aggravated TAC-induced HF. TAC surgery was performed 2 weeks after the last injection of tamoxifen. For a similar increase in blood flow velocity in both genotypes (**Figure 6a**), skNAC deletion in adult mice promoted severe HF (**Figure 6b**) with brutal left ventricular dilation, together with ejection fraction and fractional shortening reaching values, after nine weeks of pressure overload, of approximatively 20% and 10%, respectively (**Figure 6c-h, Table S7**). This correlated with more pronounced increase in left ventricular mass under echocardiographic follow-up (**Figure 6i,j**) and heart weight/tibia length ratio after sacrifice (**Figure 6k**). Histological analysis further confirmed adverse remodelling, with enhanced cardiomyocyte cross-sectional area (**Figure 6l,m**) and more pronounced increase in interstitial fibrosis (**Figure 6n,o**). Once again, the classical hypertrophic markers ANP, BNP and β-MHC were unchanged by skNAC deletion (**Figure 6p-r)**. As expected, comparison of the effects of TAC on iKO and cKO revealed similar phenotypes, but with a slower progression of HF in iKO, as confirmed by the lesser reduction in ejection fraction three weeks after surgery (**Figure S11**). Collectively, these results demonstrate that skNAC is required for the maintenance of cardiac structure and function, this protective action being even more critical under conditions of pressure overload.

**Figure 6:**
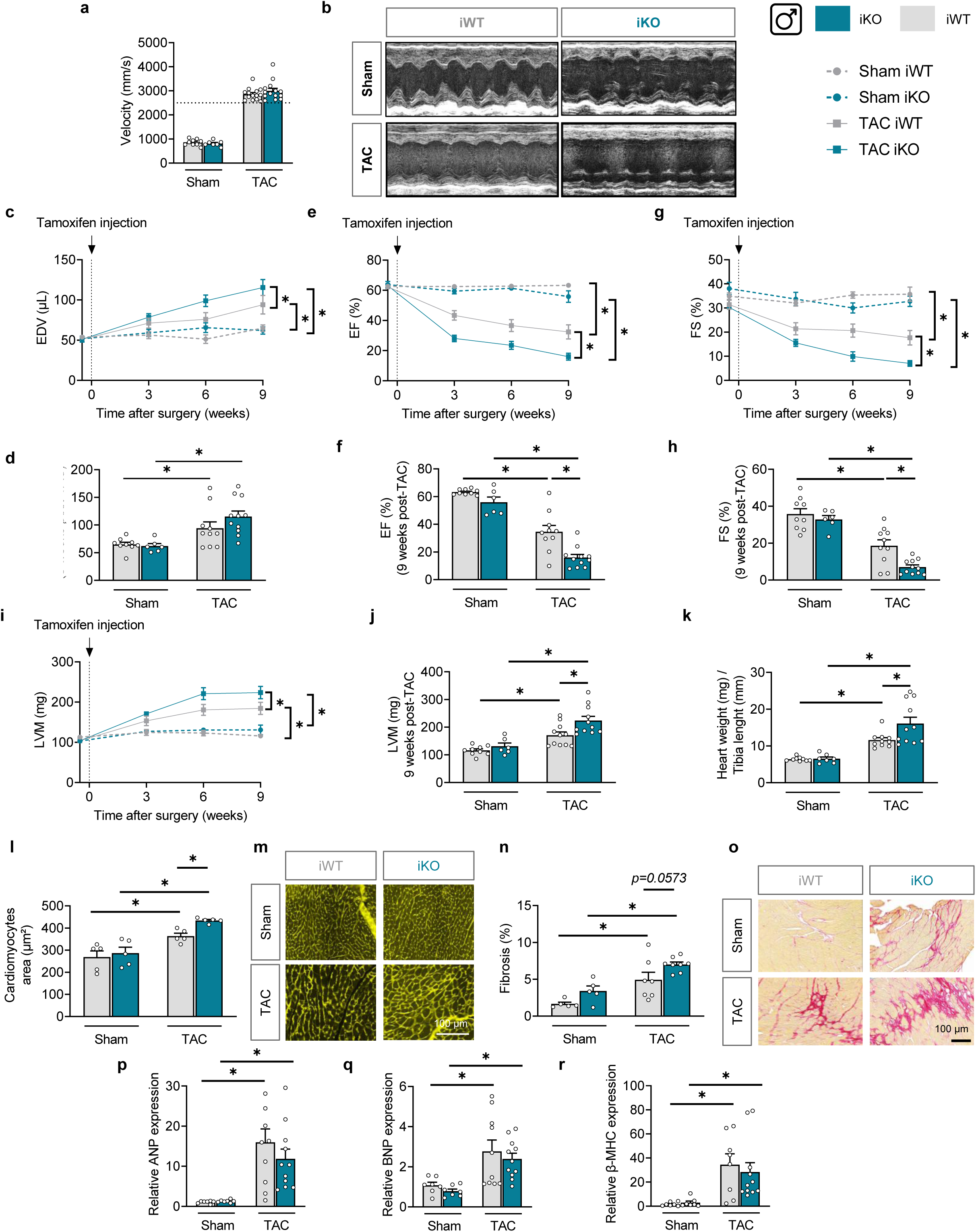
Cardiomyocyte-specific inducible deletion of skNAC in adults analogously accelerated hypertrophy and heart failure under pressure overload conditions. **a-j,** Echocardiographic analysis of sham- and transverse aortic constriction (TAC)-operated cardiomyocyte-specific inducible skNAC knockout (iKO) and control littermate (iWT) mice at 3-, 6-and 9-weeks post-surgery (N=7-14). Quantification of peak aortic velocity at the site of constriction in sham- and TAC-operated iWT and iKO mice **(a)**. Representative image of M-mode in sham-and TAC-operated iWT and iKO mice at 9 weeks post-surgery **(b)**. End-diastolic volume (EDV) **(c, d)**, ejection fraction (EF) **(e, f)** and fractional shortening (FS) **(g, h)** evaluated by echocardiography over time and at end-time point (9 weeks post-surgery). Cardiac hypertrophy (Left ventricular mass, LVM) evaluated by echocardiography over time **(i)** and at end-time point (9 weeks post-surgery) **(j)**, and at time of sacrifice by the heart weight to tibia length ratio **(k)**. **l-o,** Histological analysis of sham- and TAC-operated iWT and cKO mice (N=5-8). Quantification of cross-sectional cardiomyocyte area **(l)** and representative images **(m)** of wheat-germ agglutinin immunofluorescence staining, at 9 weeks post-surgery. Scale bar, 100µm. Quantification of fibrosis **(n)** and representative images **(o)** of collagen staining with Picrosirius red, at 9 weeks post-surgery. Scale bar, 100µm. **p-r,** ANP **(p)**, BNP **(q)** and β-MHC **(r)** mRNA level in heart mice, normalised to *Rpl32* (N=8-14). Each point represents an independent animal. Values are expressed as mean ± SEM. Statistical analyses are performed using two-way ANOVA followed by Bonferroni post hoc test. \**p<0.0*5.

### skNAC co-localises and interacts with ribosomes next to the sarcomeres

After demonstrating that skNAC repression *in vitro* and *in vivo* induces cardiac remodelling and increases susceptibility to HF, we sought to elucidate its underlying molecular mechanisms. Since it is known that αNAC is a chaperone protein associated with ribosomes and that skNAC helps to form sarcomeres during skeletal development, it is tempting to hypothesize that cardiac skNAC could act at the interface of ribosomal function and sarcomere organization in cardiomyocytes, particularly at the Z-lines, where the ribosome is highly enriched. Combined ultrastructural and immunofluorescence approaches were used to further explore this. We first evaluated the subcellular localisation of skNAC using immunofluorescence and confocal microscopy in isolated NRVMs and adult rat ventricular myocytes (ARVMs) (**Figure 7a,b, left panels**). skNAC is excluded from the nucleus and localised in the cytoplasm in a punctate pattern in both neonatal and adult cardiomyocytes. Co-localisation with ribosomal protein S6 (S6, marker of ribosomes) was then performed (**Figure 7a-e**). skNAC displayed strong co-localisation with S6, as visualised by dual image analysis of S6 (in green) and skNAC (in red) (**Figure 7a,b**). This marked skNAC/S6 co-localisation was further quantified by line scan analysis (**Figure 7c,d**) and by measuring the Pearson’s correlation coefficient, which reached values above 0.8 (**Figure 7e**). A proximity ligation assay (PLA) was performed in ARVMs to substantiate direct skNAC/S6 interaction using oligonucleotide-labelled antibodies that, when bound to the two proteins less than 40 nm apart, facilitate a ligase reaction, DNA amplification and subsequent generation of fluorescent signal (**Figure 7f,g**). Strong PLA signals were observed throughout the cytoplasm, consistent with a close association with the ribosomal machinery.

**Figure 7:**
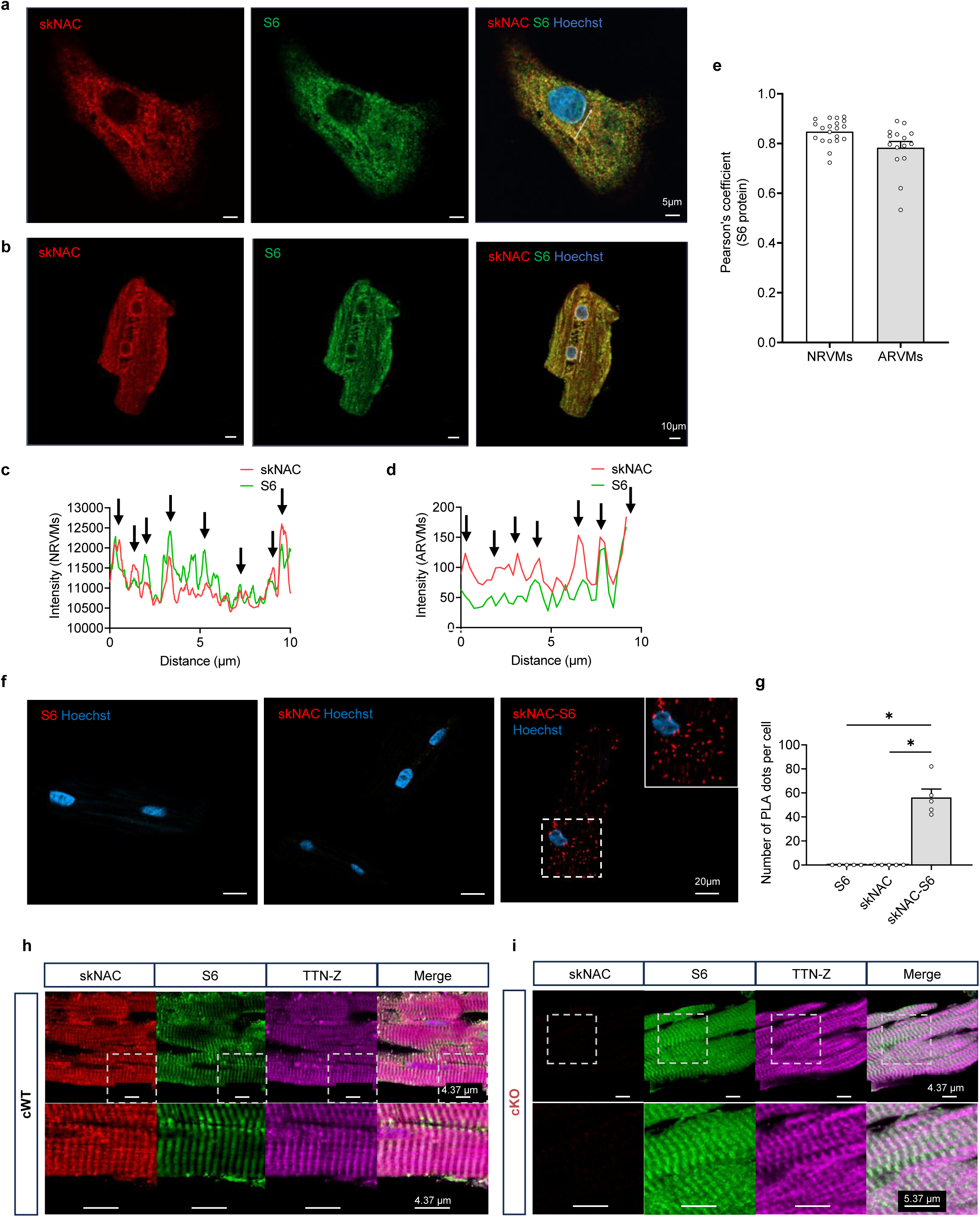
skNAC co-localises and interacts with ribosomal protein S6 in cultured cardiomyocytes. **a-b,** Confocal imaging of immunofluorescence staining in cultured cardiomyocytes. Representative immunostaining images of skNAC (red), ribosomal protein S6 (green), and both merged in the right panel, in NRVMs **(a)** and ARVMs **(b).** Nuclei are stained with Hoechst (blue). Scale bar, 5µm (NRVMs) and 10µm (ARVMs), respectively. **c-d,** Line scan profiles showing overlapping fluorescence intensity of skNAC (red) with S6 (green) in NRVMs **(c)** and ARVMs **(d)**. **e,** Quantification of skNAC/S6 co-localisation using Pearson’s correlation coefficient in NRVMs and ARVMs (N=15-20). **f-g,** *In situ* proximity ligation assay (PLA) analysis of protein complexes consisting of skNAC and S6. Representative images **(f)** and quantification **(g)** of PLA signals in ARVMs. Red dots indicate PLA-positive signals, and nuclei are stained with Hoechst. Each dot in the graphic represents an independent biological cell (N=6-20). **h-I**, Confocal imaging of immunofluorescence staining of heart sections from 3-month-old cWT and cKO mice. Representative immunostaining images of skNAC (red), S6 (green), TTN-Z (N-terminal part of Titin protein, magenta), and all merged in the right panel, from cWT **(h)** and cKO **(i)** mice. Data are presented as mean ± SEM. Statistical analyses are performed using one-way ANOVA followed by Bonferroni post hoc test. *p<0.05.

Co-localisation of skNAC with S6 was also verified by immunofluorescence confocal microscopy on heart tissue samples of young adult cWT and cKO mice, before the onset of the cardiac dysfunction (**Figure 7h-i and S13**). Z-disks were also visualized using the TTN-Z antibody directed against the N-terminal part of Titin protein ^35^. As previously shown ^36,37^, S6 co-localised with TTN-Z in the Z-disks. As observed in isolated cultured cardiomyocytes, skNAC strongly co-localised in cWT samples with S6 to the Z-disks of cWT mouse hearts. The fact that the skNAC signal is more punctiform than that of S6 and TTN-Z indicated a more specific localisation of skNAC. skNAC signals completely disappeared in cKO samples, definitively confirming the specificity of our home-made antibody.

Because αNAC is a ribosome-associated chaperone that also prevents aberrant translocation of nascent peptides to the endoplasmic/sarcoplasmic reticulum (ER/SR), we tested whether skNAC might co-localise with ER/SR subcellular structures using antibody against the KDEL signal peptide marker of proteins retained within the ER/SR (**Figure S12a-e**). skNAC only partially colocalised with the ER/SR. Indeed, only a few intensity peaks were observed simultaneously during line-scan analysis, with a Pearson correlation coefficient of between 0.5 and 0.6 (**Figure S12a-e**).

Together, these results indicate that skNAC predominantly localises to the ribosomal compartment, particularly enriched in Z-disks, and that its loss will finally disrupt myofibril integrity, linking skNAC to translational quality control and contractile capacity.

### skNAC deletion leads to rapid changes in the metabolic genetic profile

Untargeted transcriptomic and proteomic profiling was conducted on cardiac tissue from iKO mice two weeks after tamoxifen-mediated deletion of skNAC to elucidate the primary molecular perturbations, i.e. well before the first signs of dysfunction became apparent. High-depth profiling identified 37,593 transcripts (**Figure 8a-d**), of which 793 (2,1%) were downregulated (log2FC≤1) and 992 (2,6%) upregulated (log2FC≥1). In parallel, MS-based proteomics quantified 2,613 proteins (**Figure 8e-h**). skNAC deletion was confirmed in both datasets, validating the experimental model. Principal component analysis (PCA) revealed a clear separation between iKO and iWT samples at both the transcript and protein levels, indicating a genotype-driven molecular signature, even after such a short skNAC deletion period (**Figure 8b,f**).

**Figure 8:**
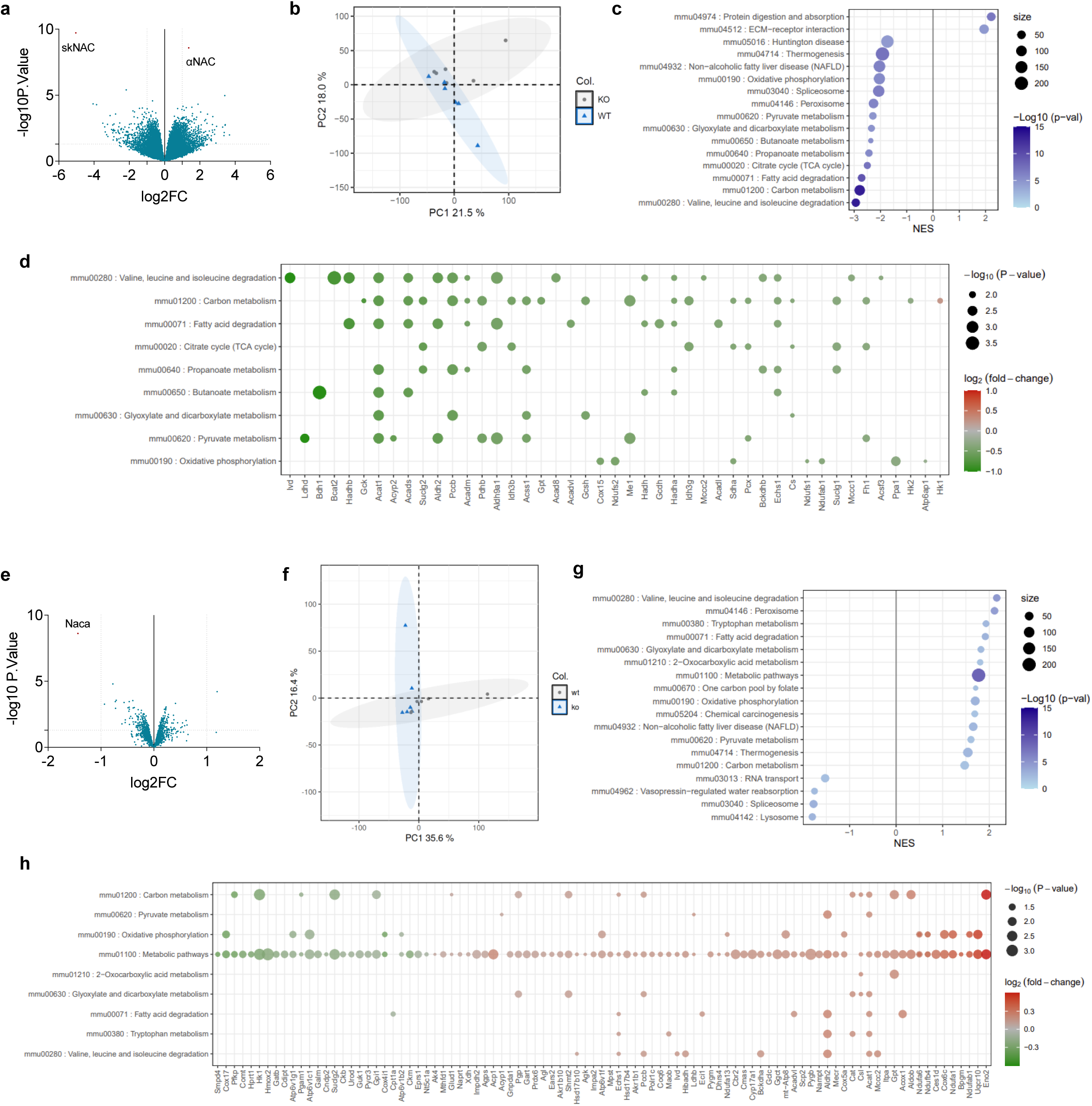
Integration of transcriptome and proteome analyses highlights proteins decoupled from mRNA levels in iKO mice. Total RNAs were extracted, and RNA sequencing analysis was performed (n=6 mice/group). **a**, Volcano plot representation of differentially expressed transcripts in iKO, as compared to iWT, two weeks after tamoxifen administration. **b**, Principal component analysis (PCA) plot of transcriptomic analysis in iKO and iWT mice. **c**, Gene set enrichment analysis (GSEA) of transcriptomic data in iKO compared to iWT. Kyoto Encyclopedia of Gene and Genomes (KEGG) database was used. Dot size represents the number of genes within the pathways. The colour indicates the -log10 P-value (p-val), ranging from low (light blue) to high (dark blue). NES = normalized enrichment score. To prioritize results of primary biological interest, pathways deemed non-relevant to the specific experimental context were excluded from the visualization. **d**, Individual representation of modulated genes comprised in GSEA pathways. Dot size represents the -log10 P-value. The color indicates the -log2 (fold-change) (FC) in iKO compared to iWT, ranking from low (green) to high (red). Genes displayed in the dot plot were selected using an FDR threshold of 0.3. **e**, Volcano plot representation of differentially abundance proteins in iKO, as compared to iWT, two weeks after tamoxifen administration. **f**, Principal component analysis (PCA) plot of proteomic analysis in iKO and iWT mice. **g**, GSEA of proteomic data in iKO compared to iWT. KEGG database was used. Dot size represents the number of genes within the pathways. The colour indicates the - log10 P-value (p-val), ranging from low (light blue) to high (dark blue). NES = normalized enrichment score. To prioritize results of primary biological interest, pathways deemed non-relevant to the specific experimental context were excluded from the visualization. **h**, Individual representation of modulated proteins comprised in GSEA pathways. Dot size represents the - log10 P-value. The colour indicates the -log2FC in iKO compared to iWT, ranking from low (green) to high (red). Genes displayed in the dot plot were selected using an FDR threshold of 0.3.

Based on the transcriptomic dataset, differential expression and gene set enrichment analyses (GSEA) using Kyoto Encyclopedia of Genes and Genomes (KEGG) terms highlighted a downregulation of mitochondrial and oxidative gene programs at mRNA level, including oxidative phosphorylation, the tricarboxylic acid cycle, fatty acid β-oxidation, and branched-chain amino acid catabolism (**Figure 8c**). Representative genes within these affected modules include *Acat1* and *Aldh2* (metabolic detoxification and acetyl-CoA metabolism), as well as core components of the respiratory chain and its assembly machinery such as *Cox15*, *Ndufs2* and *Uqcrfs1*, pointing to broad impairment of mitochondrial substrate supply and electron-transfer functions (**Figure 8d, Figure S14a**). Of note, proteomic GSEA performed on the same samples uncovered a different, quite opposite, pattern (**Figure 8g**). Major metabolic pathways and mitochondrial processes appeared preserved or even positively enriched at the protein level, despite transcriptional repression. Indeed, while some mitochondrial proteins mirrored the transcriptomic downregulation, others, including components such as *Acat1*, *Cox6c*, *Uqcrb*, and *Ndufa7* (**Figure 8h, Figure S14b**), showed upregulation in skNAC iKO mice. Such discordance between transcript and protein expression supports a model in which skNAC looks required to maintain tight coupling between transcriptional and translational programs, particularly linked to mitochondrial metabolic homeostasis.

## Discussion

Increased protein synthesis, as well as sarcomeric and metabolic reorganisation are central features of cardiac remodelling and progression to HF. In this study, we identify the muscle-restricted *NACA* isoform skNAC as an essential regulator of cardiac homeostasis and pressure overload-induced remodelling. To summarise, we show that skNAC expression is reduced in preclinical models of hypertrophy and in hypertrophic human hearts. Cardiac skNAC knockout triggers a non-canonical hypertrophic response and systolic dysfunction both at basal and under hemodynamic stress, whereas skNAC overexpression restrains pro-hypertrophic growth, supporting an anti-hypertrophic and protective function. skNAC is known to contribute to myofibrillogenesis and cardiogenesis ^11^, yet its function in the adult heart has remained largely unexplored. Our study provides mechanistic evidence that loss of skNAC provokes maladaptive cardiac hypertrophy by disrupting the molecular pathways that couple translational protein quality control to metabolic and sarcomeric homeostasis. Notably, skNAC is downregulated during hypertrophic stress, compare to its canonical αNAC counterpart, underscoring to isoform-specific regulation of *NACA* splicing during hypertrophy.

Constitutive knockout of skNAC causes severe cardiac malformations and embryonic lethality due to defects in ventricular cardiomyocyte maturation ^16^. In our models, cardio-specific skNAC deletion occurs later, specifically at mid-embryonic period for cKO and at adult stage for iKO, leading to a less severe but still highly deleterious phenotypes. Given that αMHC-driven Cre recombinase is progressively expressed from embryonic day 8.5 in cKO mice ^38^, we could expect a progressive deletion of skNAC in the second half of embryonic development. Such skNAC deletion induces an early lethal phenotype (around six months) associated with progressive systolic dysfunction and severe left ventricular dilation. Deletion of skNAC in adult mice triggered similar cardiac remodelling and dysfunction, slightly improving mouse survival. Overall, these observations highlight a dual and essential role of skNAC in both the development and maintenance of adult cardiac tissue. Because skNAC is reduced in hypertrophy and HF, we tested whether its deletion increases susceptibility to pressure overload. Following TAC, cKO mice rapidly developed severe HF and died within three weeks, whereas iKO mice exhibited a milder but progressive phenotype, reaching critical HF by nine weeks. These findings demonstrate that loss of cardiac skNAC alone is sufficient to exacerbate dysfunction under hemodynamic stress. Mechanistically, skNAC was initially described as a transcriptional co-activator during myogenic differentiation through Smyd1 interaction, and nuclear to cytosol translocation of Smyd1/skNAC has been reported in skeletal muscle models ^39^. In adult cardiomyocytes, our immunofluorescence study rules out the possibility of a nuclear localisation of skNAC. In contrast, we provide evidence that skNAC deficiency leads to myofibrillar disorganisation, inter-sarcomeric vesiculation, and autophagic vacuole accumulation. Morphologically, we showed that skNAC strongly co-localises with ribosomes at the Z-disks, consistent with ribosomal interactome data identifying skNAC as a component associated with translational machinery ^20^. Given the established role of αNAC as a ribosome-associated chaperone that prevents inappropriate translocations ^4^, our observation support the idea that skNAC may function as a muscle-specific factor at the ribosome-sarcomere interface. This is in line with work in zebrafish showing that skNAC is required for ordered myofibril assembly ^17^. In line, Kehat and colleagues demonstrated that cardiomyocytes preserve sarcomere integrity through spatially confined mechanisms that include localisation of sarcomeric RNAs and ribosomes to enable local protein synthesis, coupled to targeted removal of excess proteins by localised ubiquitin-proteasome pathway ^36^. Co-localisation of skNAC with ribosomal protein S6 at Z-disks strongly suggests a potential role for skNAC in the turnover of sarcomeric proteins. Given that αNAC functions as a chaperone binding to nascent polypeptides emerging from the ribosome, it is plausible that skNAC acts in a similar manner as a muscle-specific chaperone, facilitating the proper targeting and folding of sarcomeric proteins. skNAC deletion results in profound myofibrillar disorganisation, which may stem from impaired myofibrillogenesis due to defective chaperoning or delivery of newly synthesised sarcomeric proteins required for the continuous renewal of the contractile apparatus. Observation of autophagic vacuoles by electron microscopy possibly reflects accumulation of misfolded sarcomeric proteins and/or impairment of autophagic processes. Future ribosome profiling will identify the proteins that depend on skNAC and clarify how skNAC supports sarcomere assembly.

Beyond this structural remodelling, which take months to become apparent, skNAC deficiency triggers early alterations in the expression of metabolism-related proteins after tamoxifen-induced gene deletion. As early as two weeks after induced deletion, transcriptomic and proteomic analyses showed a downregulation of mitochondrial and oxidative mRNA transcripts (oxidative phosphorylation, tricarboxylic acid cycle, fatty acid β-oxidation), despite the preservation or increase of their corresponding protein levels. A recent study by Haddad et al. similarly reported that skNAC reduction alters mitochondrial proteins and impairs Krebs cycle and respiratory chain function ^20^. These findings support a model in which skNAC contributes to mitochondrial proteome and energetic homeostasis, analogous to the canonical NAC complex ^4,5^, and consistent with early observations in C2C12 myocytes ^40^. The apparent discrepancy between mRNA and protein expression may reflect differences in transcript and protein turnover kinetics, compensatory translational regulation, altered protein stability, and/or feedback mechanisms whereby sustained accumulation of proteins contributes to transcriptional repression of the corresponding genes. skNAC expression is modulated by muscle-enriched RNA-binding proteins (RBPs) that promote the specific inclusion of exon 3 in the mRNA ^11,41^. Rbm24 is implicated in skNAC pre-mRNA maturation and is known to cooperate with Rbm20 in the splicing of other muscle-specific pre-mRNA, including Titin ^12,42^. Here, we find that Rbm24 or Rbm20 knockdown decreases skNAC expression without affecting αNAC expression, identifying skNAC as a novel target of Rbm20/Rbm24-dependent splicing control. Rbm24 expression is reduced *in vitro* and *in vivo* during hypertrophic stress, whereas Rbm20 reduction is only evident *in vitro*. This suggests a pivotal role of Rbm24 in maintaining skNAC levels and preventing cardiac hypertrophy. Consistently, Rbm24 cardiac-specific knockout mice develop progressive dilated cardiomyopathy, HF, impaired sarcomere assembly, dysregulated Ca^2+^ handling and lethality ^43^. Rbm24 is involved in the maturation of many different pre-mRNAs, including gene involved in fibrosis, making it difficult to decipher the relative importance of skNAC knockdown in Rbm24 knockout phenotype. Interestingly, Rbm20 knockout mice also exhibit dilated cardiomyopathy associated with proarrhythmic changes in Ca^2+^ handling ^42^. Consistently, GWAS analysis of *NACA* gene identified SNPs significantly associated with specific cardiac traits including conduction abnormalities, supporting a possible role for *NACA* variation in various cardiac pathologies.

If skNAC deficiency promotes HF, our study is also the first to show that skNAC overexpression prevents cardiomyocyte hypertrophy. Therefore, modulation of skNAC levels or activity emerges as a potential therapeutic strategy to strengthen cardiomyocyte protein quality control and prevent progression to HF. Given their wide range of targets, it is certainly not advisable to target Rbm proteins to promote skNAC expression. Indeed, Rbm24 overexpression is harmful in the adult heart, as it increases the expression of TGFβ-signalling and extracellular matrix genes, promoting fibrosis ^44^. Considering these adverse effects, alternative strategies to modulate skNAC activity should be explored. In this context, modulation of skNAC O-GlcNAcylation looks to be an attractive alternative. Indeed, besides its regulation at the mRNA level, skNAC has been previously shown to be modified by O-GlcNAcylation ^18^. Recent literature pointed out O-GlcNAcylation as an important player in cardiac hypertrophy and HF ^19,45^. Accordingly, O-GlcNAcylated skNAC was systematically detected in the mass spectrometry-based non-targeted O-GlcNAc proteomic approach that we previously established and that we applied to cardiac samples ^46,47^. Such skNAC O-GlcNAcylation can be confirmed in mice treated with the O-GlcNAcylating agent NButGT ^21^ (**Figure S15**), suggesting a conceivable complexity in skNAC regulation.

Some limitations can be found in our study. First, αMHC-Cre toxicity in male is a recognised confounder, already well described in the literature. We mitigated this by exploring corresponding female mouse groups and by reproducing key phenotypes in iKO animals. Second, skNAC overexpression has been restricted to *in vitro* studies. However, the extremely large size of skNAC limits the use of AAV viral constructions to validate our data *in vivo*.

To summarise, our data demonstrate that skNAC is a pivotal regulator of cardiac proteostasis, sarcomere integrity, metabolic homeostasis, and contractile function. By integrating *in vitro*, *in vivo*, and human data, we demonstrate that skNAC deletion is sufficient to drive maladaptive hypertrophy and systolic dysfunction through mechanisms linking translational dysregulation to energetic imbalance and structural disorganisation, whereas its overexpression provides protection against pro-hypertrophic stimuli (**Figure 9**).

**Figure 9:**
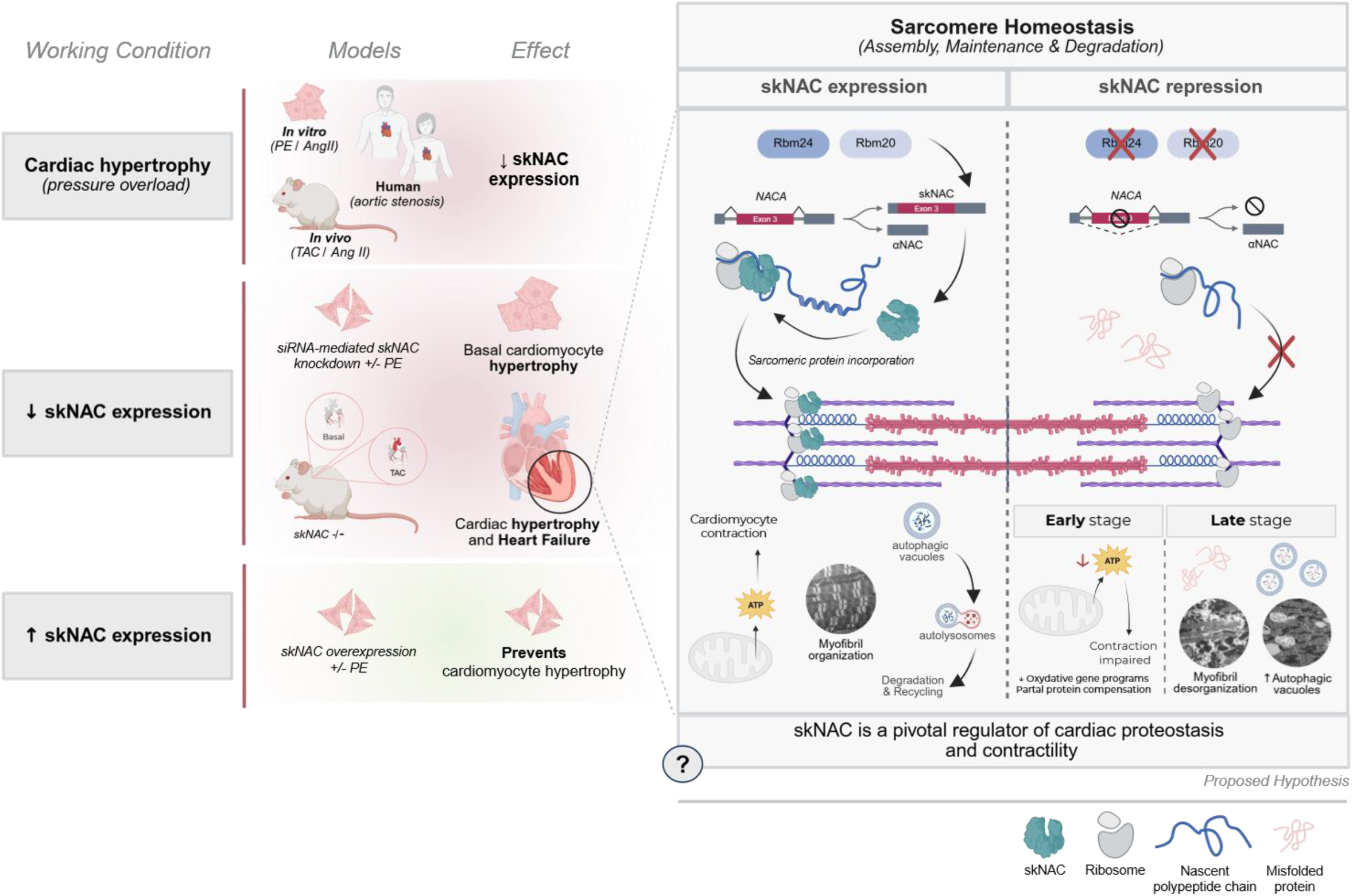
skNAC regulates cardiac proteostasis and preserves sarcomere integrity during hypertrophic remodelling. skNAC expression is reduced in experimental models of pressure overload and in response to pro-hypertrophic stimuli (TAC, AngII, PE), as well as in human aortic stenosis. skNAC loss leads to maladaptive hypertrophy and systolic dysfunction via imbalances in translation and energy metabolism. Conversely, overexpression of skNAC protects against pro-hypertrophic stress.

## Methods

A supplemental methods’ section is available in the Supplemental Materials. All data, methods and materials from this study are available from the corresponding author upon reasonable request.

### Animal care

All experiments and animal handling were approved by the UCLouvain Faculty Ethics Committee for Animal Experimentation (Comité d’éthique facultaire pour l’expérimentation animale, 2021/UCL/MD/016) and conformed to the American Heart Association Guidelines for the Use of Animals in Research. Animals were maintained under a 12-hours light/12-hours dark cycle. They were monitored daily and had unrestricted access to water and standard chow (or other specified) diets. All animals shared identical housing conditions, including cage, location, and litter, where they were maintained under standard environmental conditions. No acclimatization period was required, as all animals originated from the same breeding facility. Only the male from one litter was housed alone in a separate cage. Euthanasia was performed via an intraperitoneal injection of a solution (1 µl/g) containing ketamine (100 mg/ml) and xylazine (20 mg/ml).

### Patient Samples and Biopsy Collection

Human cardiac biopsies from patients with aortic stenosis were obtained during surgery at Cliniques Universitaires Saint-Luc (Brussels, Belgium), in accordance with the regulations of the local ethics committee (Comité éthique hospitalo-facultaire des Cliniques Universitaires Saint-Luc). Written informed consent was obtained from all patients prior to surgery. Cardiac biopsies were immediately frozen in liquid nitrogen and stored at −80°C. Detailed patient characteristics are provided in **Table S1**.

### Generation of plasmids overexpressing skNAC

The murine skNAC sequence cloned in the pCI vector was obtained from the laboratory of Prof. B. Munz (University of Tübingen, Germany). The plasmid, allowing skNAC expression under the control of the cytomegalovirus promoter (pCMV), was recovered from nitrocellulose, amplified in E. coli NEB® 5-Alpha Competent cells, and purified using the Genopure Plasmid Maxi kit (Roche).

Sequencing revealed the absence of a 1,146 bp fragment within exon 3. To restore the full sequence, the missing region was amplified from mouse heart cDNA by PCR (using Q5® D High-Fidelity DNA polymerase, NEB), digested together with the vector using *PciI* and *PspXI*, and purified by phenol:chloroform extraction. The insert was then ligated into the plasmid backbone, transformed into competent *E. coli*, and positive clones were validated by Sanger sequencing. A second plasmid allowing the expression of skNAC under the control of the troponin T promoter (pTNT) was obtained by Addgene (#69915).

### Generation of plasmids overexpressing HA-tagged skNAC

Site-directed mutagenesis was performed by PCR amplification (using Q5® Hot Start DNA polymerase, NEB) to introduce a hemagglutinin (HA) tag into the skNAC coding sequence contained in the pCI vector. We generated constructs with the tag either at the N-terminus (immediately after the start codon) or at the C-terminus (before the stop codon). Correct insertion of the HA tag was confirmed by Sanger sequencing. Validated constructs were then amplified on a large scale using a Maxi-Prep kit for downstream applications.

### Generation of a vector allowing skNAC and EGFP co-expression

A plasmid was generated to allow the cardiomyocyte-specific expression of skNAC under pTNT, as well as the ubiquitous expression of enhanced green fluorescent protein (EGFP) under pCMV promoter. The construct was obtained by homologous recombination using the In-Fusion® method (Takara). The parental plasmids, pTNT-skNAC and pCMV-EGFP, were linearized and amplified by PCR with primers carrying homologous sequences, enabling recombination into a single construct of 11,684 bp containing two independent expression cassettes.

### Preparation and treatment of neonatal rat ventricular myocytes

Neonatal rat ventricular myocytes (NRVMs) were isolated and cultured under aseptic conditions as previously described ^21^. Before treatment, NRVMs were incubated in serum-free IMDM for 2 hours, and then treated with pro-hypertrophic agents, phenylephrine (PE, 20 µM) or Angiotensin II (AngII, 100 nM) for 24 hours. At the end of the treatment, NRVMs were washed twice with ice-cold phosphate-buffered saline (PBS), and then lysed in a lysis buffer containing 50 mM HEPES (pH 7.5), 50 mM KF, 1 mM KPi, 5 mM EDTA, 5 mM EGTA, 1% (vol:vol) Triton and a cocktail of protease, phosphatase and O-GlcNAc inhibitors (0.5 mM PMSF, 1 mM benzamidine HCl, leupeptin 1 μg/mL, pepstatin A 1 μg/mL, 1 mM DTT, 1 mM vanadate, 1 mM alloxan, 1 mM PUGNAc). Lysates were centrifuged (13,000 g, 15 min, 4°C), and supernatants were stored at - 80°C.

### Isolation, culture and treatment of adult rat ventricular myocytes

Adult rat ventricular myocytes (ARVMs) were isolated from 250g male Wistar rats, as described previously ^48^. Isolated heart was retrogradely perfused through the aorta with Ca^2+^-free Krebs-Henseleit buffer. Hearts were then digested by adding 0.2 mM Ca^2+^, 1 mg/mL type II collagenase, and 0.4% (wt:vol) fatty acid-free BSA, followed by mechanical disruption with scissors. Cardiomyocytes were purified by sedimentation, resuspended in medium 199, supplemented with 2 mM carnitine, 5 mM creatine, 5 mM taurine, 10^−10^ M T_3_, 0.2% (wt:vol) fatty acid-free BSA, and antibiotics, and evenly plated on laminin-coated dishes.

### Small interfering RNA-mediated skNAC knockdown in neonatal rat ventricular myocytes

Small interfering RNAs (siRNAs) were designed to target the exon 3 of skNAC and validated in NRVMs. NRVMs were transfected at the time of plating with either a non-targeting control siRNA (50 nM) or an siRNA targeting skNAC (50 nM) using Lipofectamine^TM^ RNAiMAX transfection, following the manufacturer’s instructions. After 72 hours of transfection at 37°C and 5% CO_2_, the medium was replaced, and, when needed, cells were treated with PE (20 µM) for 24 hours to induce hypertrophy.

### skNAC overexpression in neonatal rat ventricular myocytes

One hour before transfection, NRVMs were incubated in IMDM without P/S and FBS serum. The transfection was performed with skNAC overexpressing plasmids (5µg) or a GFP plasmid (5 µg) as control, using lipofectamine^TM^ 3000 according to the manufacturer’s instruction. After 24 hours of transfection at 37°C and 5% CO_2_, the medium was replaced and cells treated with or without PE.

### Neonatal rat ventricular myocyte size measurement

α-actinin immunofluorescence staining was used to assess NRVMs size as described previously ^21^. Isolated NRVMs were plated on coverslips coated with gelatin and treated as described above. At the end of the treatment, NRVMs were fixed with 4% formaldehyde and permeabilized with 0.2% Triton X-100 for 30 min on ice. Following fixation and permeabilization, cardiomyocytes were blocked with 1% BSA in PBS for 40 min, then incubated for 1 hour at room temperature with a primary anti-α-actinin antibody diluted in 1% BSA in PBS. This was followed by incubation with an Alexa Fluor 594-conjugated anti-mouse secondary antibody. Cells were visualized using a Zeiss AxioImager.z1 microscope, and images were captured at ×20 magnification. Cell size (>100 cells per sample) was measured using ImageJ software.

### Immunofluorescence (co)staining

Immunofluorescence was performed on NRVMs and ARVMs. Cells were fixed with 4% paraformaldehyde, permeabilized with 0.2% Triton X-100 0.2% Triton, and blocked with 1% BSA in PBS for 40 min. Then, cells were incubated overnight at 4°C with primary antibodies (from two different species) raised against the two proteins of interest (rabbit anti-skNAC and mouse anti-S6 protein antibodies). This was followed by incubation with an Alexa Fluor 594-conjugated anti-rabbit secondary antibody and an Alexa Fluor 488-conjugated anti-mouse secondary antibody. Nuclei were counterstained with Hoechst for 10 min at room temperature. Cells were visualized using a LSM800 confocal microscope (Carl Zeiss) with a 63x objective lens. Colocalization was evaluated by co-staining and quantified using Pearson’s correlation coefficient. Overlapping of staining intensity of the two markers was also evaluated across straight lines of 10 µm positioned in the cytoplasmic region.

### Measurement of gene expression level

Total RNA was extracted using the RNeasy mini-kit (Qiagen), according to the manufacturer’s instructions, treated with DNAse I, and quantified by NanoDrop spectrophotometer (Thermo fisher Scientific). Reverse transcription was carried out for 45 min at 37°C with 1 µg of total RNA, using oligo(dT) primers (primer poly(dT, Roche) and Moloney Murine Leukemia Virus Reverse Transcriptase (Invitrogen). Quantitative PCR (qPCR) was performed in duplicate for each sample on an IQ5 (Bio-Rad) using the qPCR Core kit for SYBR^®^ Green I. The expression levels of selected gene transcripts were calculated, averaged, and normalized to the housekeeping gene *Rpl32* (**Table S9**), following ΔΔCt method.

### Assessment of gene copy number

Plasmids containing the cDNA of the target genes were used to establish a standard curve, allowing quantification of the absolute number of mRNA copies in each sample. The copy number was calculated using the following formula: Copies/µL=(quantity (ng) x 6.022 x 10^23^) / (Size (bp) x 660 x 10^-9^). Each sample was amplified in duplicate, including both the target gene and the reference gene (Rpl32), which served as an internal control. Data were analyzed by calculating the average copy number for each sample, corrected for the reference gene using the ΔSQmean method. The final copy number values were normalized to the Rpl32 to account for variations in input cDNA.

### Western Blot analysis

Lysate supernatants (20 µg of total protein) extracted from heart homogenate or cells were loaded and separated by sodium dodecyl sulfate-polyacrylamide gel electrophoresis gel (SDS-PAGE) and transferred onto PVDF membrane. After blocking in 5% milk in TBS-Tween 20 (0.1%), membranes were incubated with appropriate antibodies to assess protein levels. Detection was performed with HRP-conjugated secondary antibodies and the BM Chemiluminescence Blotting System. Band intensities were quantified by scanning and analyzing with ImageJ software, and values were normalized to the loading control (anti-eEF2 or anti-GAPDH, performed on the same gel). When present, vertical dotted lines indicate that samples were run on the same gel but are not contiguous.

### TAC-induced cardiac hypertrophy

Eight- to twelve-week-old male and female skNAC-deficient mice (cKO or iKO) and their control littermates (cWT or iWT) underwent transverse aortic constriction surgery to induce cardiac hypertrophy and HF ^49^. A 0.5 cm horizontal incision long was made in the second intercostal space of anesthetized mice. Under microscope, a constrictive band was placed and tightened around the aorta using a blunt 27G needle and 7-0 nylon thread. In sham-operated mice, the ligature was not tightened. The thorax was closed with polypropylene sutures, and the animals were treated intraperitoneally with buprenorphine (0.1 mg/kg) post-operatively. Doppler measurements of trans-stenotic gradients were performed one week after surgery, and only TAC mice with a velocity greater than 2,500 mm/s were included in the experiment. Three or nine weeks after TAC surgery, mice were euthanized during the morning, and hearts were washed in PBS and KCl (250 mM) to maintain diastole. The apex was freeze-clamped in liquid nitrogen and stored à -80°C, while the upper portion of the heart was fixed in 4% formaldehyde and embedded in paraffin for further analyses. For immunoblotting, 20 mg of tissue was homogenized in 200 µL of RIPA buffer (25 mM Tris-HCl, pH 7.6; 1% NP-40; 1% sodium deoxycholate; 0.1% SDS; 150 mM sodium chloride; 20 mM glycerol-2-phosphate disodium; 50 mM sodium fluoride; and 5 mM sodium pyrophosphate decahydrate) supplemented with protease/phosphatase inhibitor cocktail (Sigma) and O-GlcNAc inhibitors (Alloxan/PUGNAc, 1 mM each).

### Echocardiography measurements

Echocardiographic data acquisition and analysis adhered to the guidelines provided by the European Society of Cardiology Working Group on Myocardial Function ^50^. Cardiac systolic function was assessed by non-invasive transthoracic echocardiography using the Vevo 3100 Imaging System (FUJIFILM VisualSonics, Toronto, Canada) with a MX550D 30 MHz transducer. This procedure was performed on anesthetized animals, initially using 3% isoflurane (v:v), then reduced to 1% (v:v) to keep the animal unconscious with a heart rate of approximately 500 beats/min. cKO and iKO mice were monitored longitudinally under basal conditions, with a bimonthly follow-up starting at eight weeks or one week before tamoxifen injection, respectively. Mice subjected to TAC underwent transthoracic echocardiography before and at three-, six-, and nine-weeks post-surgery. Cardiac parameters were evaluated using two-dimensional (2D) B-mode in the long axis and M-mode. Left ventricular (LV) volumes measured in B-mode long-axis images at end-systole and end-diastole were used to determine ejection fraction (EF %). Fractional shortening (FS %) was calculated using internal LV dimensions and wall thickness (septum and posterior wall) measurements at end-systole and end-diastole obtained from M-mode recordings. LV mass was assessed from long axis measurements and normalized to tibia length measured at sacrifice. LV mass was calculated using the following equation: LV mass = 1.05[(IVS + LVID + LVPW)^3^-(LVID)^3^] where IVS is the interventricular septal thickness, LVID, the LV internal diameter and LVPW, the LV posterior wall thickness. All measurements were performed by the same operator, who was blinded to experimental groups. The complete set of echocardiographic parameters is provided in supplemental Tables S2 – S7.

### Histological analysis

Hearts were excised, cleared of blood in ice-cold PBS, and fixed with 4% formaldehyde for 24 hours at 4°C. Fixed tissues were embedded in paraffin to produce 5 μm-thick sections. Tissue sections were deparaffinized in 100% toluene and rehydrate in 100% ethanol before staining.

#### Cardiomyocytes cross-sectional area measurement

Heart sections were incubated with wheat germ agglutinin/rhodamine (1:150) for 2 hours at room temperature, then washed with PBS and mounted using Dako fluorescent mounting medium. Images were acquired with using an Axioscan.z1 slide scanner (Carl Zeiss), and cell size (>500 cells per sample) was analyzed using Visiopharm software.

#### Fibrosis measurement with Picrosirius red staining

Hearts sections were treated with 3% phosphomolybdic acid for 5 min, washed with distilled water and incubated in 0.1% Picrosirius Red solution for 2 hours at room temperature. Sections were then rinsed in 0.01 M hydrochloric acid for 2 min, washed with distilled water, and mounted using a Sakura coverslipper. Slides were digitized using a Panoramic SCAN II slide scanner (3DHistech) and analyzed using Visiopharm software.

#### Co-localisation skNAC-S6-TTN-Z on cardiomyocytes

Samples were first dehydrated through a graded ethanol series before being embedded in paraffin wax. An RM2235 Leica microtome was then used to prepare 5–7 mm-thick sections. These sections were rehydrated and treated with peroxidase buffer. Antigen retrieval was performed using citrate-EGTA buffer. After washing with PBS, the slides were blocked in 5% BSA and 0.5% Triton X-100 for 1 hour. Slides were then incubated at 4 °C overnight with primary antibodies, (listed in **Table S8**), diluted in PBS containing 0.5% BSA), followed by secondary antibodies. Once stained, samples were mounted in Mowiol containing n-propyl gallate to prevent bleaching. Immunofluorescence images were aquired using a Nikon Eclipse Ti2 confocal laser scanning microscope (Nikon Europe, Amstelveen) equipped with a Nikon Plan Apo ×60 NA 1.4 oil immersion lens. Images were recorded using NIS Elements version 4.3 software ^51^.

### Electron microscopy

Mouse hearts were fixed by retrograde aortic perfusion in 4% paraformaldehyde for 3 hours at room temperature with a flow rate at 5 µL/s. After perfusion and fixation tissue was kept at 4°C in PBS until embedded. Samples, once fixed, were subsequently washed twice in PBS. After 0.5% osmium tetroxide application, samples were counterstained with uranyl acetate in 70% ethanol, dehydrated, and embedded in Durcupan resin (Fluka). Resin blocks were crafted, and ultra-thin sections were produced using a Leica Ultracut S ultramicrotome. Sections were placed on glow-discharge Formvar carbon-coated copper grids. Captured images were obtained using a Zeiss LEO 910 electron microscope, equipped with a TRS Sharpeye CCD camera and the provided software from Tröndle ^52^.

### Evaluation of skNAC and S6 protein interaction by proximity ligation assay

The direct interaction between skNAC and S6 protein was evaluated by Proximity ligation Assay (PLA) on ARVMs, using Duolink In Situ Reagents Orange (Merck) according to the manufacturer’s instructions. Cultured ARVMs were fixed with 4% formaldehyde and permeabilized with 0.2% Triton X-100 for 10 min at RT, followed by incubation in blocking solution for 30 min at 37°C. Fixed cells were, then incubated overnight at 4°C with primary antibodies (rabbit anti-skNAC and mouse anti-S6 protein antibodies). Oligonucleotide-labelled secondary antibodies were added for 1 hour, followed by ligation, amplification and hybridization with fluorochrome-coupled oligonucleotides. Fluorescent signals were visualized using an LSM800 confocal microscope (Carl Zeiss) with a 63x objective lens, and positive PLA signals were quantified using ImageJ software.

### RNA sequencing

Total RNA was isolated from the LV of iKO and iWT mice 2 weeks post tamoxifen administration, and treated with DNAse as described above. RNA concentration was quantified using Qubit 4 Fluorometer (Thermo Fisher Scientific) and RNA integrity was evaluated on the Agilent 2100 Bioanalyzer. All samples exhibited high RNA quality, with RNA integrity number (RIN) values > 8. Libraries were prepared starting from 150 ng of total RNA using the KAPA RNA HyperPrep Kit with RiboErase according to the manufacturer’s recommendations. Libraries were equimolarly pooled and sequenced on a single lane on an Illumina NovaSeq 6000 platform. All libraries were paired-end (2 × 100 bp reads) sequenced and a minimum of 50 million of paired-end reads were generated per sample. Sequencing data were analysed using the Automated Reproducible MOdular workflow for preprocessing and differential analysis of RNA-seq data (ARMOR v1.5.4) pipeline ^53^. In this pipeline, reads underwent a quality check using FastQC (Babraham Bioinformatics). Quantification and quality control results were summarised in a MultiQC report before being mapped using Salmon^54^ to the transcriptome index which was built using all Ensembl cDNA sequences obtained in Mus_musculus.GRCm38.cdna.all.fa file.

To evaluate expression at multiple resolutions, estimated transcript and gene abundances were imported into R (v4.5.2) using the tximeta package ^55^. Differential expression analysis was then performed at both the gene and transcript levels using edgeR, applying the Benjamini-Hochberg false-discovery rate correction for multiple comparisons.

Gene Set Enrichment Analysis (GSEA) was performed against the KEGG pathway database using the fgsea v1.36.0 Bioconductor package ^56^.

All transcriptomic data visualizations, including PCA, differential expression plots and enrichment score were generated using the R/ggplot2 ecosystem.

### Mass spectrometry

20 mg of freeze-clamped hearts from the LV of iKO and iWT mice 2 weeks post tamoxifen administration, were homogenized in 200 μL in RIPA lysis buffer (25 mM Tris-HCl, pH 7.6; 1% NP-40; 1% sodium deoxycholate; 0.1% SDS; 150 mM sodium chloride; 20 mM glycerol-2-phosphate disodium; 50 mM sodium fluoride; and 5 mM sodium pyrophosphate decahydrate) supplemented with protease/phosphatase inhibitor cocktail (Sigma) and O-GlcNAc inhibitors (Alloxan/PUGNAc, 1 mM each).

Peptides separation was performed using a C18 reversed-phase analytical column (Thermo Scientific) on an Ultimate 3000-nLC RSLC system. Peptides were labelled using Tandem Mass Tag (TMT) reagents according to the manufacturer’s instructions, allowing multiplexed sample analysis. Labelled peptides were pooled prior to LC-MS/MS analysis. Peptides were subjected to Nano-Spray-Ionization source followed by tandem MS/MS in a tribrid Fusion Lumos Orbitap analyser coupled online to the nano-LC. Data were acquired in a data-dependent acquisition mode with precursor ion detection in the Orbitrap, followed by fragmentation and detection of TMT reporter ions for quantification. The resulting MS/MS data were processed using Sequest HT search engine within Proteome Discoverer 2.5 against a mouse protein database obtained from Uniprot. Trypsin was specified as cleavage enzyme allowing up to 2 missed cleavages, 4 modifications per peptide and up to 5 charges. Mass error was set to 10 ppm for precursor ions and 0.1 Da for fragment ions. Oxidation on methionine, carbamidomethyl on cysteine were considered as variable modifications. False discovery rate (FDR) was assessed using Percolator and thresholds for protein, peptide and modification site were specified at 1%. The filtered Sequest HT output files were grouped at the protein level, and protein abundance was quantified based on TMT reporter ion intensities extracted from MS/MS spectra within Proteome Discoverer.

Statistical analyses were performed in R (v4.1.1). To ensure high-confidence identifications, only proteins detected in at least 4 out of 6 biological replicates per group were retained. Following Log2-transformation and Probabilistic Quotient Normalization (PQN), differential expression between iKO and iWT groups was modeled using the limma package. A linear model with empirical Bayes moderation was applied, treating ’group’ as the primary factor. Multiple testing was accounted for using the Benjamini–Hochberg False Discovery Rate (FDR) method. Proteins were classified as differentially expressed with high confidence (q < 0.1) or very high confidence (q < 0.05). Functional enrichment was subsequently assessed via the fgsea package (v1.36.0) using the Kyoto Encyclopedia of Genes and Genomes (KEGG) database.

All proteomic data visualizations, including PCA, differential expression plots and enrichment score were generated using the R/ggplot2 ecosystem.

### Statistical analysis

The sample size for *in vivo* experiments was determined considering the procedural complexity with a Cohen effect size of 1.5 and an anticipated loss rate of 25%. The ‘*N*’ value indicates the number of animals used for *in vivo* experiments and the number of biological replicates for *in vitro* assays. In addition, each *in vitro* experiment was performed in duplicate or triplicate to ensure reproducibility. All data are presented as means ± SEM. The normality of continuous variables was assessed using the Shapiro-Wilk method and homogeneity of variances was verified prior to parametric testing. Outlier detection was performed using the Grubbs test (ESD method, extreme studentized deviate), with a significance level of 0.05. For comparisons between two groups, (un)paired Student t-tests were applied. For multiple comparisons, one-way or two-way analysis of variance (ANOVA) followed by Bonferroni post hoc correction was used. Survival analyses were performed using the log rank test. A p-value < 0.05 was considered statistically significant. All statistical analyses and graphs were generated using GraphPad Prism 9.1.2 software. Some parts of the figures were created with a licensed version of BioRender.com

## Data availability

All data pertaining to this work are shown in the text, figures and Supplementary Information. Unedited gels utilized for figures are available in Supplemental Material.

## Omics data availability

The mass spectrometry proteomics data have been deposited to the ProteomeXchange Consortium via the PRIDE partner repository with number accession PXD077524, while the transcriptomics data have been deposited to the Gene Expression Omnibus with the reference GSE325002. Please find the following dataset identifiers.

## Acknowledgments

We thank Marine De Loof, Eline Stukkens and Amélie Cammaert for their technical support. The majority of the microscopy images presented here were captured with the help of the 2IP Imaging Platform of our institute (2IP - RRID:SCR_023378). We thank Benjamin Lauzier (University of Nantes) for kindly providing NButGT. The original submitted version of this manuscript was first deposited on the preprint server bioRxiv on 26 October 2025 ^57^.

## Authors’ contributions

LBe has full access to all of the data in the study and takes responsibility for the integrity of the data and the accuracy of the analyses; Concept and design: LG, JDo, LBu, AU, WAL, YA, EH, DV, BB, LD, CBo, JA and LBe; Acquisition, analysis, or interpretation of data: LG, JDo, EV, JDr, HV, EH, LD, AU, DV, BB, WAL, HE, JA, AM, LBu and LBe; Drafting of the manuscript: LG, NF, EV, MR and LBe; Critical revision of the manuscript for important intellectual content: LG, NF, EV, MR, JA, HE, AU, DV, BB, WAL, LBu, AM, CBe, SH and LBe; Statistical analysis:, LG, JDo, HV and JA; Obtained funding: SH, CBe and LBe. All the authors have read the manuscript and approved its submission. YA help us to design and generate skNAC KO mice.

