## Supplemental data for "skNAC is a Key Driver of Cardiomyocyte Integrity Against Pathological Cardiac Hypertrophy and Heart Failure"

#### Supplemental Methods

All data, methods and materials from this study are available from the corresponding author upon reasonable request.

##### Reagents and chemicals

Phenylephrine (PE) was purchased from Biotechne (#2838). Tamoxifen (#5648), Iscove's Modified Dulbecco's Medium (IMDM) (#I3390), Penicillin-Streptomycin (PS) (#15070063), Amphotericin B (AmB) (#15290026), Duolink In Situ Detection Reagents Orange (#DUO92007), Angiotensin II (AngII) human (#05-23-0101-25MG), Gelatin solution (#G1393), Milk (#70166), Polyvinylidene Difluoride Transfer Membranes (PVDF) (#IPVH00010) and Laminin (#L2020) were obtained from Sigma-Aldrich. [$^{14}$C]-Phenylalanine (#NEC284E050UC) was purchased from Perkin Elmer. Collagenase type II (#LS004177) was purchased from Worthington. Rhodamine labeled wheat germ agglutinin (WGA) (#RL1022) was obtained from Vector Biolab. Ketamine (Nimatek) was obtained from Dechra. Xylazine (Rompum) was purchased from Bayer. Trypsin (1:250) (#27250-018), Collagenase type II (#17101-015), Hanks' Balanced Salt Solution (HBSS) (#14170-112) and Hoechst (#62249) were obtained from Thermo Fisher Scientific. BM chemiluminescence blotting system (#11500 694001) was obtained from Roche. Bovine serum albumin (BSA) for blocking solution (#3854.4) and phosphomolybdic acid (#4440.3) were purchased from Roth. Dako fluorescence mounting medium (#S3023) was obtained from Agilent. Lipofectamine 3000 (#L3000-008), Lipofectamine RNAiMAX (#13778150) and Medium 199 (M199) (#31150-022) were purchased from Invitrogen. Control non-targeting siRNA (ON-TARGETplus Non-targeting siRNA) (#D-001810-01) and Custom siRNA skNAC (Standard 0.025 µmol ON-TARGET plus) were obtained from Dharmacon. RNAeasy mini kit (#74106) and RNase-Free DNAse set (#79256) were from Qiagen. The iScript cDNA synthesis kit (#1708891) was obtained from Bio-Rad. The qPCR core kit for SYBR Green I (#RTSN1005NR) was purchased from Eurogentech. All antibodies, primers, siRNA mentioned in this study are listed in **Supplemental Tables 8 and 9**.

#### Generation of the polyclonal skNAC antibody

The production of a polyclonal antibody targeting both rat and mouse skNAC protein sequences was carried out in collaboration with the Belgian manufacturer Eurogentec. Immunization was performed in two rabbits following a standard 87-day protocol, which included four peptide injections (200 µg each) on days 0, 14, 28, and 56. The immunogenic peptide used was a combination of two sequences (VTSPSPQKTPKSVS and APAPSAKQPVLKNNK) located in the internal region of exon 3, identical to the rat and mouse protein sequences. KLH was used as the carrier protein. Eurogentec conducted an ELISA test to assess adequate skNAC antibody production, and 5 mL of serum was purified using antigen-specific IgG affinity purification. The specificity of the skNAC antibody was successfully validated by western blot and immunofluorescence staining, using siRNA-mediated skNAC knockdown in neonatal rat ventricular myocytes (NRVMs).

#### Generation of skNAC$^{flx/flx}$ mice

skNAC$^{flx/flx}$ mice were generated by introducing loxP sites flanking exon 3 of the skNAC isoform using the CRISPR/Cas9 system. Guide RNAs (crRNAs) targeting intronic regions upstream and downstream of exon 3 were designed using the CRISPRdirect tool [1]. Left crRNA: 5’-GAGAATGACGGTCACCACAC-3’; right crRNA: 5’-GAGTTTGGGCATAGTACATA-3’). Homology-directed repair (HDR) templates were synthesized as two skNAC ultramer oligonucleotides containing the respective loxP sequences flanked by homology arms. Left:
5’AATCACCGAACTATGCACAGGCAAAGTGGATATTGCTTGTCACCTGTTTCTAATCCCGT
GGCTAGCATAACTTCGTATAATGTATGCTATACGAAGTTATTGGTGACCGTCATTCTCAG
ATTTTAAACTTTGTTGTTTCCTGTGAGGCATGTTACCAACT3’; right:
5’CAATACTAACTAGGACGCGTCGCTTTGTACAGTATGTCTGCTTGAGTTTGGGCATAGT
ACGAATTCATAACTTCGTATAATGTATGCtTATACGAAGTTATATAAGGTGGTAACTTTAA
ATGTAGAAGCTCTAGAAATATGATCTAAATCTTGGTTTCTGT3’). Annealed crRNA/tracrRNA (2.4 pmol/µL), Cas9 nuclease V3 (200 ng/µL) and both DNA donor templates (5 ng/µL each) were mixed in IDTE buffer and injected into the pronucleus of B6D2F2 mouse

zygotes. Injected zygotes were incubated in KSOM medium at 37°C for 2 hours before being transferred into oviducts of CD1 pseudo-pregnant female mice. Identification of the positive founders with insertion of both loxP sites was performed by PCR genotyping and confirmed by sequencing. Positive founders were bred with wild-type C57BL/6N mice to obtain heterozygotes floxed offspring (skNAC$^{flx/+}$). These animals were then backrossed to seven generations (N7) onto C57BL/6N background to ensure a homogeneous strain. Finally, (skNAC$^{flx/+}$) mice were intercrossed to generate homozygous floxed mice (skNAC$^{flx/flx}$). crRNAs, Ultramers, tracrRNA (1072534), Cas9 nuclease V3 (1081059) and IDTE buffer (11-01-02-02) were obtained from integrated DNA Technologies[1].

**Generation of cardiac-specific skNAC deficient mice**

Two cardiac-specific skNAC knockout (KO) mouse models, a constitutive (cKO) and an inducible (iKO) deficient mouse model, were generated using commercially available Cre-expressing mouse strains, that had been previously backcrossed onto C57BL/6N background. skNAC$^{flx/flx}$ animals were crossed with αMyHC-Cre$^{+/-}$ transgenic mice (Jackson Laboratory, stock #011038) to generate cKO mice. This breeding produced skNAC$^{-/-}$/αMyHC-Cre$^{+/-}$ mice. Littermate αMyHC-Cre$^{+/-}$ mice were used as controls. Next, skNAC$^{flx/flx}$ mice were bred with αMHC-MerCreMer$^{+/-}$ transgenic mice (Jackson Laboratory, stock #005657) to generate the iKO model, skNAC$^{flx/flx}$ /MerCreMer$^{+/-}$ mice. Tamoxifen (20 mg/kg, intraperitoneal injection) was administered daily for five consecutive days to induce recombination and deletion. Mice were used for experiments two weeks after the final injection. Littermate skNAC$^{flx/flx}$/MerCreMer$^{+/-}$ mice treated with olive oil (vehicle) served as controls.

**Genome-wide association study-based identification and mapping of *NACA*-associated SNPs**

We conducted a comprehensive analysis of the GWAS Catalog (https://www.ebi.ac.uk/gwas) to investigate potential associations between the *NACA* gene and cardiovascular diseases. Initially, we retrieved all single nucleotide polymorphisms (SNPs) and their associated genes from the catalog. We then classified and grouped the corresponding trait annotations for each SNP using the Experimental Factor Ontology (EFO) system. This dataset was filtered to identify GWAS entries with SNPs located within the *NACA* gene, and these entries were subsequently organized according to their EFO trait categories. To visualize the genomic distribution of these *NACA*-associated SNPs, we retrieved the genomic structure from the University of California Santa Cruz (UCSC) Genome Browser Database. The genomic coordinates of the identified SNPs were aligned to the *NACA* gene model, using the Gviz Bioconductor package. The resulting plots displayed the SNPs positions relative to the *NACA* exon-intron boundaries and transcriptional orientation, with each SNP color-coded according to its EFO trait classification.

**Isoform-level expression analysis of skNAC and αNAC using public RNA-seq Datasets**

To investigate the isoform-specific expression of skNAC and αNAC, we reanalyzed publicly available bulk RNA-sequencing datasets from the Gene Expression Omnibus (GEO) under accession numbers GSE101977 [2]. Raw FASTQ files were downloaded using SRA Toolkit (v3.0.7) and aligned to the Mus musculus GRCm39 reference genome using STAR (v2.7.1a) for exon-level quantification and Salmon (v1.10.3) for transcript-level quantification. The resulting exon and transcript count matrices were imported into R (v4.2.0), and normalized expression values were calculated as Counts Per Million (CPM) using the edgeR package (v3.38.4). Specifically, the CPM values for exon 3 derived from the STAR alignment, were used as a proxy for skNAC isoform expression, since exon 3 is uniquely present in the skNAC transcript.

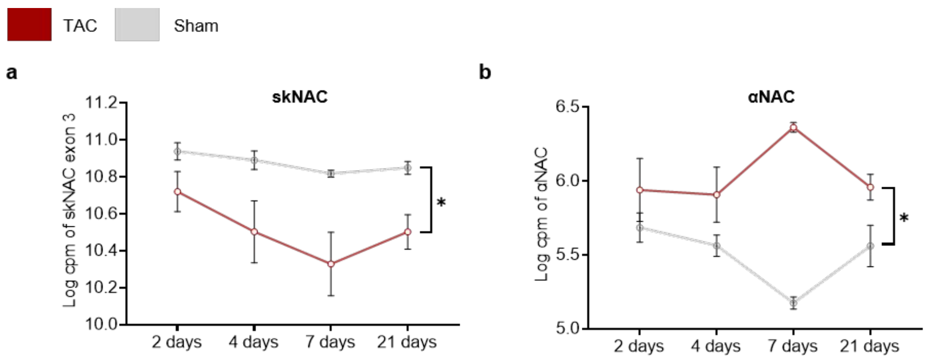

**Figure S1: Differential regulation of skNAC and αNAC isoforms during cardiac hypertrophy.**

**a-b,** Time-course (2-, 4-, 7- and 21-days post-surgery) analysis of skNAC **(a)** and αNAC **(b)** mRNA expression in mouse heart following transverse aortic constriction (TAC) or Sham surgery (N=3). Quantification is performed from publicly bulk RNA-seq datasets (GSE101977). Expression levels are shown as log2 counts per million (log cpm). Data are represented ± SEM. Statistical analyses are performed two-way ANOVA followed by Bonferroni post hoc test. *p<0.05.

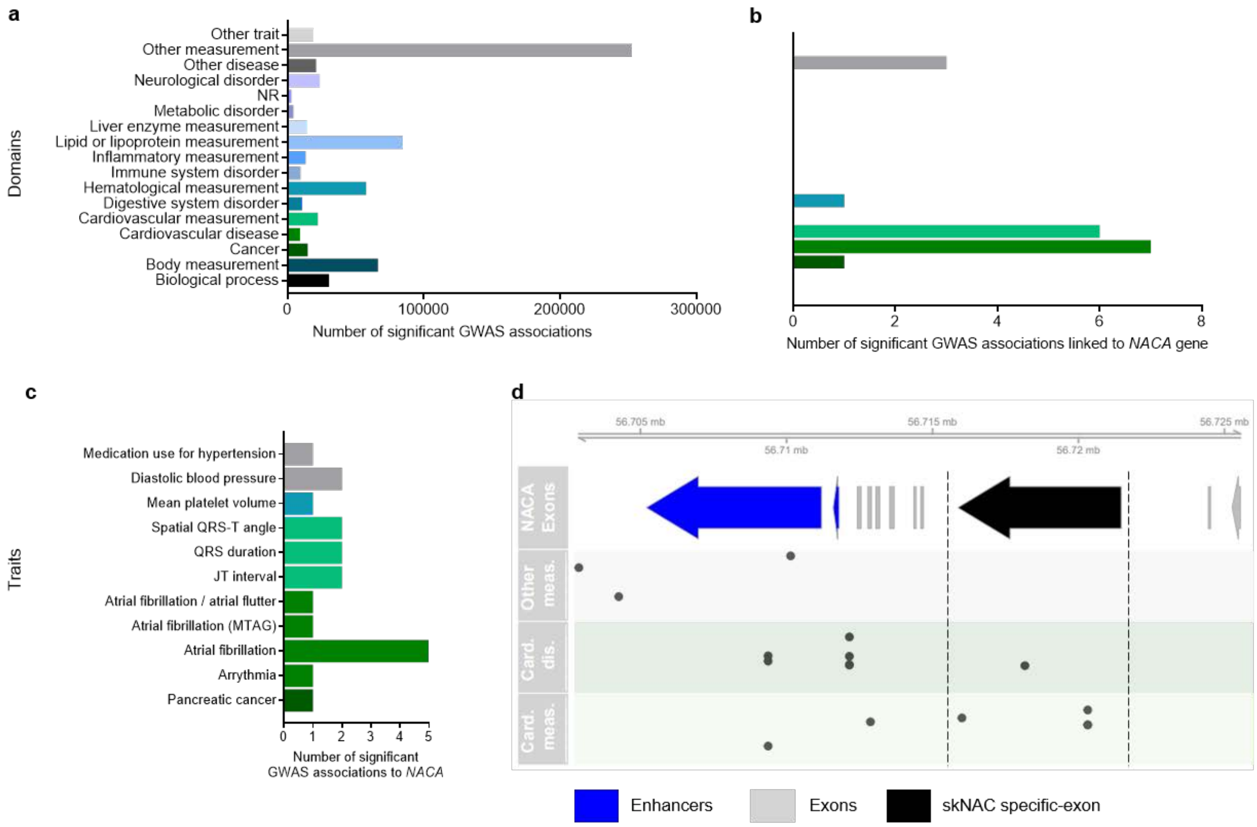

**Figure S2: Variants of *NACA* gene are associated with cardiovascular traits.**

**a,** Distribution of Experimental Factor Ontology (EFO) terms across all significant trait-SNP associations in the GWAS Catalog**. b,** Distribution of EFO terms for significant associations involving SNPs mapped to the *NACA* gene. **c,** Distribution of disease-related traits for significant associations involving SNPs mapped to the *NACA* gene. **d,** Genomic mapping of SNPs associated with the *NACA* gene stratified by their EFO term categories.

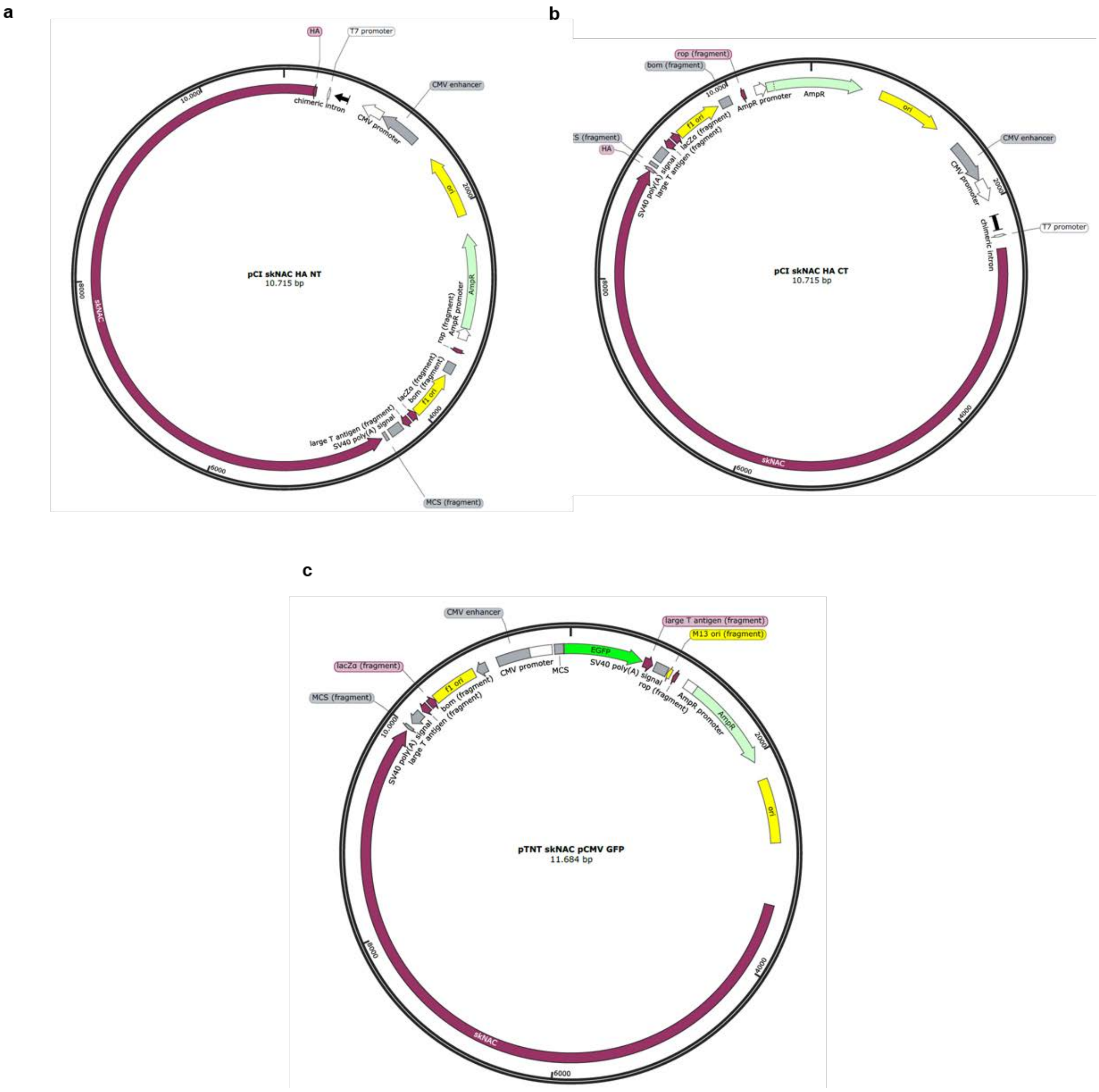

**Figure S3: skNAC expressing vectors used in the study.**

**a-c,** Schematic representation of the plasmid constructs used for skNAC overexpression, generated using SnapGene software version 8.2. N-terminus HA-tagged skNAC (NT-HA) plasmid map **(a)**, C-terminus HA-tagged skNAC (CT-HA) plasmid map **(b)** and skNAC/GFP plasmid map **(c)**.

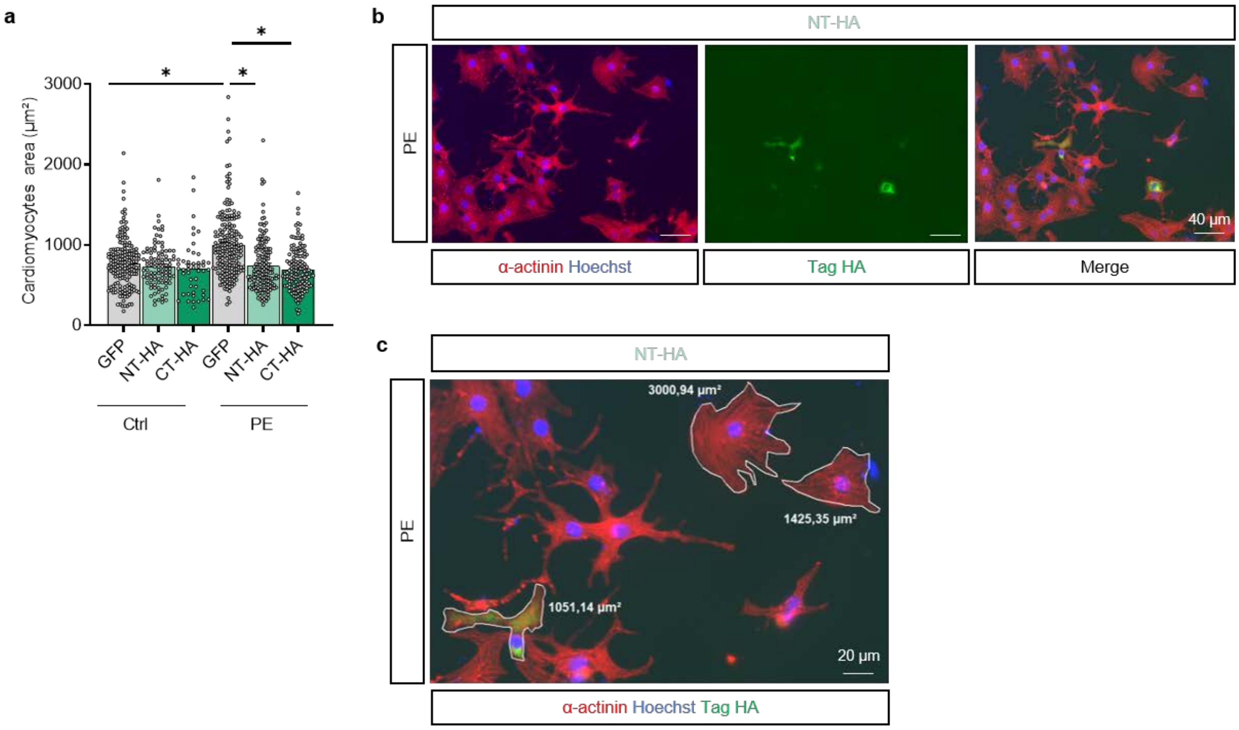

**Figure S4: skNAC overexpression prevented NRVMs hypertrophy development.**

**a-c,** Representative experiment of the study of skNAC overexpression on phenylephrine (PE)-induced NRVMs hypertrophy. NRVMs are treated or not with PE (20µM) for 24h and transfected with GFP (for control), skNAC NT-HA tag plasmid or skNAC CT-HA tag plasmids for 72h. Quantification of NRVMs area (µm²) analyzed after α-actinin immunostaining **(a)**. Each bar represents an average of 100–200 transfected cells analyzed per condition, points representing transfected cells. Data are represented ± SEM. Statistical analyses are performed using two-way ANOVA followed Bonferroni post hoc test. Representative immunostaining images of α-actinin (red) and NT-HA tag (green) **(b)**. Example of cell surface area quantification of transfected and untransfected cells (the latter not being taken into account for **(a)** in a same dish **(c)**. Scale bars, 40 µm **(b)** or 20 µm **(c)**.

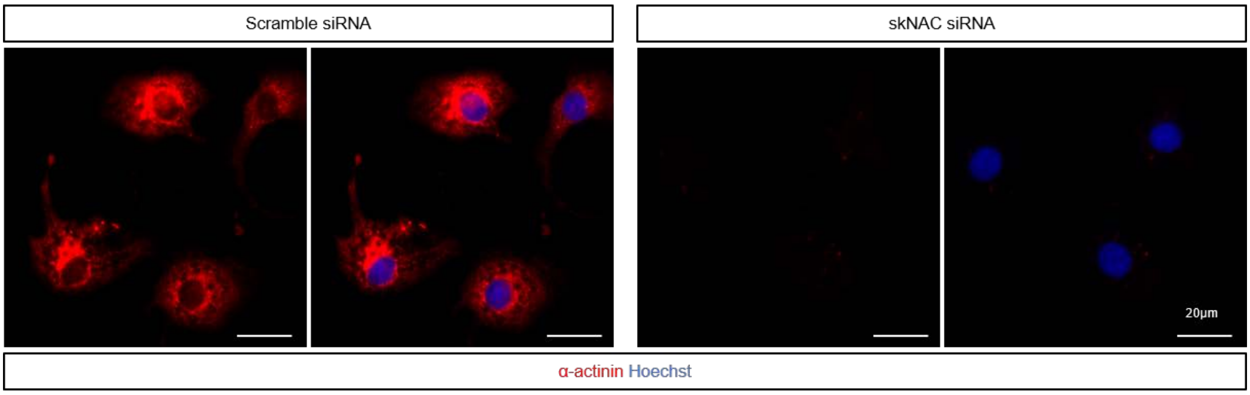

**Figure S5: Cytoplasmic localisation of skNAC in NRVMs.**

Representative images of skNAC immunofluorescence staining (red) in control (Scramble) and siRNA-mediated skNAC knockdown NRVMs, Scale bar, 20µm. Hoechst is used to stain nucleus (blue).

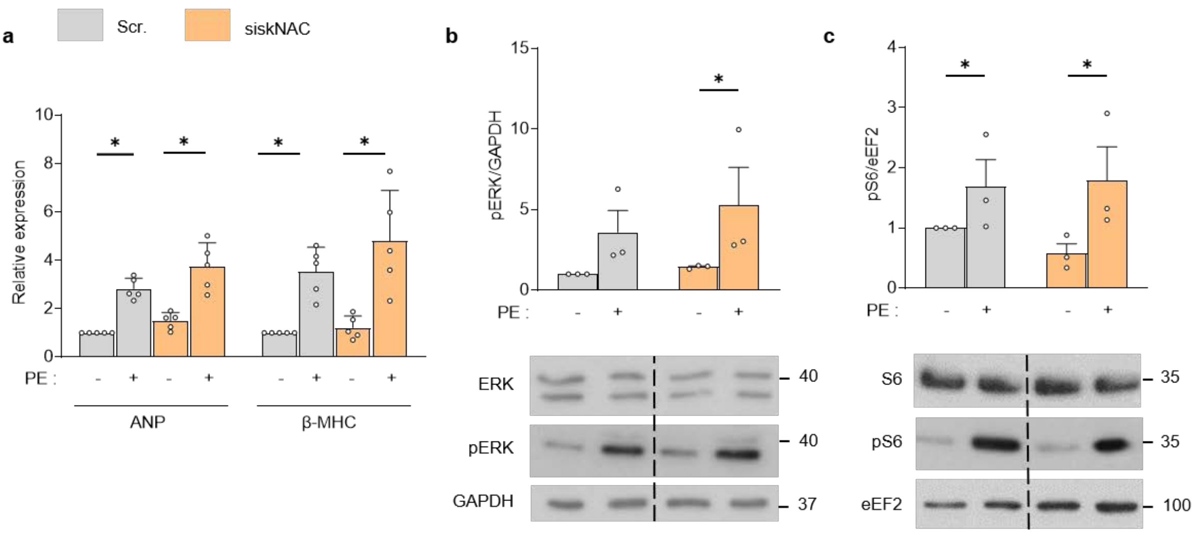

**Figure S6: siRNA-mediated skNAC silencing is not associated with increase in classical hypertrophic markers.**

**a-c,** Effect of siRNA-induced skNAC knockdown on ANP and β-MHC expression, as well as on ERK and S6 phosphorylation state. NRVMs are treated with control non-targeting siRNA (Scramble, Scr.) or skNAC siRNA (siskNAC) for 72h and treated or not with phenylephrine (PE, 20μM) for 24h. ANP and β-MHC mRNA level, normalised to *Rpl32* (N=5) **(a)**. **(b, c)** Immunoblots and quantification of ERK **(b)** and S6 **(c)** phosphorylation state, normalised to GAPDH and eEF2, respectively (N=3). Each point represents an independent biological replicate. Data are represented ± SEM. Statistical analyses are performed using two-way ANOVA followed Bonferroni post hoc test. *$p<0.05$ vs. corresponding untreated cells.

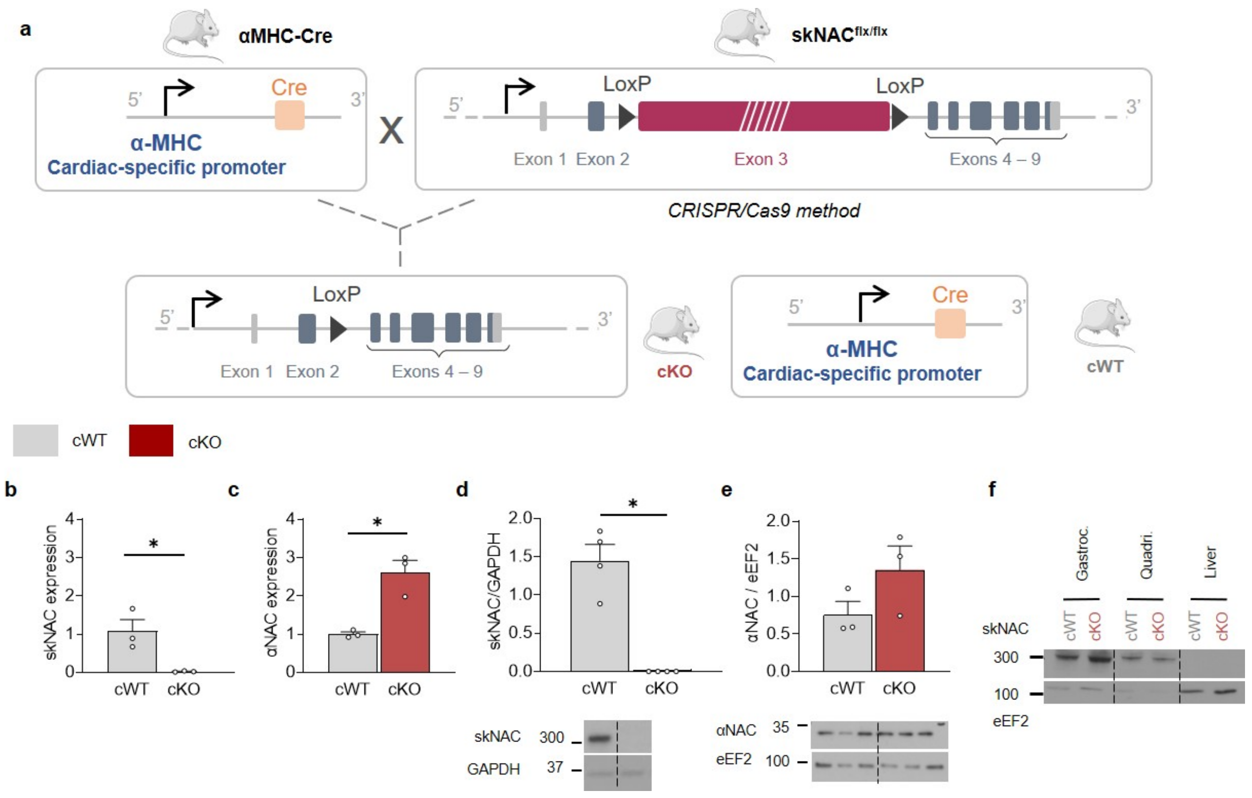

**Figure S7: Constitutive cardiac-specific skNAC deficient mouse model.**

**a,** Schematic representation of the constitutive cardiac-specific skNAC deficient mouse skNAC$^{-/-}$/αMyHC-Cre$^{+/-}$ (cKO). αMyHC-Cre$^{+/-}$ (αMHC-Cre) mice crossed with skNAC$^{flx/flx}$ mice allow constitutive cardiac-specific deletion of skNAC isoform. Control mice (cWT) correspond to αMHC-Cre $^{+/-}$ mice. **b-f,** Validation of the cKO model. skNAC **(b)** and αNAC **(c)** mRNA level in cWT and cKO mice, normalised to *Rpl32* (N=3). skNAC **(d)** and αNAC **(e)** protein level in cWT and cKO mice, normalised to GAPDH and eEF2, respectively (N=4-3). skNAC protein level in other tissues, skeletal muscle gastrocnemius (gastroc.), quadriceps (quadri.) and liver (as negative control) to validate cardiac-specific deletion **(f)**. Each point represents an animal. Data are represented ± SEM. Statistical analyses are performed using two-way ANOVA followed Bonferroni post hoc test. *$p<0.05$ vs. cWT mice.

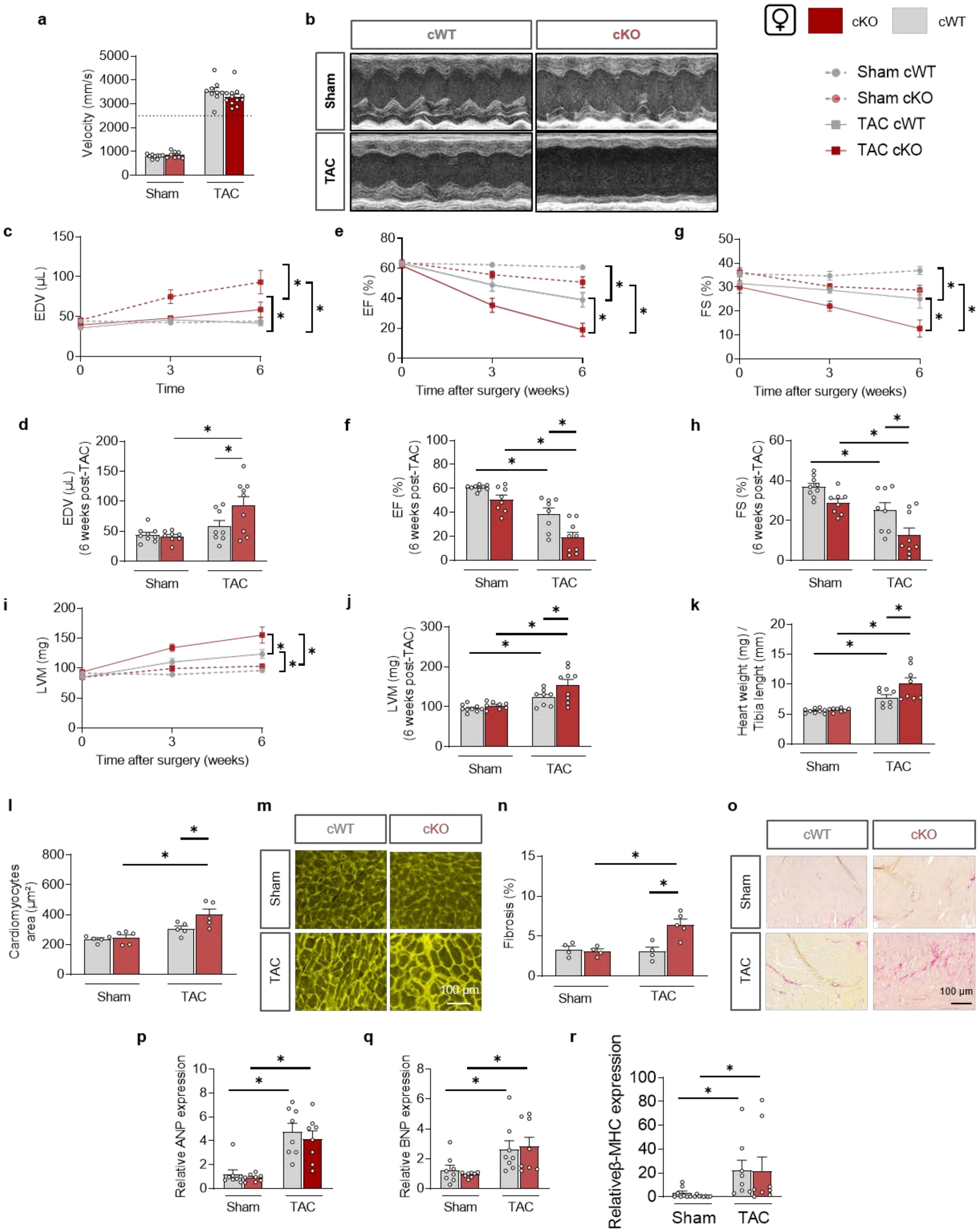

**Figure S8: Constitutive cardiomyocyte-specific skNAC deficiency exacerbates TAC-induced heart failure in females.**

**a-j,** Echocardiographic analysis of sham- and transverse aortic constriction (TAC)-operated cardiomyocyte-specific skNAC knockout (cKO) and control littermate (cWT) female mice at baseline and 3- and 6-weeks post-surgery (N=8-13). Quantification of peak aortic velocity at the site of constriction in sham- and TAC-operated cWT and cKO female mice **(a)**. Representative image of M-mode in sham- and TAC-operated cWT and cKO female mice at 6 weeks post-surgery **(b)**. End-diastolic volume (EDV) **(c, d)**, ejection fraction (EF) **(e, f)** and fractional shortening (FS) **(g, h)** evaluated by echocardiography over time and at end-time point (6 weeks post-surgery). Cardiac hypertrophy (Left ventricular mass, LVM) evaluated by echocardiography over time **(I)** and at end-time point (6 weeks post-surgery) **(j)**, and at time of sacrifice by the heart weight to tibia length ratio **(k)**. **l-o,** Histological analysis of sham- and TAC-operated cWT and cKO female mice (N=5). Quantification of cross-sectional cardiomyocyte area **(l)** and representative images **(m)** of wheat-germ agglutinin immunofluorescence staining, at 6 weeks post-surgery. Scale bar, 100µm. Quantification of fibrosis **(n)** and representative images **(o)** of collagen staining with Picrosirius red, at 6 weeks post-surgery. Scale bar, 100µm. **p-r,** ANP **(p)**, BNP **(q)** and β-MHC **(r)** mRNA level in heart mice, normalised to *Rpl32* (N=8-14). Each point represents an independent animal. Values are expressed as mean ± SEM. Statistical analyses are performed using two-way ANOVA followed by Bonferroni post hoc test. *$p<0.05$.

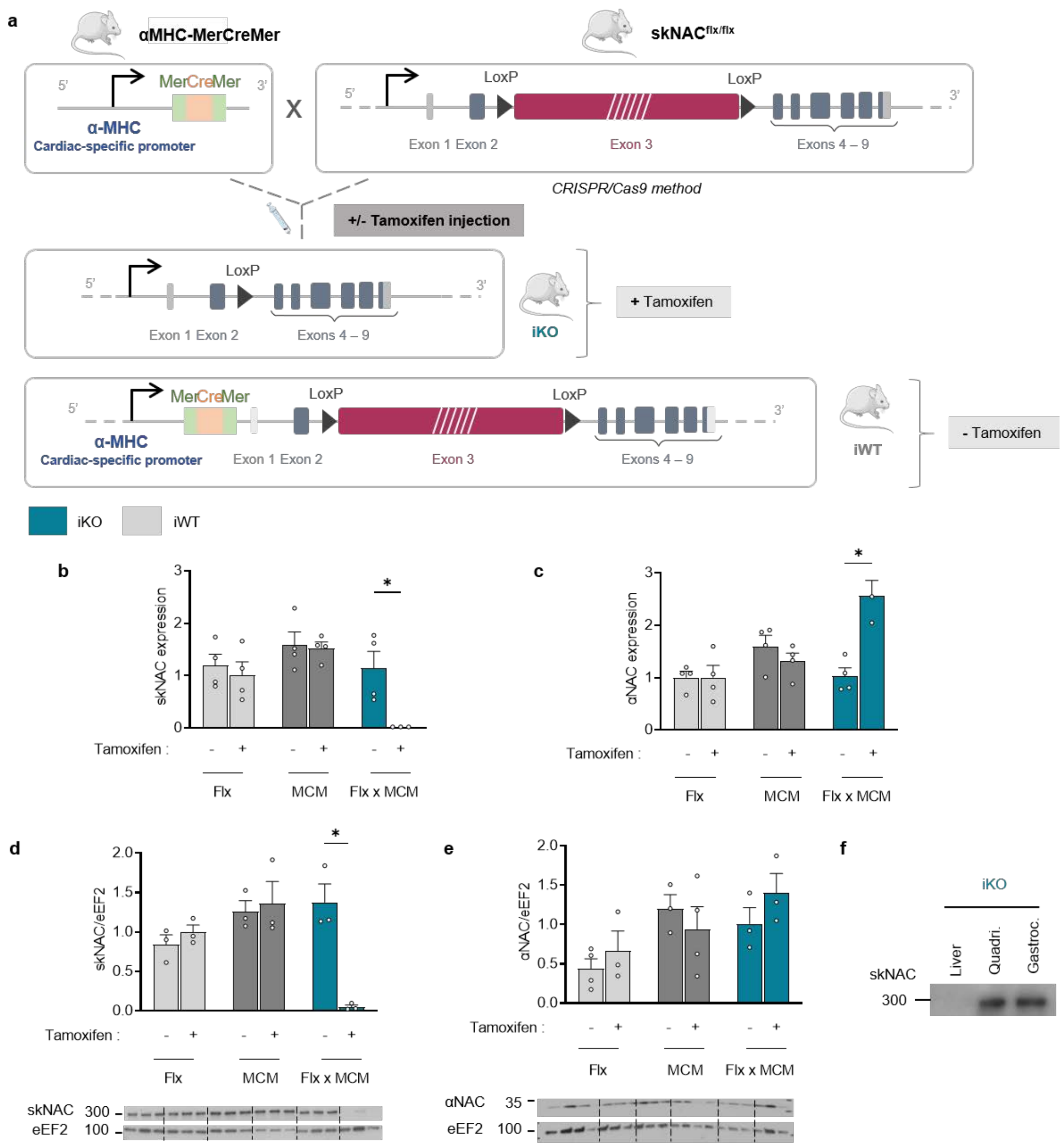

**Figure S9: Inducible cardiac-specific skNAC deficient mouse model.**

**a,** Schematic representation of the inducible cardiac-specific skNAC deficient mouse skNAC$^{-/-}$/MerCreMe$^{+/-}$ (iKO). MerCreMer$^{+/-}$ (αMHC-MerCreMer) mice crossed with skNAC$^{flx/flx}$ mice allow inducible cardiac-specific deletion of skNAC isoform, under Tamoxifen treatment. Tamoxifen (20 mg/kg) is injected intraperitoneally in 2 months-old mice. Tamoxifen is injected in mice once a day for five days. Control mice (iWT) correspond to skNAC$^{flx/flx}$/MerCreMe $^{+/-}$ mice injected with olive oil. **b-f,** Validation of the cKO model. skNAC **(b)** and αNAC **(c)** mRNA level in iWT and iKO mice, normalised to *Rpl32* (N=3-4). skNAC **(d)** and αNAC **(e)** protein level in iWT and iKO mice, normalised to eEF2 (N=3). skNAC protein level in other tissues, skeletal muscle gastrocnemius (gastroc.), quadriceps (quadri.) and liver (as negative control) to validate cardiac-specific deletion **(f)**. Each point represents an animal. Data are represented ± SEM. Statistical analyses are performed using unpaired student t-test. **p<0.05* vs. iWT mice.

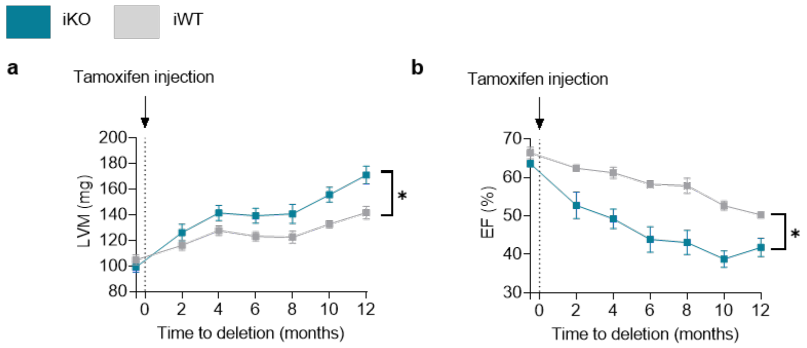

**Figure S10: Inducible cardiac-specific skNAC deficient mice are characterised by progressive basal cardiac hypertrophy and heart failure.**

**a-b,** Longitudinal echocardiographic evaluation of left ventricular mass (LVM) **(a)** and ejection fraction (EF) **(b)** of iWT and iKO mice (N=11-12). Each point represents an independent animal. Values are expressed as mean ± SEM. Statistical analyses are performed using a two-way ANOVA followed Bonferroni post hoc test. **p<0.05* vs. iKO mice.

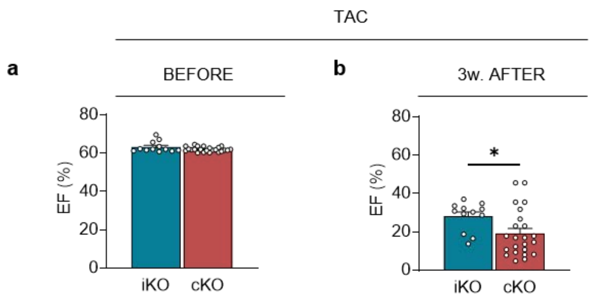

**Figure S11: Comparison of iKO and cKO cardiac systolic function under pressure-overload condition.**

**a-b,** Echocardiographic evaluation of ejection fraction (EF) before transverse aortic constriction (TAC) surgery **(a)** and at 3-week timing post-TAC **(b)** in iKO and cKO mice. Each point represents an independent animal. Values are expressed as mean ± SEM. Statistical analyses are performed unpaired student t-test. *$p<0.05$.

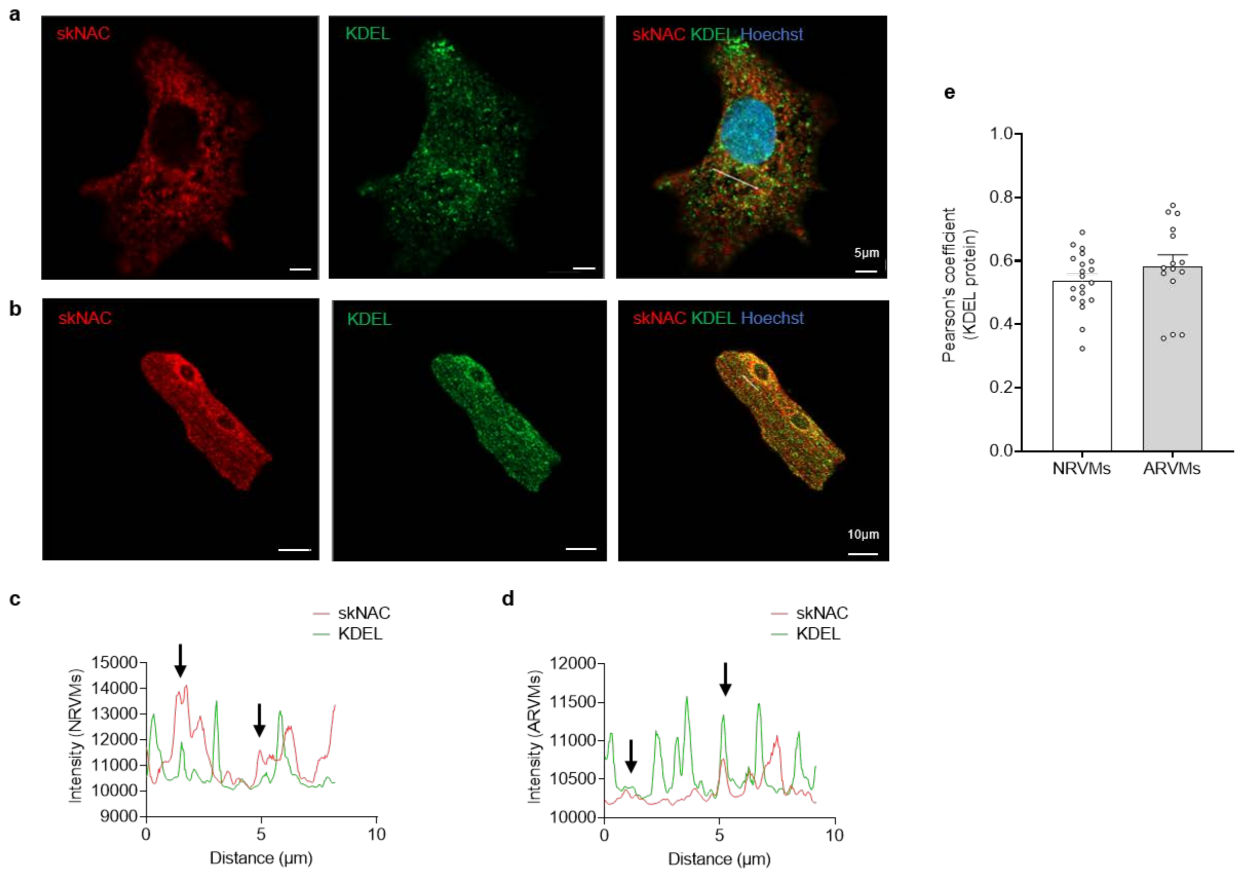

**Figure S12: skNAC partially co-localises with endoplasmic reticulum.**

**a-b,** Confocal imaging of immunofluorescence staining in cultured cardiomyocytes. Representative immunostaining images of skNAC (red), KDEL signal peptide (green), and both merged in the right panel, in NRVMs **(a)** and ARVMs **(b).** Nuclei are stained with Hoechst (blue). Scale bar, 5µm (NRVMs) and 10µm (ARVMs), respectively. **c-d,** Line-scan profiles showing overlapping fluorescence intensity of skNAC (red) with KDEL (green) in NRVMs **(c)** and ARVMs **(d)**. **e,** Quantification of skNAC/S6 co-localisation using Pearson's correlation coefficient in NRVMs and ARVMs (N=15-20). Each dot in the graphic represents an independent biological cell (N=6-20). Data are presented as mean ± SEM.

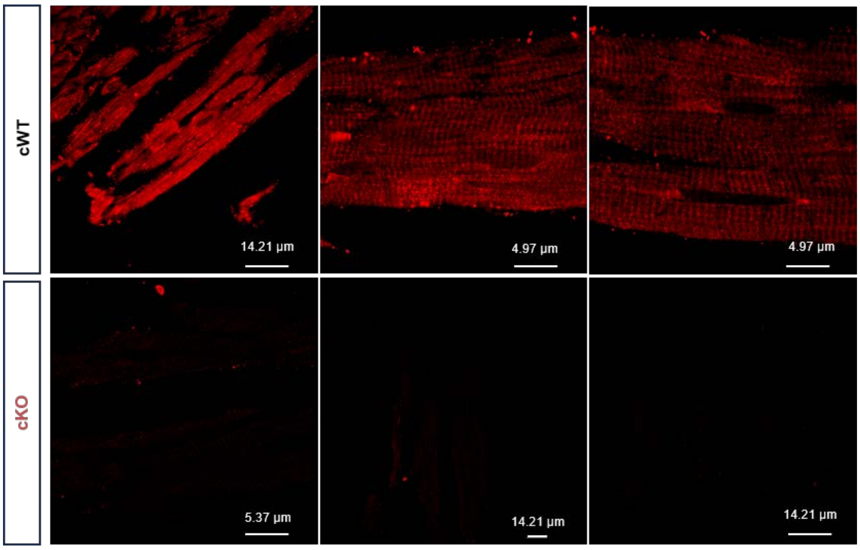

**Figure S13: skNAC immunostaining reveals a sarcomeric pattern in cWT mice which is absent in cKO mice.**

Additional immunostaining images of skNAC (red) in cWT and cKO mouse heart sections.

a

b

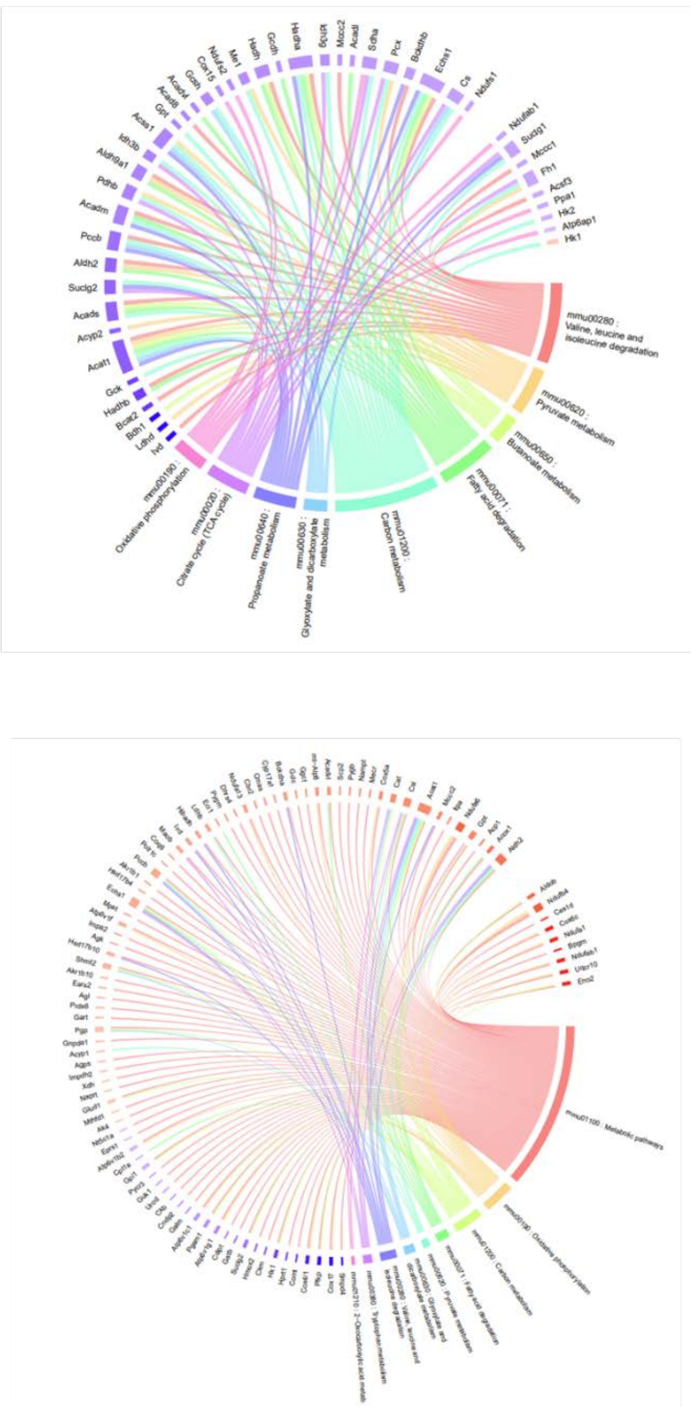

**Figure S14: Chord diagrams of KEGG pathways in iKO mice from transcriptomic and proteomic data.**

**a-b,** Chord diagram representing the relationships between modulated genes and enriched GSEA pathways in transcriptomic **(a)** and proteomic **(b)** analysis. Genes are connected to their associated KEGG pathways through colored links. Gene modulation is represented by color according to the −log2 (FC) in iKO compared to iWT, ranging from low (blue) to high (red). Genes included in the diagram were selected using an FDR threshold of 0.3.

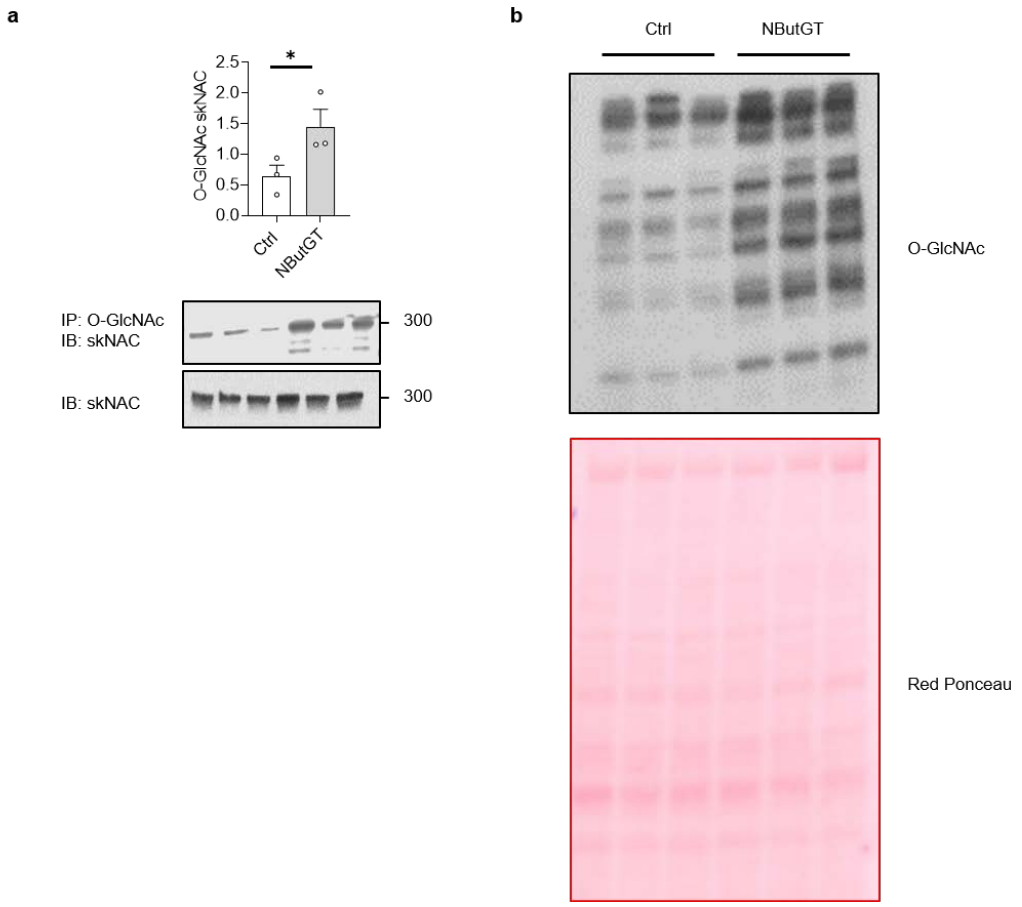

**Figure S15: Cardiac skNAC can be O-GlcNAcylated.**

Mice were treated with NButGT (50 mg/kg, intraperitoneal) or vehicle (NaCl, 0.9%) for 6 hours. **a,** Relative O-GlcNAcylated skNAC level in control (Ctrl) and NButGT-treated mice. Representative immunoblots of total skNAC and O-GlcNAcylated skNAC (after immunoprecipitation of O-GlcNAcylated proteins and immunoblot anti-skNAC) (N=3). **b** Immunoblots of total cardiac protein O-GlcNAcylation in the same samples, ponceau staining being used as loading control (N=3). Each point represents an independent animal. Values are expressed as mean ± SEM. Statistical analyses are performed using unpaired student t-test. *p<0.05. vs. corresponding untreated Ctrl mice.

**Supplemental Tables S1–S9**

**Supplementary Table S1: Clinical characteristics and echocardiographic parameters of the human cohort.**

| **Characteristics** | | **Mean ± SEM** |
|---|---|---|
| **Age** | | 69.12 ± 2.66 |
| **Sex (males) (%)** | | 70 |
| **Size (cm)** | | 168.05 ± 2.33 |
| **Weight (kg)** | | 73.63 ± 3.03 |
| **Log NT-proBNP** | | 2.30 ± 0.17 |
| **BMI (kg/m²)** | | 43.66 ± 1.58 |
| Echocardiographie parameters | | |
| | **EF (%)** | 71.77 ± 2.60 |
| | **EDV (mL)** | 105.79 ± 8.18 |
| | **ESV (mL)** | 30.30 ± 3.81 |
| | **IVSd (cm)** | 1.23 ± 0.07 |
| | **LVDd (cm)** | 2.69 ± 0.15 |
| | **LVDs (cm)** | 2.73 ± 0.14 |
| | **PWd (cm)** | 1.13 ± 0.05 |
| | **LVM (g)** | 212.42 ± 12.37 |

Values are expressed as mean ± SEM. BMI, Body mass Index (weight (kg)/size($m^2$)); EDV, Ejection Diastolic Volume; EF, Ejection Fraction; ESV, Ejection Systolic Volume; IVSd, Interventricular Septum Thickness at end Diastole; LVDd, Left-Ventricular Diameter in diastole; LVDs, Left-Ventricular Diameter in systole; LVM, Left Ventricular Mass. PWd, Posterior Wall Thickness in diastole.

**Supplementary Table S2: Echocardiographic parameters of cWT and cKO females performed every 2 months.**

| | Time | cWT | cKO |
|---|---|---|---|
| **LVM (mg)** | 2 months | 76.94 ± 4.95 | 90.03 ± 3.72* |
| | 4 months | 80.94 ± 4.86 | 107.43 ± 2.59* |
| | 6 months | 97.28 ± 5.44 | 126.80 ± 7.76* |
| | 8 months | 85.66 ± 1.02 | 154.63 ± 17.21* |
| **EDV (µL)** | 2 months | 37.68 ± 2.75 | 46.00 ± 2.20* |
| | 4 months | 40.25 ± 2.70 | 52.54 ± 2.19* |
| | 6 months | 48.16 ± 1.27 | 98.07 ± 10.13* |
| | 8 months | 47.87 ± 1.35 | 124.37 ± 27.02* |
| **ESV (µL)** | 2 months | 14.74 ± 1.18 | 21.04 ± 1.33* |
| | 4 months | 16.10 ± 1.32 | 26.52 ± 1.46* |
| | 6 months | 23.91 ± 1.89 | 80.49 ± 11.11* |
| | 8 months | 20.72 ± 0.93 | 107.92 ± 26.96* |
| **EF (%)** | 2 months | 61.03 ± 0.75 | 54.60 ± 1.37* |
| | 4 months | 60.20 ± 1.23 | 49.95 ± 1.83* |
| | 6 months | 51.83 ± 3.74 | 20.77 ± 4.79* |
| | 8 months | 56.78 ± 1.03 | 17.17 ± 6.38* |
| **HR (bpm)** | 2 months | 517.67 ± 11.41 | 512.48 ± 10.59 |
| | 4 months | 520.44 ± 14.11 | 547.03 ± 18.67 |
| | 6 months | 514.44 ± 10.96 | 529.36 ± 7.35 |
| | 8 months | 508.88 ± 20.14 | 511.52 ± 5.78 |
| **CO (mL/min)** | 2 months | 12.39 ± 1.18 | 12.00 ± 0.53 |
| | 4 months | 12.61 ± 0.93 | 12.51 ± 0.76 |
| | 6 months | 13.05 ± 3.08 | 9.48 ± 1.69 |
| | 8 months | 13.94 ± 0.65 | 10.03 ± 1.97* |
| **FS (%)** | 2 months | 36.64 ± 1.72 | 33.09 ± 1.23 |
| | 4 months | 37.83 ± 2.71 | 27.74 ± 1.01* |
| | 6 months | 29.84 ± 3.74 | 12.22 ± 2.48* |
| | 8 months | 32.86 ± 3.25 | 7.61 ± 4.70* |
| **IVSd (mm)** | 2 months | 0.70 ± 0.05 | 0.64 ± 0.20 |
| | 4 months | 0.66 ± 0.01 | 0.71 ± 0.02 |
| | 6 months | 0.69 ± 0.03 | 0.69 ± 0.04 |
| | 8 months | 0.71 ± 0.02 | 0.74 ± 0.04 |
| **IVSs (mm)** | 2 months | 1.02 ± 0.03 | 1.00 ± 0.02 |
| | 4 months | 1.05 ± 0.04 | 1.04 ± 0.02 |
| | 6 months | 1.02 ± 0.03 | 0.83 ± 0.06* |
| | 8 months | 1.07 ± 0.04 | 0.83 ± 0.04* |
| **LVIDd (mm)** | 2 months | 3.12 ± 0.07 | 3.46 ± 0.05* |
| | 4 months | 3.27 ± 0.10 | 3.41 ± 0.05 |
| | 6 months | 3.43 ± 0.11 | 4.55 ± 0.24* |

| | | | |
|---|---|---|---|
| | 8 months | 3.45 ± 0.10 | 5.00 ± 0.50* |
| **LVIDs (mm)** | 2 months | 1.98 ± 0.07 | 2.32 ± 0.06* |
| | 4 months | 2.04 ± 0.12 | 2.47 ± 0.06* |
| | 6 months | 2.42 ± 0.18 | 4.04 ± 0.31* |
| | 8 months | 2.33 ± 0.15 | 4.69 ± 0.66* |
| **LVPWd (mm)** | 2 months | 0.63 ± 0.03 | 0.63 ± 0.01 |
| | 4 months | 0.64 ± 0.02 | 0.70 ± 0.01 |
| | 6 months | 0.68 ± 0.02 | 0.72 ± 0.05 |
| | 8 months | 0.77 ± 0.02 | 0.77 ± 0.05 |
| **LVPWs (mm)** | 2 months | 1.00 ± 0.02 | 0.95 ± 0.03 |
| | 4 months | 0.95 ± 0.04 | 0.99 ± 0.02 |
| | 6 months | 0.91 ± 0.03 | 0.86 ± 0.08 |
| | 8 months | 1.17 ± 0.05 | 0.84 ± 0.06* |

All parameters are expressed as means ± SEM. Statistical analyses were performed t-test student. *$p<0.05$ vs. control mice (cWT). CO, Cardiac Output; EDV, Ejection Diastolic Volume; EF, Ejection Fraction; ESV, Ejection Systolic Volume; FS, Fractional Shortening; HR, Heart Rate; IVSd, Interventricular Septum Thickness at end Diastole, IVSs, Interventricular Septum Thickness at end Systole; LVIDd, Left-Ventricular Internal Diameter in diastole; LVIDs, Left-Ventricular Internal Diameter in systole; LVM, Left Ventricular mass; LVPWd, Left Ventricular Posterior Wall Thickness in diastole; LVPWs, Left Ventricular Posterior Wall Thickness in systole.

**Supplementary Table S3: Echocardiographic parameters of cWT and cKO males performed every 2 months.**

| | Time | cWT | cKO |
|---|---|---|---|
| **LVM (mg)** | 2 months | 105.60 ± 6.44 | 109.05 ± 5.26 |
| | 4 months | 122.35 ± 7.51 | 141.50 ± 5.67 |
| | 6 months | 130.34 ± 4.61 | 156.74 ± 8.42 |
| | 8 months | 169.97 ± 5.63 | 181.96 ± 15.92 |
| **EDV (µL)** | 2 months | 49.33 ± 2.49 | 49.53 ± 2.09 |
| | 4 months | 56.57 ± 3.74 | 55.37 ± 2.40 |
| | 6 months | 87.67 ± 9.00 | 95.52 ± 9.77 |
| | 8 months | 123.04 ± 24.39 | 113.76 ± 15.56 |
| **ESV (µL)** | 2 months | 18.41 ± 0.88 | 20.59 ± 1.02 |
| | 4 months | 23.20 ± 1.37 | 27.64 ± 2.10 |
| | 6 months | 55.76 ± 11.84 | 69.01 ± 11.72 |
| | 8 months | 100.83 ± 26.12 | 94.40 ± 16.56 |
| **EF (%)** | 2 months | 62.62 ± 0.41 | 58.44 ± 0.99* |
| | 4 months | 58.66 ± 1.53 | 49.89 ± 1.78* |
| | 6 months | 39.58 ± 5.77 | 27.59 ± 3.48 |
| | 8 months | 20.60 ± 5.63 | 20.08 ± 4.42 |
| **HR (bpm)** | 2 months | 491.71 ± 13.98 | 508.67 ± 13.60 |
| | 4 months | 505.39 ± 11.21 | 550.20 ± 8.89 |
| | 6 months | 510.89 ± 8.23 | 535.14 ± 8.90 |
| | 8 months | 581.83 ± 7.51 | 523.97 ± 13.02 |
| **CO (mL/min)** | 2 months | 14.12 ± 0.75 | 14.64 ± 0.63 |
| | 4 months | 16.88 ± 1.44 | 13.36 ± 0.45* |
| | 6 months | 16.36 ± 1.81 | 12.97 ± 1.85 |
| | 8 months | 12.98 ± 2.23 | 10.18 ± 1.35 |
| **FS (%)** | 2 months | 33.93 ± 1.10 | 34.15 ± 0.95 |
| | 4 months | 31.76 ± 1.06 | 26.23 ± 0.96* |
| | 6 months | 16.72 ± 2.74 | 12.38 ± 1.24 |
| | 8 months | 8.65 ± 2.76 | 12.51 ± 2.85 |
| **IVSd (mm)** | 2 months | 0.89 ± 0.04 | 0.90 ± 0.03 |
| | 4 months | 0.80 ± 0.03 | 0.93 ± 0.03* |
| | 6 months | 0.80 ± 0.02 | 0.80 ± 0.02 |
| | 8 months | 0.80 ± 0.04 | 0.74 ± 0.02 |
| **IVSs (mm)** | 2 months | 1.32 ± 0.05 | 1.28 ± 0.04 |
| | 4 months | 1.16 ± 0.04 | 1.27 ± 0.04 |
| | 6 months | 1.07 ± 0.04 | 1.10 ± 0.02 |
| | 8 months | 0.95 ± 0.08 | 0.88 ± 0.03 |
| **LVIDd (mm)** | 2 months | 3.53 ± 0.10 | 3.22 ± 0.08* |

| | | | |
|---|---|---|---|
| | 4 months | 3.72 ± 0.07 | 3.32 ± 0.08* |
| | 6 months | 4.57 ± 0.25 | 4.39 ± 0.21 |
| | 8 months | 4.80 ± 0.34 | 4.74 ± 0.31 |
| **LVIDs (mm)** | 2 months | 2.33 ± 0.06 | 2.12 ± 0.07 |
| | 4 months | 2.54 ± 0.05 | 2.46 ± 0.08 |
| | 6 months | 3.84 ± 0.33 | 3.86 ± 0.26 |
| | 8 months | 4.41 ± 0.44 | 4.20 ± 0.40 |
| **LVPWd (mm)** | 2 months | 0.73 ± 0.02 | 0.78 ± 0.02 |
| | 4 months | 0.75 ± 0.03 | 0.82 ± 0.02 |
| | 6 months | 0.78 ± 0.03 | 0.72 ± 0.03 |
| | 8 months | 0.77 ± 0.08 | 0.76 ± 0.03 |
| **LVPWs (mm)** | 2 months | 1.18 ± 0.05 | 1.11 ± 0.04 |
| | 4 months | 1.11 ± 0.05 | 1.12 ± 0.04 |
| | 6 months | 0.95 ± 0.07 | 0.87 ± 0.03 |
| | 8 months | 0.84 ± 0.10 | 0.81 ± 0.06 |

All parameters are expressed as means ± SEM. Statistical analyses were performed t-test student. *$p<0.05$ vs. control mice (cWT). CO, Cardiac Output; EDV, Ejection Diastolic Volume; EF, Ejection Fraction; ESV, Ejection Systolic Volume; FS, Fractional Shortening; HR, Heart Rate; IVSd, Interventricular Septum Thickness at end Diastole, IVSs, Interventricular Septum Thickness at end Systole; LVIDd, Left-Ventricular Internal Diameter in diastole; LVIDs, Left-Ventricular Internal Diameter in systole; LVM, Left Ventricular mass; LVPWd, Left Ventricular Posterior Wall Thickness in diastole; LVPWs, Left Ventricular Posterior Wall Thickness in systole.

**Supplementary Table S4: Echocardiographic parameters of cWT and cKO males at baseline, 1-, 2- and 3 weeks post-surgery.**

| | | cWT | | cKO | |
|---|---|---|---|---|---|
| | **Time** | **Sham** | **TAC** | **Sham** | **TAC** |
| **LVM (mg)** | baseline | 98.15 ± 3.87 | 104.18 ± 3.78 | 103.85 ± 4.43 | 101.54 ± 5.66 |
| | 1 week | 97.50 ± 7.17 | 113.15 ± 6.71 | 98.56 ± 4.42 | 128.19 ± 7.72† |
| | 2 weeks | 93.37 ± 6.22 | 122.04 ± 8.63 | 103.47 ± 5.04 | 133.41 ± 9.44† |
| | 3 weeks | 97.33 ± 3.08 | 130.90 ± 8.05 | 108.51 ± 5.35 | 177.78 ± 9.88*† |
| **EDV (µL)** | baseline | 49.60 ± 2.85 | 56.95 ± 4.13 | 49.07 ± 3.16 | 53.50 ± 2.53 |
| | 1 week | 50.82 ± 5.35 | 52.86 ± 2.89 | 47.01 ± 3.80 | 52.93 ± 4.01 |
| | 2 weeks | 46.55 ± 4.89 | 57.76 ± 5.07 | 50.40 ± 3.20 | 68.30 ± 8.47 |
| | 3 weeks | 48.12 ± 2.05 | 71.08 ± 6.41 | 56.84 ± 4.08 | 102.13 ± 8.72*† |
| **ESV (µL)** | baseline | 18.59 ± 1.23 | 20.95 ± 1.50 | 18.81 ± 1.33 | 20.62 ± 1.01 |
| | 1 week | 19.83 ± 1.93 | 22.98 ± 1.92 | 18.85 ± 1.66 | 30.85 ± 3.77† |
| | 2 weeks | 18.40 ± 1.94 | 30.08 ± 3.27 | 20.83 ± 2.49 | 45.56 ± 8.12† |
| | 3 weeks | 18.42 ± 0.70 | 45.45 ± 6.28 | 29.30 ± 3.03 | 87.38 ± 9.22*† |
| **EF (%)** | baseline | 62.64 ± 0.67 | 63 ± 0.98 | 61.75 ± 0.90 | 61.48 ± 0.38 |
| | 1 week | 60.80 ± 1.04 | 56.73 ± 1.91 | 59.77 ± 1.73 | 42.22 ± 4.30*† |
| | 2 weeks | 60.37 ± 1.83 | 48.11 ± 3.04 | 59.38 ± 3.05 | 36.53 ± 4.56† |
| | 3 weeks | 61.61 ± 0.74 | 38.90 ± 3.78* | 49.78 ± 2.48 | 18.31 ± 3.95*† |
| **HR (bpm)** | baseline | 515.07 ± 12.45 | 528.41 ± 11.96 | 527.62 ± 11.86 | 504.13 ± 8.26 |
| | 1 week | 523.25 ± 18.43 | 594.97 ± 59.83 | 517.62 ± 11.86 | 522.90 ± 9.70 |
| | 2 weeks | 593.31 ± 22.05 | 541.53 ± 10.62 | 545.36 ± 14.78 | 526.79 ± 8.07 |
| | 3 weeks | 546.21 ± 14.12 | 528.20 ± 7.78 | 533.44 ± 11.57 | 537.39 ± 15.86 |
| **CO (mL/min)** | baseline | 15.91 ± 0.80 | 17.26 ± 1.23 | 16.52 ± 0.92 | 15.17 ± 0.86 |
| | 1 week | 16.26 ± 2.09 | 17.97 ± 2.40 | 15.28 ± 1.32 | 11.42 ± 1.12 |
| | 2 weeks | 16.64 ± 1.85 | 13.87 ± 0.98 | 16.15 ± 1.24 | 11.97 ± 0.94 |
| | 3 weeks | 16.18 ± 0.75 | 13.58 ± 0.85* | 15.18 ± 1.15 | 8.10 ± 0.87*† |
| **FS (%)** | baseline | 37.91 ± 2.11 | 34.60 ± 1.28 | 32.17 ± 1.56 | 31.73 ± 1.85 |
| | 1 week | 32.44 ± 1.70 | 26.64 ± 2.93 | 29.93 ± 2.41 | 23.58 ± 2.81 |
| | 2 weeks | 34.38 ± 3.53 | 28.40 ± 2.17 | 30.05 ± 1.52 | 20.90 ± 2.91† |
| | 3 weeks | 34.56 ± 2.20 | 23.96 ± 1.95 | 27.87 ± 1.58 | 10.27 ± 1.20*† |
| **IVSd (mm)** | baseline | 0.72 ± 0.02 | 0.67 ± 0.02 | 0.75 ± 0.02 | 0.73 ± 0.02 |
| | 1 week | 0.74 ± 0.02 | 0.84 ± 0.03 | 0.72 ± 0.02 | 0.87 ± 0.02† |
| | 2 weeks | 0.77 ± 0.05 | 0.86 ± 0.02 | 0.76 ± 0.01 | 0.86 ± 0.04† |
| | 3 weeks | 0.72 ± 0.01 | 0.83 ± 0.03 | 0.75 ± 0.02 | 0.83 ± 0.02† |
| **IVSs (mm)** | baseline | 1.13 ± 0.04 | 1.11 ± 0.02 | 1.13 ± 0.03 | 1.08 ± 0.02 |
| | 1 week | 1.12 ± 0.02 | 1.19 ± 0.03 | 1.10 ± 0.03 | 1.23 ± 0.04† |
| | 2 weeks | 1.15 ± 0.05 | 1.21 ± 0.03 | 1.14 ± 0.03 | 1.17 ± 0.05 |
| | 3 weeks | 1.12 ± 0.03 | 1.10 ± 0.03 | 1.08 ± 0.03 | 0.97 ± 0.04* |

| | | | | | |
|---|---|---|---|---|---|
| **LVIDd (mm)** | baseline | 3.47 ± 0.09 | 3.53 ± 0.12 | 3.45 ± 0.08 | 3.34 ± 0.08 |
| | 1 week | 3.10 ± 0.09 | 3.42 ± 0.12 | 3.51 ± 0.12 | 3.46 ± 0.14 |
| | 2 weeks | 3.33 ± 0.10 | 3.46 ± 0.12 | 3.44 ± 0.11 | 3.76 ± 0.23 |
| | 3 weeks | 3.41 ± 0.07 | 3.95 ± 0.18 | 3.66 ± 0.08 | 4.33 ± 0.17$^{\dagger}$ |
| **LVIDs (mm)** | baseline | 2.16 ± 0.10 | 2.30 ± 0.06 | 2.35 ± 0.09 | 2.29 ± 0.09 |
| | 1 week | 2.09 ± 0.07 | 2.52 ± 0.17 | 2.46 ± 0.13 | 2.66 ± 0.18 |
| | 2 weeks | 2.18 ± 0.12 | 2.49 ± 0.14 | 2.41 ± 0.10 | 3.01 ± 0.28 |
| | 3 weeks | 2.24 ± 0.08 | 3.02 ± 0.19* | 2.6 4± 0.99 | 3.93 ± 0.23*$^{\dagger}$ |
| **LVPWd (mm)** | baseline | 0.66 ± 0.02 | 0.67 ± 0.01 | 0.72 ± 0.02 | 0.70 ± 0.01 |
| | 1 week | 0.69 ± 0.03 | 0.82 ± 0.03 | 0.66 ± 0.02 | 0.82 ± 0.05$^{\dagger}$ |
| | 2 weeks | 0.70 ± 0.04 | 0.82 ± 0.03 | 0.71 ± 0.03 | 0.82 ± 0.05 |
| | 3 weeks | 0.69 ± 0.02 | 0.79 ± 0.03 | 0.71 ±0.01 | 0.87 ± 0.03$^{\dagger}$ |
| **LVPWs (mm)** | baseline | 1.04 ± 0.06 | 1.01 ± 0.05 | 1.07 ± 0.03 | 1.07 ± 0.04 |
| | 1 week | 1.06 ± 0.05 | 1.12 ± 0.05 | 0.99 ± 0.02 | 1.08 ± 0.06 |
| | 2 weeks | 1.05 ± 0.06 | 1.10 ± 0.05 | 1.02 ± 0.03 | 1.06 ± 0.05 |
| | 3 weeks | 1.06 ± 0.03 | 1.03 ± 0.04 | 0.99 ± 0.02 | 0.99 ± 0.04 |

All parameters are expressed as means ± SEM. Statistical analyses were performed using one-way ANOVA followed by Bonferroni post-hoc. *$p<0.05$ vs. respective sham group and $^{\dagger}p<0.05$ vs. cWT TAC. CO, Cardiac Output; EDV, Ejection Diastolic Volume; EF, Ejection Fraction; ESV, Ejection Systolic Volume; FS, Fractional Shortening; HR, Heart Rate; IVSd, Interventricular Septum Thickness at end Diastole, IVSs, Interventricular Septum Thickness at end Systole; LVIDd, Left-Ventricular Internal Diameter in diastole; LVIDs, Left-Ventricular Internal Diameter in systole; LVM, Left Ventricular mass; LVPWd, Left Ventricular Posterior Wall Thickness in diastole; LVPWs, Left Ventricular Posterior Wall Thickness in systole.

**Supplementary Table S5: Echocardiographic parameters of cWT and cKO females at baseline, 3 weeks and 6 weeks post-surgery.**

| | | cWT | | cKO | |
|---|---|---|---|---|---|
| | **Time** | **Sham** | **TAC** | **Sham** | **TAC** |
| **LVM (mg)** | baseline | 93.69 ± 0.68 | 85.66 ± 3.31 | 85.36 ± 2.37 | 95.12 ± 4.81 |
| | 3 weeks | 88.88 ± 2.21 | 110.25 ± 6.77* | 100.93 ± 5.02 | 137.60 ± 11.07*$^{\dagger}$ |
| | 6 weeks | 96.03 ± 1.03 | 123.58 ± 9.19* | 103.18 ± 4.02 | 155.22 ± 18.14*$^{\dagger}$ |
| **EDV (µL)** | baseline | 44.53 ± 4.32 | 39.60 ± 1.85 | 35.79 ± 2.45 | 45.70 ± 3.56* |
| | 3 weeks | 43.62 ± 3.92 | 48.25 ± 2.24 | 47.09 ± 2.56 | 72.20 ± 11.92*$^{\dagger}$ |
| | 6 weeks | 44.32 ± 5.41 | 58.85 ± 11.96 | 41.81 ± 4.08 | 93.14 ± 19.80*$^{\dagger}$ |
| **ESV (µL)** | baseline | 16.48 ± 1.78 | 15.15 ± 1.45 | 12.78 ± 1.26 | 17.36 ± 1.39* |
| | 3 weeks | 16.76 ± 1.30 | 23.87 ± 2.82 | 21.35 ± 2.42 | 51.80 ± 13.27*$^{\dagger}$ |
| | 6 weeks | 17.52 ± 2.50 | 38.60 ± 11.29 | 24.68 ± 4.44 | 79.78 ± 20.81*$^{\dagger}$ |
| **EF (%)** | baseline | 63.17 ± 0.68 | 61.94 ± 2.80 | 64.58 ± 1.16 | 61.98 ± 1.51 |
| | 3 weeks | 61.22 ± 2.03 | 50.49 ± 5.28 | 55.29 ± 2.82 | 33.11 ± 7.83*$^{\dagger}$ |
| | 6 weeks | 60.70 ± 1.03 | 38.97 ± 6.17* | 50.80 ± 4.61 | 19.03 ± 5.81*$^{\dagger}$ |
| **HR (bpm)** | baseline | 517.92 ± 19.28 | 562.63 ± 6.64 | 568.19 ± 23.95 | 535.32 ± 18.24 |
| | 3 weeks | 530.11 ± 13.14 | 519.98 ± 9.62 | 547.46 ± 14.56 | 520.18 ± 14.85 |
| | 6 weeks | 583.86 ± 19.04 | 555.61 ± 23.14 | 577.86 ± 18.67 | 562.93 ± 20.90 |
| **CO (mL/min)** | baseline | 14.98 ± 1.12 | 12.25 ± 1.19 | 13 ± 0.57 | 15.18 ± 1.23 |
| | 3 weeks | 14.21 ± 1.67 | 12.63 ± 1.35 | 14.11 ± 0.63 | 10.56 ± 1.62* |
| | 6 weeks | 15.58 ± 1.71 | 11.25 ± 1.54 | 13.32 ± 1.61 | 7.53 ± 1.61* |
| **FS (%)** | baseline | 35.69 ± 1.39 | 30.36 ± 3.61 | 37.04 ± 2.15 | 31.74 ± 2.13 |
| | 3 weeks | 35.48 ± 2.70 | 28.87 ± 1.52 | 31.37 ± 1.53 | 18.05 ± 4.36*$^{\dagger}$ |
| | 6 weeks | 36.94 ± 2.28 | 25.10 ± 4.84* | 28.79 ± 2.56 | 12.75 ± 4.80*$^{\dagger}$ |
| **IVSd (mm)** | baseline | 0.65 ± 0.04 | 0.78 ± 0.08 | 0.67 ± 0.01 | 0.74 ± 0.03 |
| | 3 weeks | 0.72 ± 0.02 | 0.85 ± 0.03* | 0.69 ± 0.02 | 0.81 ± 0.05* |
| | 6 weeks | 0.73 ± 0.04 | 0.90 ± 0.02* | 0.75 ± 0.02 | 0.81 ± 0.03$^{\dagger}$ |
| **IVSs (mm)** | baseline | 1.01 ± 0.04 | 1.07 ± 0.02 | 1.04 ± 0.03 | 1.10 ± 0.07 |
| | 3 weeks | 1.10 ± 0.05 | 1.17 ± 0.05 | 1.04 ± 0.04 | 1.06 ± 0.07 |
| | 6 weeks | 1.18 ± 0.06 | 1.24 ± 0.06 | 1.05 ± 0.04 | 1 ± 0.07$^{\dagger}$ |
| **LVIDd (mm)** | baseline | 3.35 ± 0.20 | 3.20 ± 0.13 | 3.12 ± 0.12 | 3.46 ± 0.09 |
| | 3 weeks | 3.32 ± 0.13 | 3.39 ± 0.10 | 3.46 ± 0.06 | 3.99 ± 0.33*$^{\dagger}$ |
| | 6 weeks | 3.28 ± 0.16 | 3.69 ± 0.36 | 3.40 ± 0.10 | 4.54 ± 0.43*$^{\dagger}$ |
| **LVIDs (mm)** | baseline | 2.16 ± 0.30 | 2.23 ± 0.15 | 1.96 ± 0.10 | 2.37 ± 0.12 |
| | 3 weeks | 2.14 ± 0.14 | 2.42 ± 0.10 | 2.38 ± 0.04 | 3.33 ± 0.43*$^{\dagger}$ |
| | 6 weeks | 2.08 ± 0.15 | 2.84 ± 0.44 | 2.43 ± 0.14 | 4.05 ± 0.57*$^{\dagger}$ |

| | | | | | |
|---|---|---|---|---|---|
| **LVPWd (mm)** | baseline | 0.63 ± 0.04 | 0.69 ± 0.03 | 0.67 ± 0.03 | 0.67 ± 0.03 |
| | 3 weeks | 0.67 ± 0.03 | 0.82 ± 0.06* | 0.66 ± 0.04 | 0.73 ± 0.04 |
| | 6 weeks | 0.67 ± 0.03 | 0.80 ± 0.04* | 0.67 ± 0.03 | 0.77 ± 0.04 |
| **LVPWs (mm)** | baseline | 0.91 ± 0.05 | 0.91 ± 0.03 | 1.00 ± 0.04 | 0.98 ± 0.05 |
| | 3 weeks | 0.98 ± 0.05 | 1.08 ± 0.09 | 0.93 ± 0.04 | 0.90 ± 0.08$^{\dagger}$ |
| | 6 weeks | 1.02 ± 0.05 | 0.99 ± 0.09 | 0.99 ± 0.05 | 0.90 ± 0.07 |

All parameters are expressed as means ± SEM. Statistical analyses were performed using one-way ANOVA followed by Bonferroni post-hoc. $^{*}p<0.05$ vs. respective sham group and $^{\dagger}p<0.05$ vs. cWT TAC. CO, Cardiac Output; EDV, Ejection Diastolic Volume; EF, Ejection Fraction; ESV, Ejection Systolic Volume; FS, Fractional Shortening; HR, Heart Rate; IVSd, Interventricular Septum Thickness at end Diastole, IVSs, Interventricular Septum Thickness at end Systole; LVIDd, Left-Ventricular Internal Diameter in diastole; LVIDs, Left-Ventricular Internal Diameter in systole; LVM, Left Ventricular mass; LVPWd, Left Ventricular Posterior Wall Thickness in diastole; LVPWs, Left Ventricular Posterior Wall Thickness in systole.

**Supplementary Table S6: Echocardiographic parameters of iWT and iKO males performed every 2 months.**

| | Time | iWT | iKO |
|---|---|---|---|
| **LVM (mg)** | baseline | 104.77 ± 3.84 | 99.47 ± 3.91 |
| | 2 months | 116.16 ± 3.70 | 126.15 ± 6.46 |
| | 4 months | 127.67 ± 3.69 | 141.46 ± 5.90 |
| | 6 months | 123.15 ± 3.59 | 139.22 ± 5.84* |
| | 8 months | 122.44 ± 4.71 | 140.79 ± 7.48* |
| | 10 months | 132.61 ± 2.68 | 155.69 ± 5.84* |
| | 12 months | 141.70 ± 7.93 | 171.0 ± 6.97* |
| **EDV (µL)** | baseline | 45.82 ± 4.14 | 46.38 ± 2.34 |
| | 2 months | 51.18 ± 1.86 | 58.61 ± 5.67 |
| | 4 months | 51.03 ± 2.01 | 56.98 ± 4.40 |
| | 6 months | 52.81 ± 2.45 | 53.82 ± 2.80 |
| | 8 months | 56.77 ± 2.14 | 56.48 ± 4.41 |
| | 10 months | 56.34 ± 2.15 | 58.84 ± 5.02 |
| | 12 months | 64.93 ± 3.36 | 63.96 ± 3.40 |
| **ESV (µL)** | baseline | 16.55 ± 1.87 | 20.70 ± 4.30 |
| | 2 months | 19.14 ± 0.76 | 29.34 ± 5.58 |
| | 4 months | 20.50 ± 1.31 | 29.77 ± 3.94* |
| | 6 months | 22.16 ± 1.15 | 31.49 ± 3.15* |
| | 8 months | 23.68 ± 0.73 | 32.34 ± 3.69* |
| | 10 months | 26.59 ± 1.02 | 36.75 ± 4.50 |
| | 12 months | 30.75 ± 1.92 | 37.12 ± 2.17 |
| **EF (%)** | baseline | 64.32 ± 1.27 | 63.62 ± 0.84 |
| | 2 months | 62.62 ± 0.71 | 52.75 ± 3.43* |
| | 4 months | 59.94 ± 1.69 | 49.27 ± 2.54* |
| | 6 months | 58.00 ± 1.10 | 42.89 ± 3.82* |
| | 8 months | 57.86 ± 1.97 | 44.08 ± 2.55* |
| | 10 months | 52.69 ± 1.16 | 38.79 ± 2.14* |
| | 12 months | 53.06 ± 1.92 | 41.73 ± 2.15* |
| **HR (bpm)** | baseline | 548.42 ± 1.14 | 521.33 ± 12.96 |
| | 2 months | 565.21 ± 8.00 | 548.22 ± 6.02 |
| | 4 months | 548.30 ± 12.16 | 550.18 ± 8.23 |
| | 6 months | 579.78 ± 16.83 | 556.28 ± 15.19 |
| | 8 months | 561.85 ± 16.35 | 561.09 ± 7.99 |
| | 10 months | 535.52 ± 12.40 | 530.95 ± 12.30 |
| | 12 months | 546.75 ± 14.69 | 519.19 ± 30.20 |
| **CO (mL/min)** | baseline | 16.18 ± 1.07 | 15.34 ± 0.80 |
| | 2 months | 18.46 ± 0.96 | 16.25 ± 0.64 |
| | 4 months | 17.27 ± 0.86 | 15.06 ± .037 |
| | 6 months | 17.90 ± 1.29 | 12.95 ± 1.00 |
| | 8 months | 18.66 ± 1.38 | 13.53 ± 0.79* |
| | 10 months | 15.91 ± 0.81 | 11.60 ± 0.54* |

| | | | |
|---|---|---|---|
| | 12 months | 15.45 ± 0.06 | 12.75 ± 0.58 |
| **FS (%)** | baseline | 35.67 ± 0.92 | 34.19 ± 0.80 |
| | 2 months | 34.04 ± 1.16 | 28.30 ± 2.05* |
| | 4 months | 36.98 ± 2.15 | 28.24 ± 2.60* |
| | 6 months | 32.07 ± 1.62 | 26.40 ± 2.25 |
| | 8 months | 37.66 ± 1.85 | 26.84 ± 2.20* |
| | 10 months | 32.78 ± 0.69 | 26.87 ± 1.24* |
| | 12 months | 33.35 ± 1.77 | 22.39 ± 1.88* |
| **IVSd (mm)** | baseline | 0.78 ± 0.01 | 0.81 ± 0.06 |
| | 2 months | 0.83 ± 0.03 | 0.81 ± 0.02 |
| | 4 months | 0.82 ± 0.03 | 0.82 ± 0.02 |
| | 6 months | 0.88 ± 0.06 | 0.88 ± 0.03 |
| | 8 months | 0.93 ± 0.03 | 0.89 ± 0.03 |
| | 10 months | 0.88 ± 0.02 | 0.95 ± 0.03 |
| | 12 months | 0.93 ± 0.03 | 0.98 ± 0.07 |
| **IVSs (mm)** | baseline | 1.14 ± 0.03 | 1.09 ± 0.03 |
| | 2 months | 1.17 ± 0.04 | 1.10 ± 0.04 |
| | 4 months | 1.16 ± 0.04 | 1.09 ± 0.03 |
| | 6 months | 1.18 ± 0.04 | 1.19 ± 0.06 |
| | 8 months | 1.35 ± 0.03 | 1.25 ± 0.04 |
| | 10 months | 1.33 ± 0.04 | 1.24 ± 0.02* |
| | 12 months | 1.33 ± 0.05 | 1.26 ± 0.07 |
| **LVIDd (mm)** | baseline | 3.33 ± 0.12 | 3.47 ± 0.09 |
| | 2 months | 3.57 ± 0.09 | 3.75 ± 0.16 |
| | 4 months | 3.50 ± 0.10 | 3.64 ± 0.13 |
| | 6 months | 3.46 ± 0.10 | 3.38 ± 0.13 |
| | 8 months | 3.37 ± 0.12 | 3.62 ± 0.12 |
| | 10 months | 3.74 ± 0.06 | 3.65 ± 0.14 |
| | 12 months | 3.73 ± 0.11 | 3.82 ± 0.14 |
| **LVIDs (mm)** | baseline | 2.15 ± 0.09 | 2.29 ± 0.08 |
| | 2 months | 2.35 ± 0.07 | 2.70 ± 0.18 |
| | 4 months | 2.22 ± 0.12 | 2.64 ± 0.18 |
| | 6 months | 2.36 ± 0.12 | 2.50 ± 0.15 |
| | 8 months | 2.37 ± 0.12 | 2.66 ± 0.13 |
| | 10 months | 2.52 ± 0.06 | 2.67 ± 0.13 |
| | 12 months | 2.50 ± 0.13 | 2.97 ± 0.13* |
| **LVPWd (mm)** | baseline | 0.68 ± 0.02 | 0.68 ± 0.03 |
| | 2 months | 0.78 ± 0.02 | 0.77 ± 0.02 |
| | 4 months | 0.80 ± 0.02 | 0.76 ± 0.02 |
| | 6 months | 0.81 ± 0.02 | 0.83 ± 0.03 |
| | 8 months | 0.89 ± 0.04 | 0.84 ± 0.03 |
| | 10 months | 0.86 ± 0.03 | 0.80 ± 0.04 |
| | 12 months | 0.87 ± 0.04 | 0.81 ± 0.03* |
| **LVPWs (mm)** | baseline | 0.99 ± 0.03 | 1.02 ± 0.03 |
| | 2 months | 1.08 ± 0.03 | 1.06 ± 0.05 |
| | 4 months | 1.12 ± 0.04 | 1.06 ± 0.05 |

| | | |
|---|---|---|
| 6 months | 1.11 ± 0.03 | 1.17 ± 0.06 |
| 8 months | 1.27 ± 0.05 | 1.12 ± 0.04* |
| 10 months | 1.24 ± 0.04 | 1.13 ± 0.04 |
| 12 months | 1.29 ± 0.06 | 1.03 ± 0.03* |

All parameters are expressed as means ± SEM. Statistical analyses were performed using t-test followed Bonferroni. *$p<0.05$ vs. iWT mice. CO, Cardiac Output; EDV, Ejection Diastolic Volume; EF, Ejection Fraction; ESV, Ejection Systolic Volume; FS, Fractional Shortening; HR, Heart Rate; IVSd, Interventricular Septum Thickness at end Diastole, IVSs, Interventricular Septum Thickness at end Systole; LVIDd, Left-Ventricular Internal Diameter in diastole; LVIDs, Left-Ventricular Internal Diameter in systole; LVM, Left Ventricular mass; LVPWd, Left Ventricular Posterior Wall Thickness in diastole; LVPWs, Left Ventricular Posterior Wall Thickness in systole.

**Supplementary Table S7: Echocardiographic parameters of iWT and iKO males at baseline, 3- , 6- and 9-weeks post-surgery.**

| | | iWT | | iKO | |
|---|---|---|---|---|---|
| | **Time** | **Sham** | **TAC** | **Sham** | **TAC** |
| **LVM (mg)** | baseline | 112.80 ± 4.80 | 103.73 ± 3.58 | 103.05 ± 2.93 | 103.89 ± 4.51 |
| | 3 weeks | 124.75 ± 7.44 | 142.34 ± 5.31 | 127.08 ± 3.54 | 170.85 ± 5.07*† |
| | 6 weeks | 123.25 ± 6.89 | 172.61 ± 12.09* | 130.54 ± 5.36 | 220.89 ± 14.76*† |
| | 9 weeks | 115.86 ± 5.68 | 174.57 ± 12.63* | 130.89 ± 11.98 | 223.71 ± 15.39*† |
| **EDV (µL)** | baseline | 53.82 ± 1.65 | 51.08 ± 6.31 | 51.77 ± 3.16 | 51.07 ± 3.33 |
| | 3 weeks | 56.28 ± 4.36 | 71.19 ± 6.11 | 67.50 ± 3.90 | 78.57 ± 4.38 |
| | 6 weeks | 51.31 ± 5.31 | 75.94 ± 8.23 | 57.14 ± 5.85 | 98.97 ± 7.09* |
| | 9 weeks | 64.94 ± 3.94 | 94.12 ± 11.55 | 62.04 ± 4.56 | 115.42 ± 9.94* |
| **ESV (µL)** | baseline | 20.06 ± 0.76 | 19.67 ± 0.90 | 18.93 ± 1.86 | 18.97 ± 1.49 |
| | 3 weeks | 21.94 ± 1.29 | 39.32 ± 4.56* | 27.56 ± 2.38 | 56.89 ± 4.31*† |
| | 6 weeks | 24.05 ± 3.92 | 48.43 ± 7.97* | 22.22 ± 5.60 | 77.15 ± 7.30*† |
| | 9 weeks | 23.93 ± 1.45 | 65.54 ± 12.47* | 27.86 ± 3.80 | 98.34 ± 10.22*† |
| **EF (%)** | baseline | 62.63 ± 1.08 | 61.55 ± 0.55 | 63.78 ± 2.05 | 63.11 ± 0.81 |
| | 3 weeks | 62.50 ± 0.88 | 45.90 ± 2.06* | 59.43 ± 1.78 | 20.08 ± 2.17*† |
| | 6 weeks | 62.74 ± 0.88 | 38.87 ± 3.60* | 61.34 ± 1.15 | 23.48 ± 2.66*† |
| | 9 weeks | 63.29 ± 0.66 | 34.60 ± 4.59* | 55.83 ± 3.78 | 15.98 ± 2.31*† |
| **HR (bpm)** | baseline | 494.59 ± 6.90 | 515.48 ± 12.67 | 518.11 ± 9.52 | 529.57 ± 16.38 |
| | 3 weeks | 536.77 ± 12.22 | 514.92 ± 12.25 | 508.55 ± 12.93 | 529.04 ± 18.19 |
| | 6 weeks | 551.30 ± 9.84 | 542.80 ± 12.35 | 535.00 ± 18.51 | 526.67 ± 14.85 |
| | 9 weeks | 532.79 ± 5.56 | 530.20 ± 18.99 | 549.80 ± 18.18 | 526.64 ± 10.62 |
| **CO (mL/min)** | baseline | 16.67 ± 0.56 | 16.27 ± 0.93 | 17.05 ± 1.04 | 16.85 ± 0.90 |
| | 3 weeks | 19.48 ± 0.84 | 16.41 ± 0.93 | 20.20 ± 1.02 | 11.4 ± 0.99*† |
| | 6 weeks | 17.78 ± 1.16 | 15.67 ± 1.22 | 18.45 ± 1.42 | 11.57 ± 1.26* |
| | 9 weeks | 18.49 ± 1.17 | 15.05 ± 1.13 | 18.70 ± 1.22 | 8.99 ± 0.98*† |
| **FS (%)** | baseline | 33.38 ± 1.89 | 31.05 ± 1.97 | 38.04 ± 2.57 | 30.26 ± 0.98 |
| | 3 weeks | 32.84 ± 1.46 | 23.11 ± 2.03* | 31.80 ± 2.43 | 15.54 ± 1.47*† |
| | 6 weeks | 34.57 ± 1.32 | 21.68 ± 2.79* | 31.92 ± 2.74 | 9.89 ± 1.97*† |
| | 9 weeks | 35.72 ± 2.92 | 18.59 ± 3.19* | 32.78 ± 2.13 | 7.02 ± 1.19*† |
| **IVSd (mm)** | baseline | 0.81 ± 0.02 | 0.71 ± 0.02 | 0.75 ± 0.03 | 0.76 ± 0.03 |
| | 3 weeks | 0.83 ± 0.06 | 0.91 ± 0.05 | 0.82 ± 0.03 | 0.98 ± 0.03 |
| | 6 weeks | 0.90 ± 0.03 | 0.95 ± 0.03 | 0.87 ± 0.04 | 0.98 ± 0.06 |
| | 9 weeks | 0.78 ± 0.03 | 0.84 ± 0.03 | 0.88 ± 0.03 | 0.94 ± 0.05 |
| **IVSs (mm)** | baseline | 1.25 ± 0.06 | 1.09 ± 0.03 | 1.18 ± 0.06 | 1.14 ± 0.03 |
| | 3 weeks | 1.20 ± 0.10 | 1.27 ± 0.07 | 1.17 ± 0.07 | 1.24 ± 0.04 |
| | 6 weeks | 1.32 ± 0.05 | 1.24 ± 0.05 | 1.21 ± 0.07 | 1.13 ± 0.06 |
| | 9 weeks | 1.24 ± 0.04 | 1.14 ± 0.08 | 1.29 ± 0.04 | 1.08 ± 0.04 |

| | | | | | |
|---|---|---|---|---|---|
| **LVIDd (mm)** | baseline | 3.40 ± 0.12 | 3.54 ± 0.05 | 3.52 ± 0.15 | 3.62 ± 0.11 |
| | 3 weeks | 3.56 ± 0.11 | 3.70 ± 0.10 | 3.67 ± 0.10 | 4.05 ± 0.16 |
| | 6 weeks | 3.46 ± 0.07 | 3.75 ± 0.20 | 3.54 ± 0.16 | 4.32 ± 0.21* |
| | 9 weeks | 3.78 ± 0.16 | 4.01 ± 0.20 | 3.75 ± 0.19 | 4.69 ± 0.24* |
| **LVIDs (mm)** | baseline | 2.28 ± 0.11 | 2.44 ± 0.06 | 2.20 ± 0.17 | 2.53 ± 0.09 |
| | 3 weeks | 2.40 ± 0.11 | 2.86 ± 0.13 | 2.51 ± 0.13 | 3.43 ± 0.17*† |
| | 6 weeks | 2.27 ± 0.07 | 2.98 ± 0.25 | 2.42 ± 0.18 | 3.93 ± 0.26*† |
| | 9 weeks | 2.45 ± 0.19 | 3.30 ± 0.28 | 2.54 ± 0.19 | 4.38 ± 0.26*† |
| **LVPWd (mm)** | baseline | 0.73 ± 0.04 | 0.67 ± 0.04 | 0.71 ± 0.02 | 0.71 ± 0.02 |
| | 3 weeks | 0.78 ± 0.02 | 0.87 ± 0.05 | 0.82 ± 0.04 | 0.93 ± 0.04 |
| | 6 weeks | 0.83 ± 0.04 | 0.90 ± 0.03 | 0.81 ± 0.03 | 0.93 ± 0.03 |
| | 9 weeks | 0.82 ± 0.04 | 0.96 ± 0.04 | 0.87 ± 0.02 | 0.98 ± 0.06 |
| **LVPWs (mm)** | baseline | 1.10 ± 0.04 | 1.00 ± 0.05 | 1.04 ± 0.05 | 1.01 ± 0.03 |
| | 3 weeks | 1.14 ± 0.04 | 1.15 ± 0.18 | 1.08 ± 0.40 | 1.10 ± 0.05 |
| | 6 weeks | 1.22 ± 0.06 | 1.16 ± 0.03 | 1.13 ± 0.07 | 1.05 ± 0.06 |
| | 9 weeks | 1.20 ± 0.06 | 1.16 ± 0.05 | 1.22 ± 0.05 | 1.07 ± 0.07 |

All parameters are expressed as means ± SEM. Statistical analyses were performed using one-way ANOVA followed by Bonferroni post-hoc. *$p<0.05$ vs. respective sham group and †$p<0.05$ vs. iWT TAC. CO, Cardiac Output; EDV, Ejection Diastolic Volume; EF, Ejection Fraction; ESV, Ejection Systolic Volume; FS, Fractional Shortening; HR, Heart Rate; IVSd, Interventricular Septum Thickness at end Diastole, IVSs, Interventricular Septum Thickness at end Systole; LVIDd, Left-Ventricular Internal Diameter in diastole; LVIDs, Left-Ventricular Internal Diameter in systole; LVM, Left Ventricular mass; LVPWd, Left Ventricular Posterior Wall Thickness in diastole; LVPWs, Left Ventricular Posterior Wall Thickness in systole.

**Supplementary Table 8: List of antibodies used for western-blotting and immunostaining in this work.**

| PRIMARY ANTIBODY | COMPANY | REFERENCE # | WB | IF/IHC |
|---|---|---|---|---|
| | | | WORKING CONCENTRATION | |
| skNAC | Homemade | - | 1/500 | 1/500 |
| αNAC | Invitrogen | 40-1000 | 1/500 | - |
| LC3 | Cell signaling | 2775 | 1/500 | |
| P62 | Cell signaling | 23214 | 1/1000 | |
| O-linked N acetylglucosamine-HRP | Abcam | Ab201995 | 1/1000 | - |
| ERK1/2 | Cell signaling | 9102 | 1/1000 | - |
| p-ERK1/2 ($Thr^{202}/Tyr^{204}$) | Cell signaling | 9101 | 1/1000 | - |
| S6 ribosomal protein | Cell signaling | 2217 | 1/1000 | - |
| p-S6 ($S^{235/236}$) | Cell signaling | 4856 | 1/1000 | - |
| eEF2 | Invitrogen | PA5-17794 | 1/1000 | - |
| GAPDH (14C10) | Cell signaling | 2118S | 1/50000 | - |
| Alpha-actinin | Sigma | A7811 | - | 1/1000 |
| S6 ribosomal protein | Cell signaling | 2317 | - | 1/200 |
| KDEL | Enzo | ADI-SPA-827-D | - | 1/200 |
| TTN-Z | Custom made with Myomedix | - | - | 1/100 |

| SECONDARY ANTIBODY | COMPANY | REFERENCE # | WB | IF/IHC |
|---|---|---|---|---|
| | | | WORKING CONCENTRATION | |
| Anti-rabbit IgG-HRP | Sigma | A0545 | 1/20,000 | - |
| Anti-mouse IgG-HRP | BD Bioscience | 554002 | 1/5000 | - |
| Alexa Fluor 594 donkey anti-mouse | Invitrogen | A2103 | | 1/1000 |
| Alexa Fluor 488 donkey anti-mouse | Invitrogen | A21206 | | 1/1000 |
| Cy3 Goat anti-rabbit IgG | Jackson ImmunoResearch | 111-165-003 | | 1/400 |
| Alexa 488 Goat anti-mouse IgG | Jackson ImmunoResearch | 115-546-146 | | 1/400 |
| Alexa 647 Goat anti-rabbit IgG | Jackson ImmunoResearch | 111-606-047 | | 1/400 |

**Supplementary Table 9: List of primers and siRNA homemade used in this work.**

| | ORGANISMS | GENES | SEQUENCES 5' → 3' |
|---|---|---|---|
| qPCR | Rattus Norvegicus | ***skNAC*** | CTGCTACAGAGCAGGAGTTGC |
| | | | CAGCAGTCACTAGAGGGCTTT |
| | | ***αNAC*** | CTGCTACAGAGCAGGAGTTGC |
| | | | TGTGGCTGTCTGTGTGGAATC |
| | | ***Nppa* (ANP)** | TGATGGATTTCAAGAACCTGCT |
| | | | CTTCATCGGTCTGCTCGCTC |
| | | ***Nppb* (BNP)** | AACAATCCACGATGCAGAAGC |
| | | | GGCGCTGTCTTGAGACCTAA |
| | | ***Myh7* (β-MHC)** | CCGAGTCCCAGGTCAACAAG |
| | | | TACTCTTCATTCAGGCCCTTGG |
| | | ***Rbm24* (RNA-binding protein 24)** | CCCTTATCCAGAGACCTTTTG |
| | | | GCGGCTCCAGTGTAATCTAT |
| | | ***Rbm20* (RNA-bonding protein 20)** | GGCTTACACAGAAGCTGCTCAAG |
| | | | GTGGATGTCCTGGATGATAGCAG |
| | | ***Rpl32* (Ribosomal protein L32)** | CACAGCTGGCCATCAGAGTCA |
| | | | AAACAGGCACACAAGCCATCTATTC |
| | Mus Musculus | ***skNAC*** | TACAGAGCAGGAGTTGCCACAG |
| | | | AGTTACTATAGGGCTTTGCTGGG |
| | | ***αNAC*** | CAGAAACCGTCCCTGCTACA |
| | | | GTTCCTCGAGCTCTGGTACTG |
| | | ***Nppa* (ANP)** | CTGCTTCGGGGGTAGGATTG |
| | | | GCTCAAGCAGAATCGACTGC |
| | | ***Nppb* (BNP)** | CTTCGGTCTCAAGGCAGCA |
| | | | CAACTTCAGTGCGTTACAGCCC |
| | | ***Myh7* (β-MHC)** | GCATTCTCCTGCTGTTTCCT |
| | | | CCCAAATGCAGCCATCTC |
| | | ***Rbm24* (RNA-binding protein 24)** | TTGCCTTTGGCGTTCAACAG |
| | | | ATGTACGGTGTGGTGGAAGC |
| | | ***Rpl32* (Ribosomal protein L32)** | GGCACCAGTCAGACCGATAT |
| | | | CAGGATCTGGCCCTTGAAC |
| | Human | ***skNAC*** | TACAACTACAGCCAGCCCTTT |
| | | | GCCAGAGGAGCACAGGTATT |
| | | ***Rpl32* (Ribosomal protein L32)** | AGGCATTGACAACAGGGTTC |
| | | | GTTGCACATCAGCAGCACTT |
| PCR | Mus Musculus | **Genotyping Cre** | GCC TGC ATT ACC GGT CGA TGC AAC GA |
| | | | GTG GCA GAT GGC GCG GCA ACA CCA TT |
| | | **Genotyping *skNAC* (LoxP)** | TTCCAGCCGATCCTCGTCAA |
| | | | CTTACTTAGCCTGGCCCACTT |
| | | **Genotyping *skNAC* (LoxP)** | GGTGTGGGCCTTTGCTCTTC |
| | | | CAGCAGCAACTTTCAAGGCT |
| siRNA | Rattus Norvegicus | **skNAC** | GACAGUCCCUGUUGAGAAUU |

### Full unedited gel of Figures

Representative images shown in the Figures

**Figure 1a**

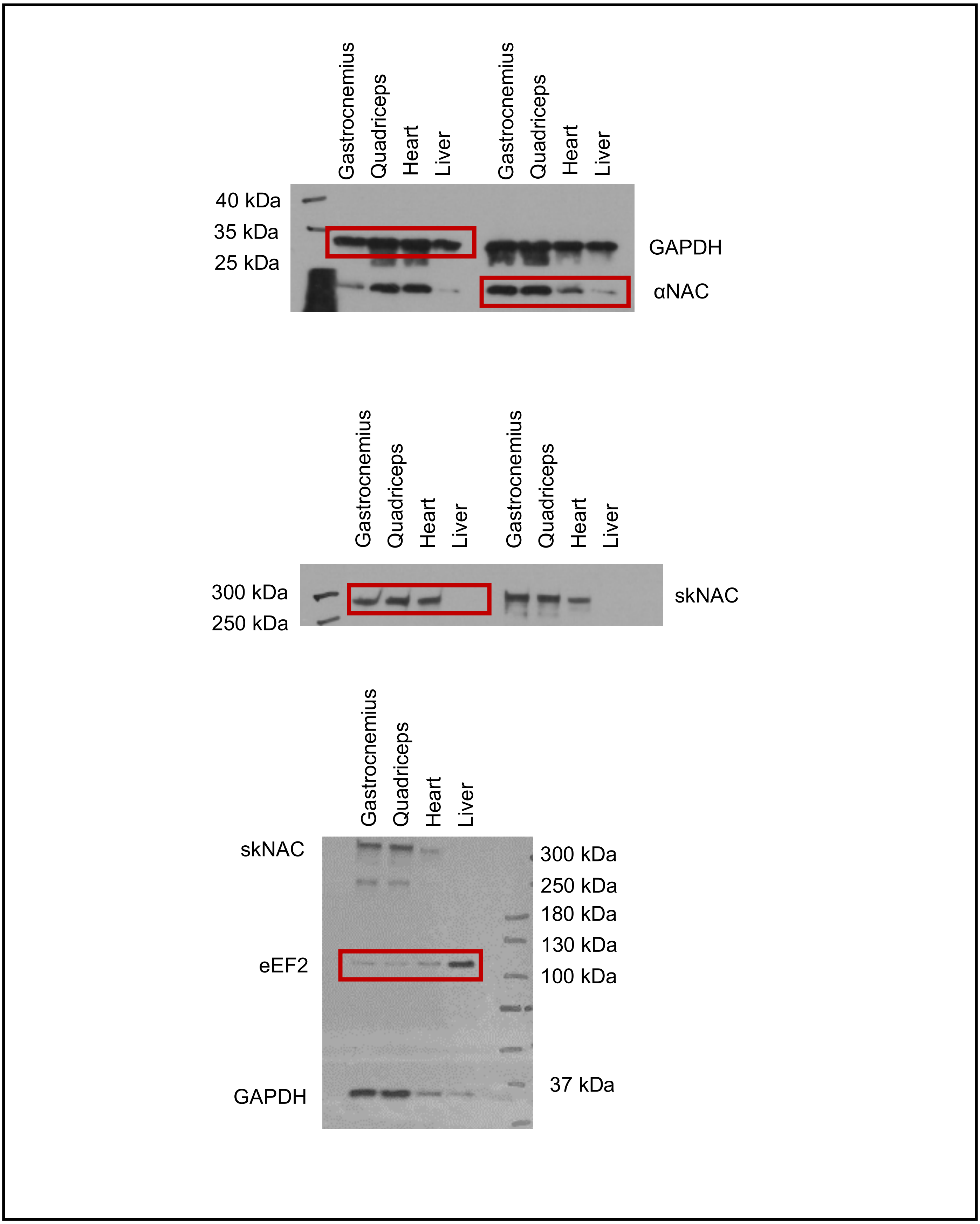

Gastrocnemius
Quadriceps
Heart
Liver
Gastrocnemius
Quadriceps
Heart
Liver
40 kDa
35 kDa
25 kDa
GAPDH
αNAC
Gastrocnemius
Quadriceps
Heart
Liver
Gastrocnemius
Quadriceps
Heart
Liver
300 kDa
250 kDa
skNAC
Gastrocnemius
Quadriceps
Heart
Liver
skNAC
eEF2
GAPDH
300 kDa
250 kDa
180 kDa
130 kDa
100 kDa
37 kDa

**Figure 3g**

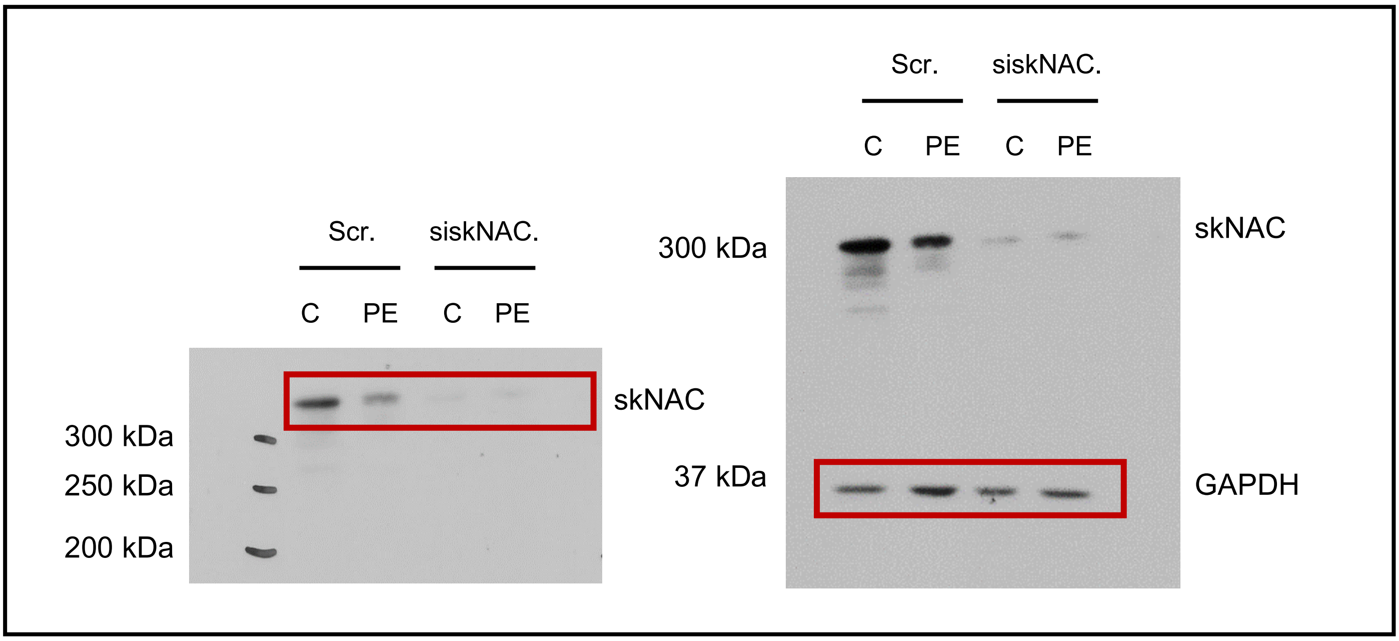

**Figure 4j,k**

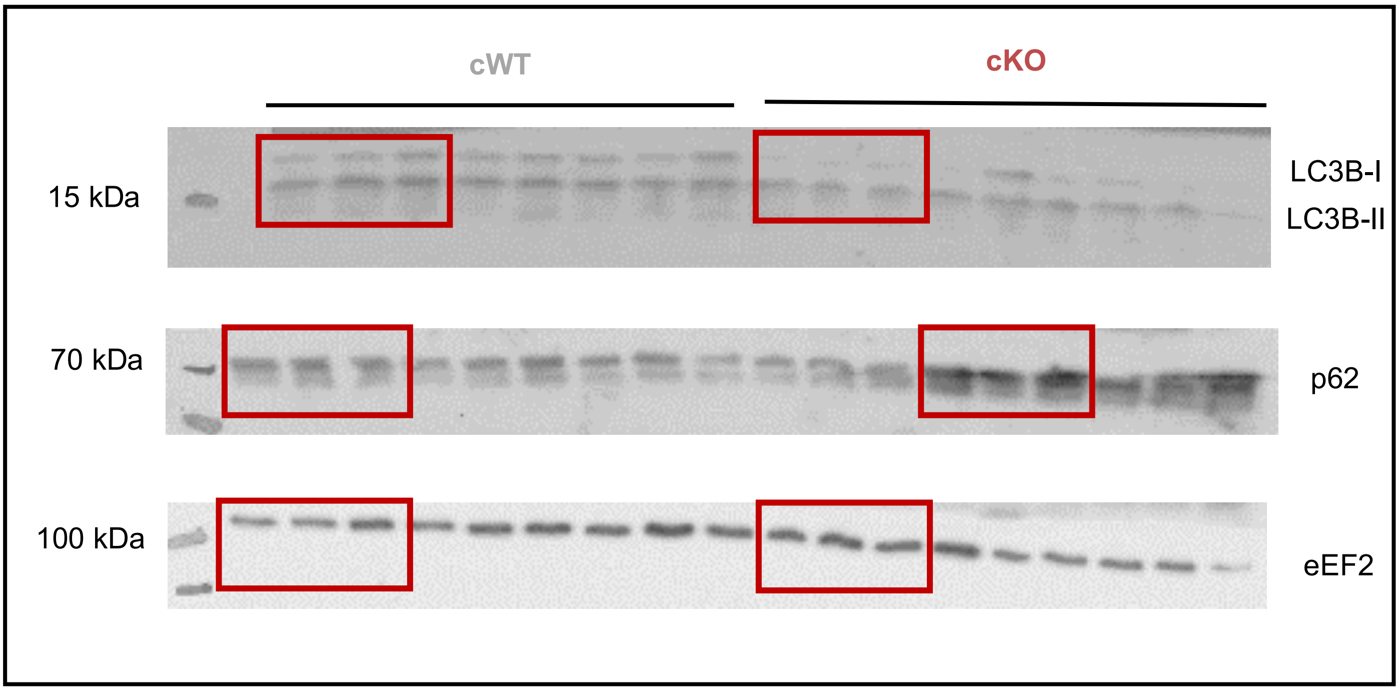

**Figure S6b**

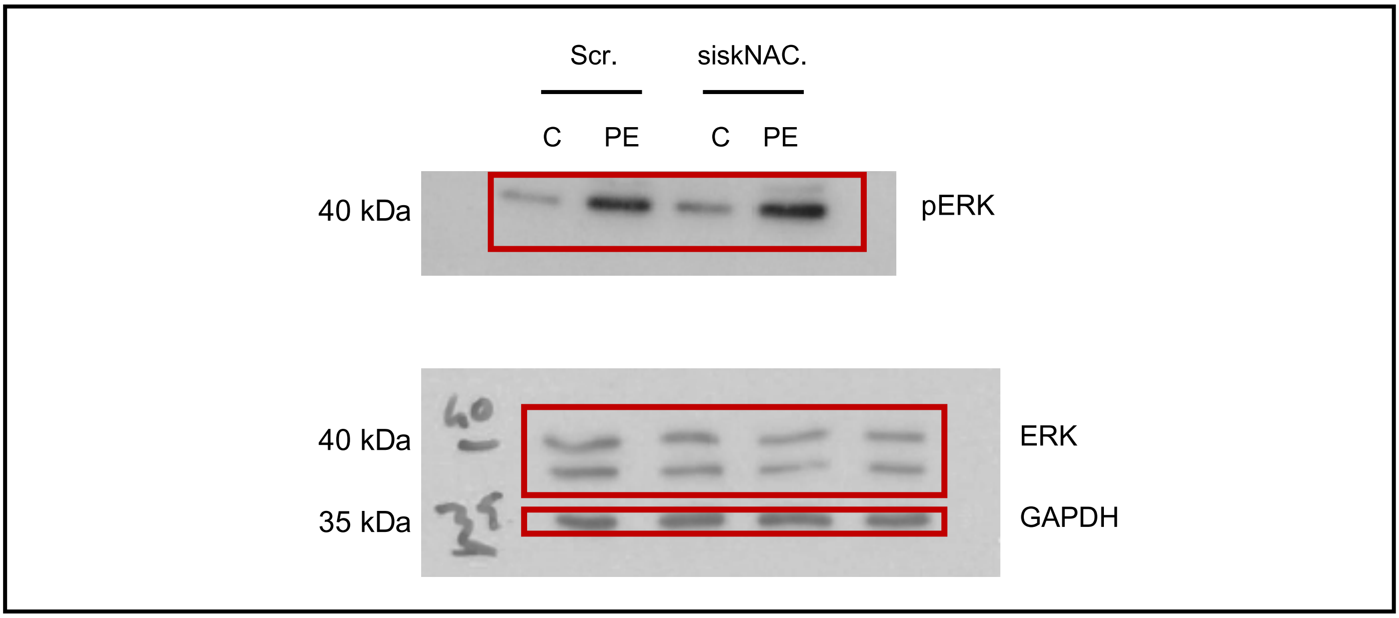

**Figure S6c**

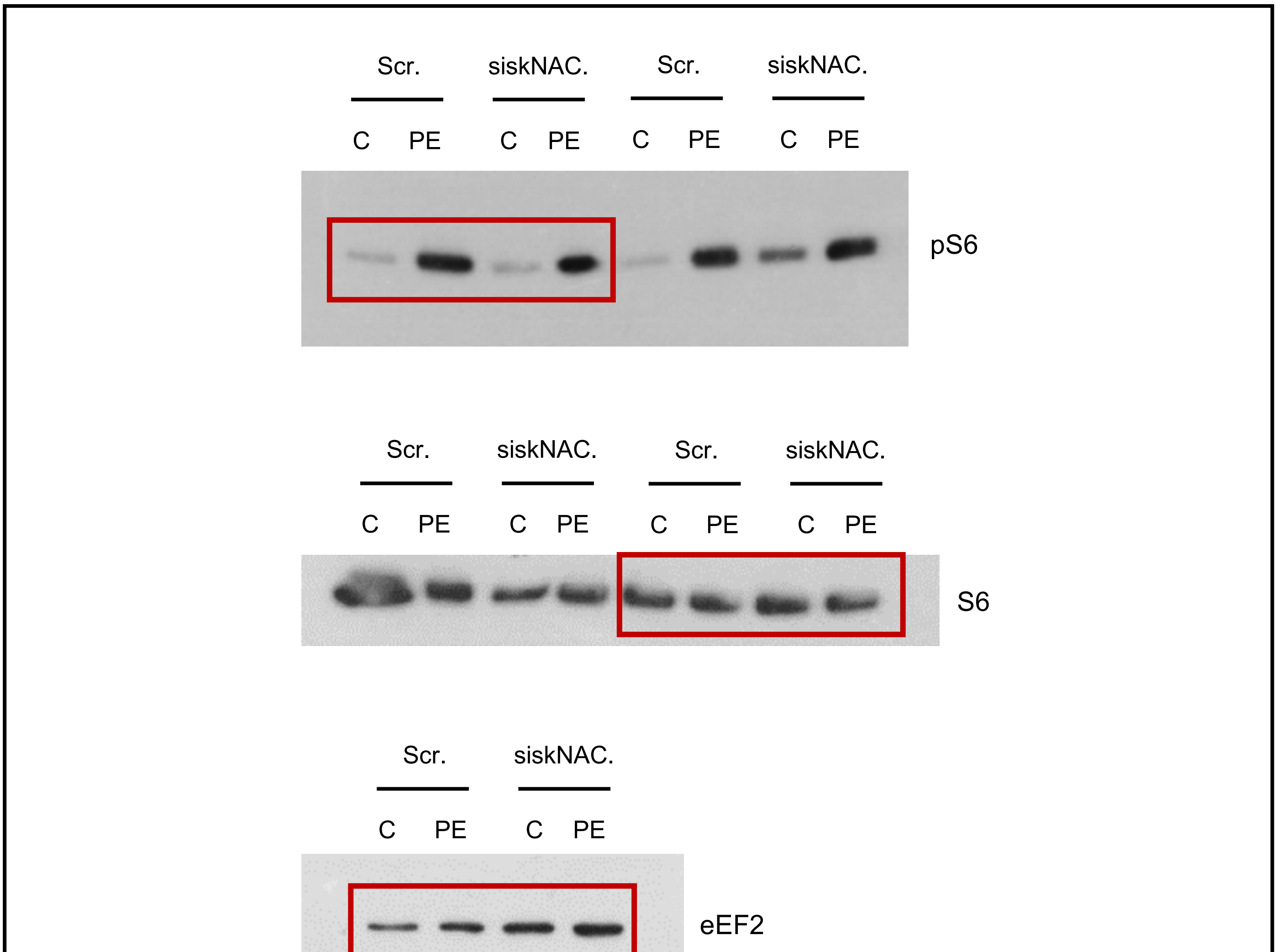

**Figure S7d**

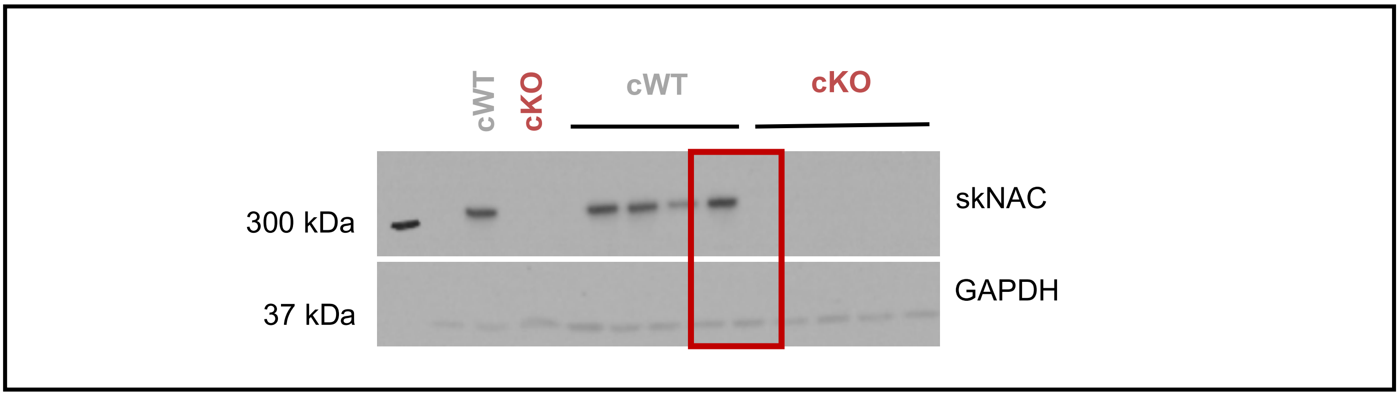

**Figure S7e**

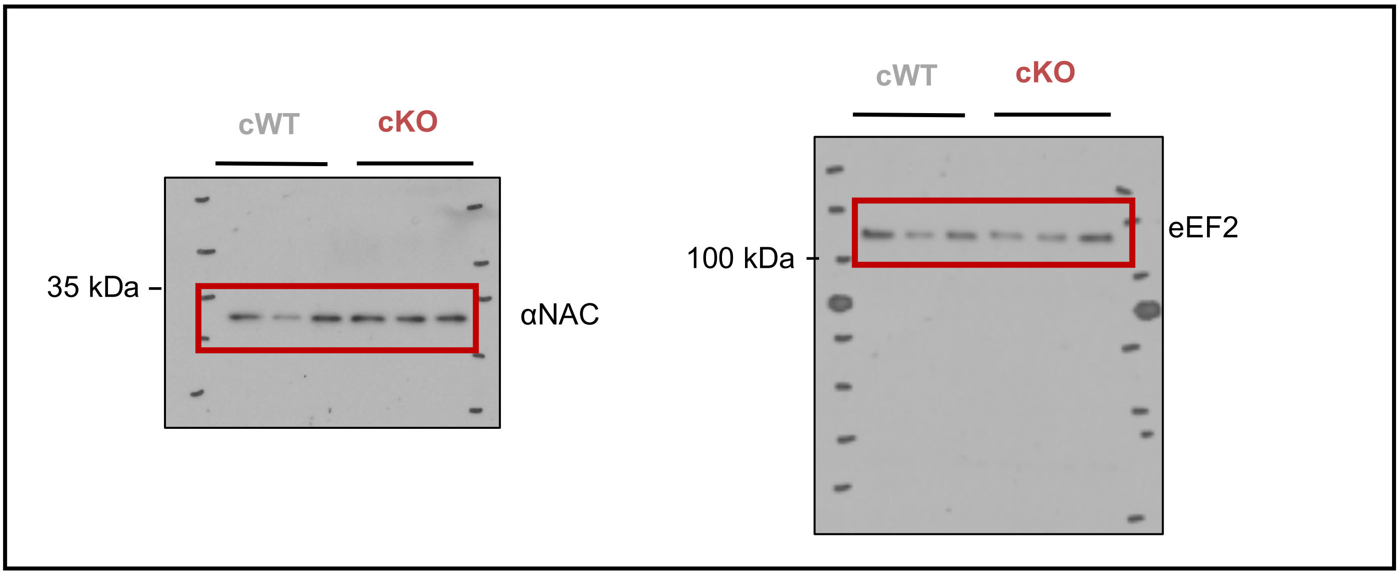

**Figure S7f**

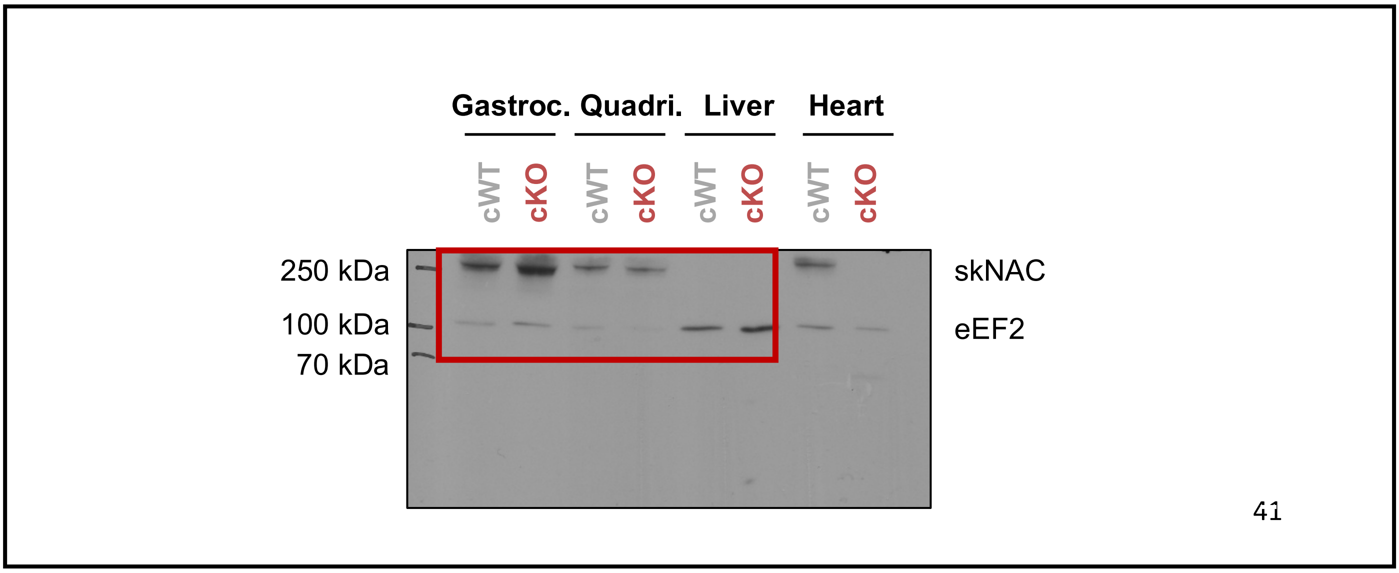

**Figure S9d**

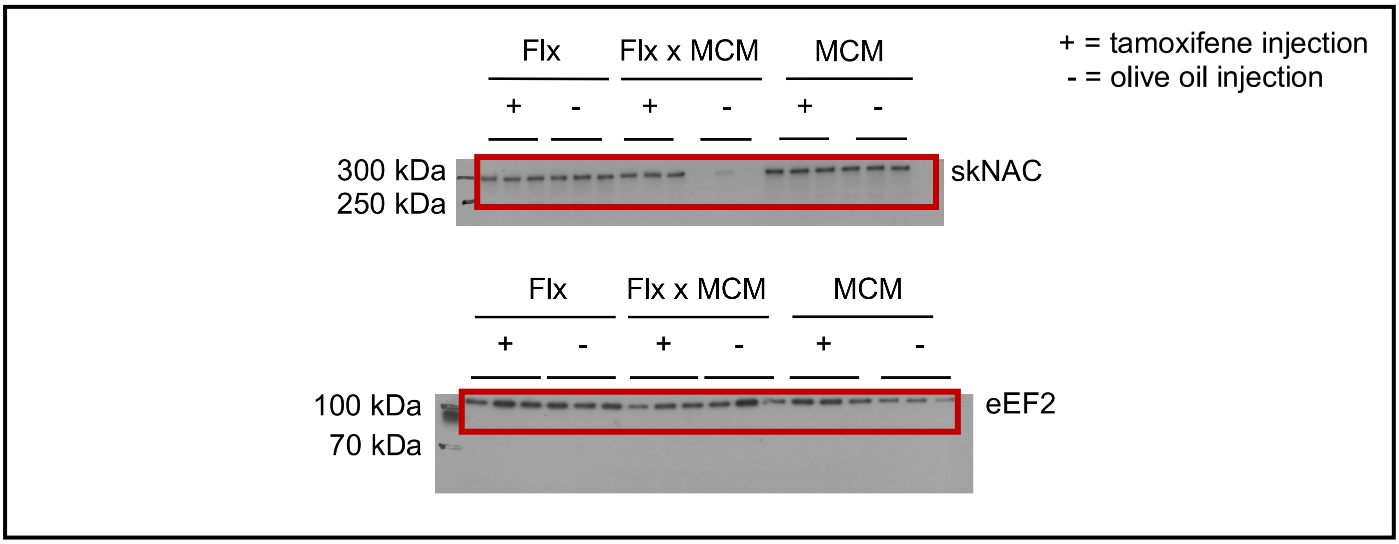

**Figure S9e**

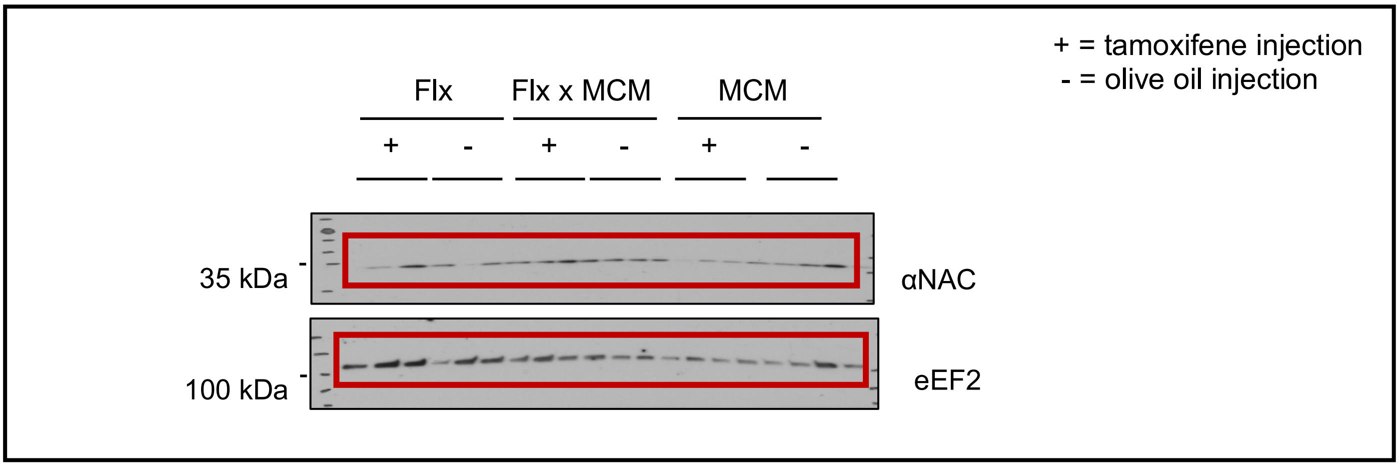

**Figure S9f**

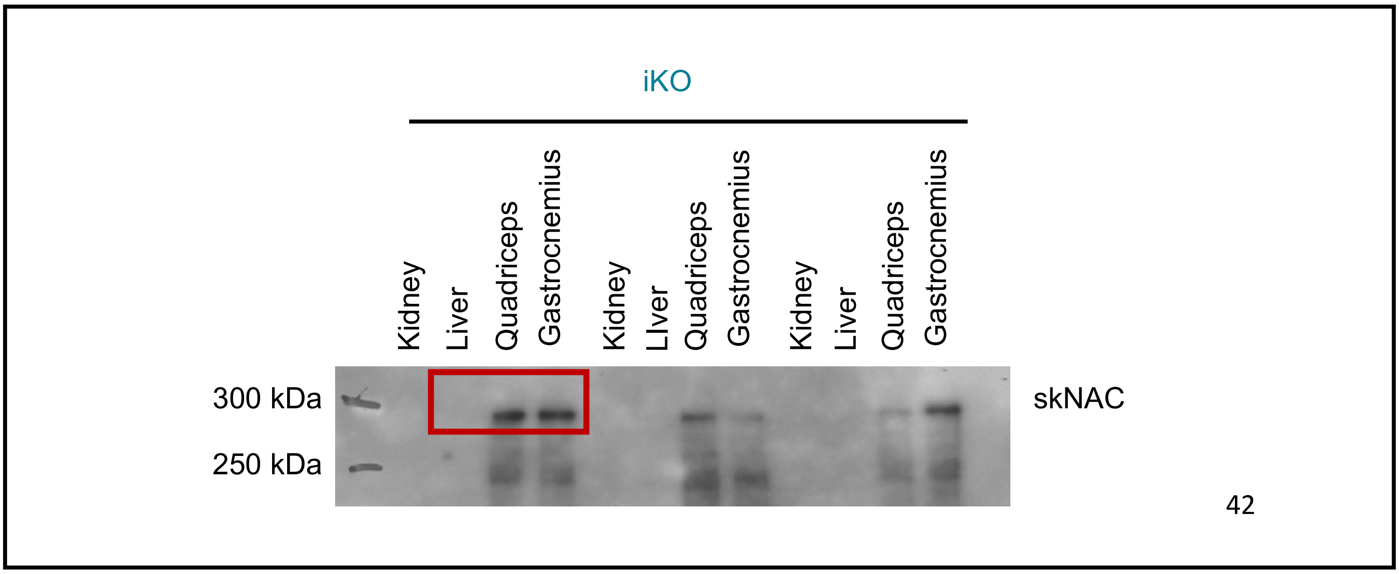

**Figure S15a,b**

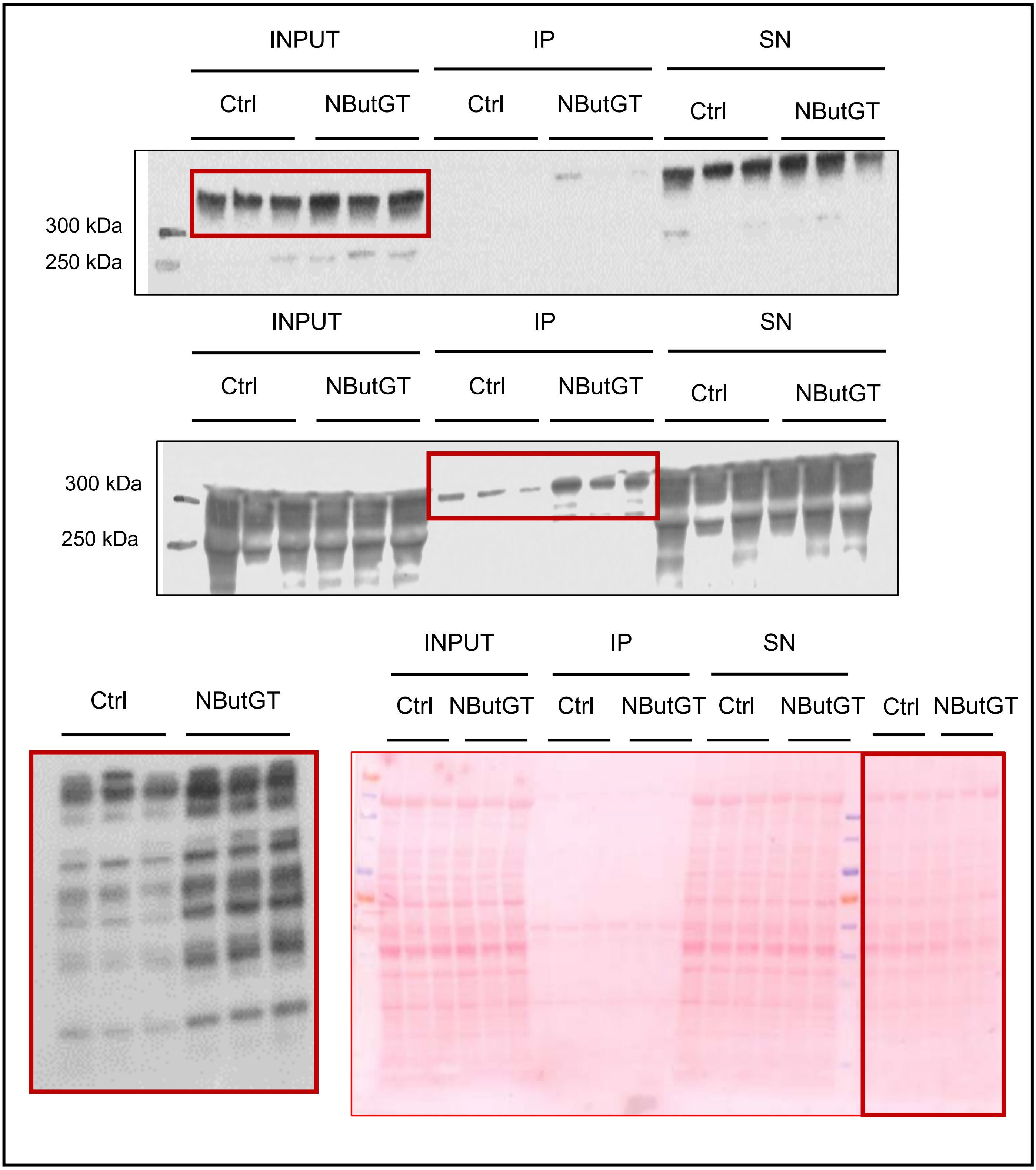
